# A Divergent Class of Arylamine N-Acetyltransferases Catalyzes Convergent Amidative Condensation of Polyketides in Manumycins Biosynthesis

**DOI:** 10.64898/2026.04.15.718676

**Authors:** Xiaoli Yan, Yan Gao, Bao-Di Ma, Qiaqia Zhou, Minghe Luo, Guangzheng Wei, Zhi Lin, Zixin Deng, Xu-Dong Kong, Xudong Qu

**Author notes:** Authors contributed equally (Xiaoli Yan, Yan Gao, Bao-Di Ma).

## Abstract

Polyketides are prized for their structural complexity and therapeutic potential, yet the incorporation of amide bonds into their frameworks typically relies on linear, nonribosomal peptide synthetase (NRPS)-dependent assembly. The direct, convergent coupling of distinct polyketide chains via amide bond formation—an ideal strategy for combinatorial biosynthesis—has remained largely elusive. Here, we report the discovery of a novel family of arylamine N-acetyltransferases (NATs) from manumycin-type biosynthetic pathways that catalyze an unprecedented intermolecular amidative chain transfer/condensation between an acyl carrier protein (ACP)-tethered polyketide donor and a free polyketide acceptor. Biochemical, structural, and molecular dynamics studies reveal that the representative enzyme, ColC2, possesses a distinctive substrate-binding pocket that diverges from canonical arylamine NATs, conferring exceptional promiscuity toward diverse acyl donors and acceptors. We demonstrate the utility of this biocatalyst by coupling arylamines with either synthetic acyl-thioesters or polyketide synthase (PKS) machinery to generate a library of non-natural polyketide amides and manumycin derivatives. These findings establish a new paradigm for amide bond formation in polyketide biosynthesis and position arylamine NATs as powerful tools for the development of novel therapeutics through combinatorial synthesis.

## Introduction

Polyketides represent a structurally diverse class of natural products with profound pharmacological significance^1^. Their carbon scaffolds are typically assembled by polyketide synthases (PKSs) via decarboxylative Claisen condensations of acyl- and malonyl-thioesters^2^. Amide bonds are critical motifs in drug discovery, offering metabolic stability, hydrogen-bonding sites, and structural rigidity that enhance target affinity^3^. In polyketides, these bonds are usually introduced by nonribosomal peptide synthetase (NRPS) modules, where condensation (C) domains incorporate amino acids, or facilitate cyclization via thioesterase (TE) domains^4,5^. While rare specialized mechanisms link polyketides to ribosomally synthesized and post-translationally modified peptides (RiPPs)^6^ or nucleosides^7^ (Fig. 1a), the direct convergent coupling of two independent polyketide chains via an amide bond has long been speculative, proposed primarily for the manumycin-type metabolites^8,9^.

**Fig. 1.**
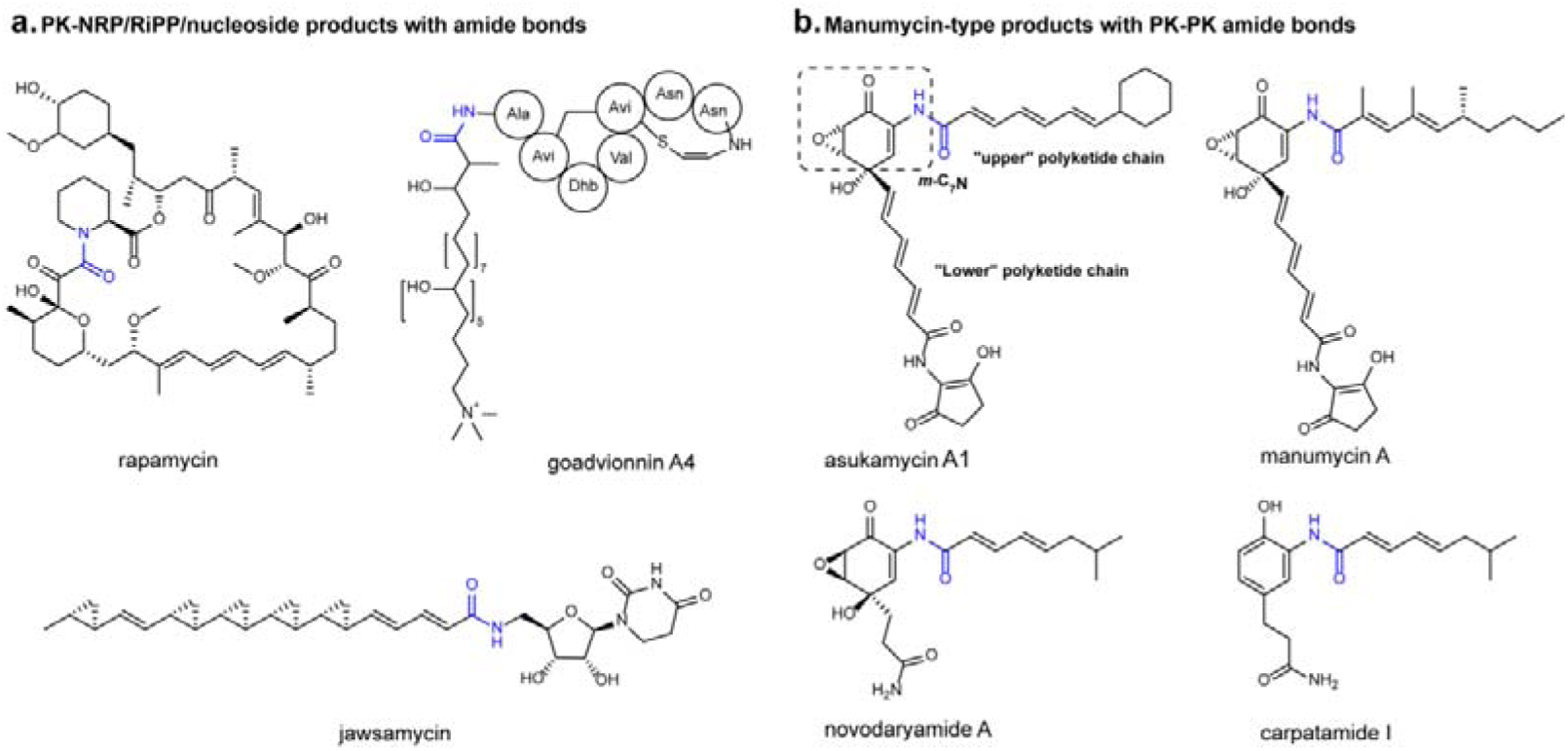
Representative polyketide-containing natural products involving amidative condensation in biosynthesis. **a**. Representative PK-NRP, PK-RiPP, and PK-nucleoside natural products that involve amidative condensation in their biosynthesis. Amide bonds are highlighted in blue. Rapamycin represents a PK-NRP, goadvionnin A4 represents a PK-RiPP, and jawsamycin represents a PK-nucleoside. **b**. Representative manumycin-type natural products proposed to undergo amidative condensation of PK-PK in their biosynthesis.

Manumycin-type metabolites comprise over 50 members, including asukamycin^10^, novodaryamide^9^, and carpatamide^11^ (Fig. 1b) with potent anticancer activity^9,11–15^. They inhibit farnesyl transferase to block oncogenic RAS maturation or act as molecular glues between UBR7 and TP53 to trigger p53-dependent cell death^16,17^. These compounds feature an “upper” polyketide chain connected via a central amide bond to a “lower” chain containing an m-C_7_N core. In asukamycin, these chains are produced by distinct PKS systems—an unusual system involving specialized β-ketoacyl-acyl carrier protein synthase III (KAS III) enzymes for the “upper” chain and a highly reducing type II PKS for the lower chain^8,18^. Condensation of these chains yields protoasukamycin, which is subsequently tailored oxidatively to install the epoxyquinone moiety essential for bioactivity^19^. Despite the elucidation of these pathways, the enzyme responsible for the pivotal amide-forming condensation has remained uncharacterized. While earlier work suggested an arylamine N-acetyltransferase (NAT) might mediate this step at the level of a dual acyl carrier protein (ACP) (Extended Data Fig. 1)^8,9^, arylamine NATs are primarily recognized for xenobiotic detoxification through acetyl transfer from acetyl-CoA to arylamines^20–22^. Their involvement in natural product biosynthesis is currently limited to only a few cases, including N-acylation of arylamines with diterpene acyl-CoA and lactam formation in ansamycins assembly (Extended Data Fig. 2)^23–25^. Thus, elucidating the mechanism of this condensation step in manumycin-type pathways is of great significance for both PKS biosynthesis and the mechanisms of arylamine NATs.

Using novodaryamide A as a model, we herein demonstrate that the NAT homolog Dar12 catalyzes an unprecedented intermolecular amidation between a free “lower” polyketide chain and an ACP-tethered “upper” chain. Our biochemical and structural data reveal that Dar12 and its homolog ColC2 constitute a distinct arylamine NAT subclass specialized for polyketide amidative chain transfer/condensation. Unlike canonical arylamine NATs, these enzymes utilize a unique substrate-entrance tunnel and binding mode to accommodate diverse acyl donors and acceptors. This broad substrate tolerance enables the convergent synthesis of diverse non-natural proto-manumycin analogues, establishing these enzymes as powerful biocatalysts for combinatorial polyketide biosynthesis.

## Results

### Elucidating the Condensation of “Upper” and “Lower” Polyketide Chains in Novodaryamide A Biosynthesis

We initially aimed to investigate the asukamycin enzymes,^8,18^ however, the expression of the arylamine NAT (AsuC2) and the ketosynthase/chain-length factor heterodimer (AsuC13/AsuC14)—key enzymes responsible for synthesis of the “lower” chain in *Escherichia coli*—yielded insoluble proteins. To address this, we shifted our focus to the enzymes in the novodaryamide A biosynthetic gene cluster (BGC) of *Streptomyces* sp. CNQ-085^9,26^. Dar6 is a didomain protein containing two ACP domains. Its N-terminal (Dar6_N) is structurally homologous to the “lower” chain-carrying ACP AsuC11 of asukamycin, according to AlphaFold2 predictions (Supplementary Fig. 1). Consequently, both Dar12 (60.31% identity to AsuC2) and Dar6_N (48.28% identity to AsuC11) were selected for expression in *E. coli*.

The preliminary hypothesis regarding the condensation of the “upper” and “lower” chains suggests that these reactions occur at the dual ACP level.^8,9^ Specifically, the condensation takes place between the “upper” and “lower” polyketide chains bound to the individual ACP, facilitated by the catalytic activity of an arylamine NAT. This enzymatic process will lead to the formation of a single, elongated polyketide chain product, which remains bound to the “lower” ACP (Extended Data Fig. 1).

To investigate this further, we first prepared the “lower” chain bound to Dar6_N. 3-(3-amino-4-hydroxyphenyl)-propanoic acid (**1**) was converted to its CoA form using the promiscuous CoA synthetase UkaQ^FAV^ (ref 27) and subsequently loaded onto Dar6_N via the phosphopantetheinyl transferase *Sfp*^28^ in a two-step, one-pot reaction (reaction mixture A, Extended Data Fig. 3). The ACP-bound “upper” chain was generated using the established asukamycin PKS system^18^, including AsuC3 (KASIII), AsuC5 (ACP), AsuC7 (ketoreductase), AsuC8/C9 (dehydratase) with isovaleryl-CoA as the starter unit (reaction mixture B, Extended Data Fig. 3b). However, when these two reaction mixtures were combined with Dar12, no Dar6_N-bound condensation product was detected by HPLC-HRMS (Extended Data Fig. 3c). To rule out the possibility that Dar12 does not recognize the “upper” chain ACP (AsuC5) from asukamycin, we replaced AsuC5 with Dar15 from the novodaryamide A biosynthetic pathway and used the Dar15-bound “upper” chain as the substrate (reaction mixture B, Extended Data Fig. 3b). However, the results remained unchanged.

Nevertheless, to our delight, we detected a compound (**2**) with a molecular weight corresponding to the ACP-free condensation product in both reactions (Fig. 2, trace I/III). The formation of **2** could be due either to the spontaneous hydrolysis of the Dar6_N-bound condensation product or direct condensation of the free molecule **1**, which remained in the reaction mixture, with the ACP-bound “upper” chain. To confirm this, we incubated **1** with Dar12 and either the AsuC5- or Dar15-bound “upper” chain. Both reactions yielded **2** (Fig. 2, trace II/IV), confirming that Dar12 indeed does not accept the ACP-bound “lower” chain but rather the free polyketide chain. Moreover, adding chemically synthesized 3-(3-amino-4-hydroxyphenyl)-propanamide (**3**) resulted in the production of carpatamide I (Fig. 2, trace V), although with lower conversion compared to **1**. This suggests that Dar12 has flexible substrate specificity, albeit its natural substrate is **1**.

**Fig. 2.**
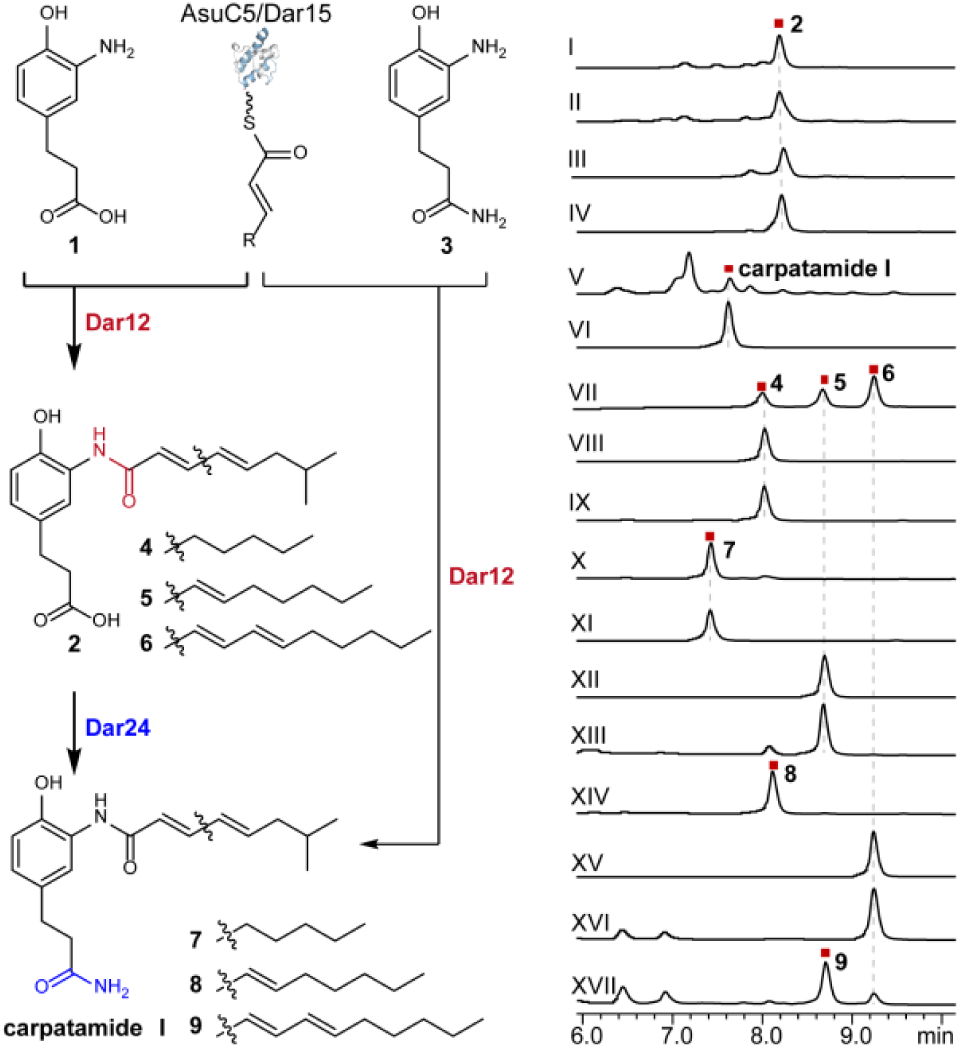
UPLC analysis of the *in vitro* enzymatic reactions and the feeding assay. (I) Compound **2** was synthesized by combining the reaction mixture A (**1**, HSCoA, ATP, UkaQ^FAV^, Sfp, MgCl_2_, and Dar6_N), reaction mixture B (isovaleric acid, HSCoA, ATP, UkaQ^FAV^, malonic acid, *Sc*MatB, *Sa*FabD, AsuC3, AsuC5, AsuC7, AsuC8/C9^18^), and Dar12; (II) Compound **2** was synthesized by combining **1**, reaction mixture B and Dar12; (III) Compound **2** was synthesized by combining the reaction mixture A, reaction mixture B (AsuC5 was replaced by Dar15), and Dar12; (IV) Compound **2** was synthesized by combining **1**, reaction mixture B (AsuC5 was replaced by Dar15), and Dar12; (V) Carpatamide I was synthesized from **3**, reaction mixture B, and Dar12; (VI) Carpatamide I standard isolated from *Streptomyces parvus* 1268^16^; (VII) Compound **4**, **5**, **6** were biosynthesized by combining **1**, reaction mixture B (with isovaleric acid was replaced to hexanoic acid) and Dar12; (VIII) Compound **4** standard; (IX) Compound **4** fed to *S. lividans* TK24; (X) Compound **4** fed to *S. lividans* TK24 harboring Dar24, producing **7**; (XI) Compound **7** standard; (XII) Compound **5** standard; (XIII) Compound **5** fed to *S. lividans* TK24; (XIV) Compound **5** fed to *S. lividans* TK24 harboring Dar24, producing **8**; (XV) Compound **6** standard; (XVI) Compound **6** fed to *S. lividans* TK24; (XVII) Compound **6** fed to *S. lividans* TK24 harboring Dar24, producing **9**. Note: Panels V and XV–XVII were detected at λ = 300 nm; all others were detected at λ = 254 nm.

To clarify the terminal amidation step, we expressed Dar24, an asparagine synthetase capable of carboxyl amidation. While Dar24 was insoluble in *E. coli*, it exhibited enzymatic activity when expressed in *Streptomyces lividans* TK24. Supplementing *S. lividans* cultures expressing Dar24 with **1** did not yield **3** (Supplementary Fig. 2). However, feeding synthesized compounds **4**–**6** led to the production of amidated products **7**–**9** (Fig. 2, traces X, XIV, XVII), establishing that terminal amidation by Dar24 occurs after polyketide chain condensation.

These findings provide a clear elucidation of the key polyketide chain transfer/condensation step in the biosynthesis of manumycin-type metabolites, which occurs between an ACP-bound “upper” chain and a free “lower” chain. To our knowledge, such convergent amidation between two polyketide chains has not been reported before. Furthermore, the function of the arylamine NAT is also unprecedented, which represents a new class of polyketide chain transfer/condensation enzymes that catalyze amidation.

### Deciphering the Catalytic Mechanism and the Unique Substrate-Binding Mode of ColC2

To investigate the transfer/condensation mechanism, we attempted to solve the structure of Dar12, however, crystallization was unsuccessful. As an alternative, we determined the structure of ColC2, a homologous arylamine NAT from the colabomycin BGC^29^. ColC2 shares 50.18% sequence identity with Dar12 and exhibits identical condensation and amidation activities (Extended Data Fig. 4).

The ligand-free ColC2 structure diffracted to 2.15 Å resolution and belonged to space group P2□2□2□. The asymmetric unit contained two ColC2 molecules that are highly similar, with a Cα RMSD of only 0.242 Å over 273 residues (Extended Data Fig. 5a). ColC2 adopts the conserved arylamine NAT fold, comprising three domains: domain I (α-helical bundle, residues 1–85), domain II (β-barrel, residues 86–178), and domain III (lid domain, residues 207–273), domain II and III were connected by an interdomain region (residues 179–206). The conserved catalytic triad—Cys72, His110, and Asp125—is positioned identically to that in other arylamine NATs (Extended Data Fig. 5b). Site-directed mutagenesis confirmed the functional importance of this triad: the C72A mutant completely lost activity, whereas the H110A and D125A mutants exhibited substantially reduced catalytic efficiency (Fig. 3f). Together, these results indicate that ColC2 employs a canonical arylamine NAT catalytic mechanism, in which Cys72 attacks the thioester bond of the ACP-bound “upper” chain to form a transient acyl–enzyme intermediate with concomitant release of the ACP, followed by transfer of the acyl group to the amine moiety of the “lower” chain to generate the condensation product.

**Fig. 3.**
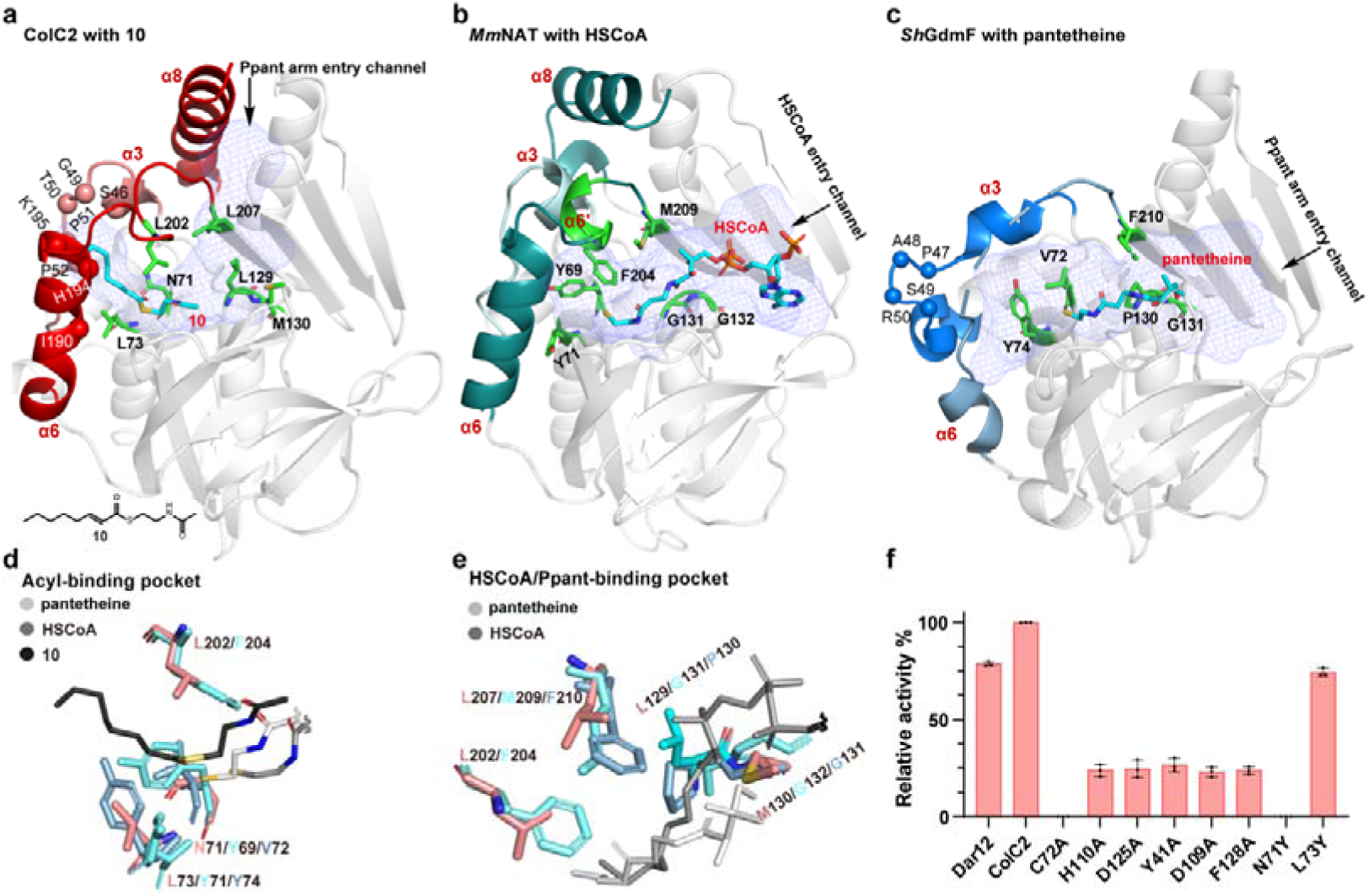
Structural and mutational analysis of ColC2. **a**, Crystall structure of ColC2 in complex with **10** (PDB ID: 8K56). Cα atoms of S46, G49, T50, P51, P52, I190, H194, K195 in α3 and α6 are shown as spheres. The Chemical structure of **10** is shown in the bottom. **b**, Complex structure of *Mm*NAT with HSCoA (PDB ID: 2VFC). **c**, Complex structure of *Sh*GdmF with pantetheine (PDB ID: 8OOM). Cα atoms of P47, A48, S49, R50 are shown as spheres. The substrate binding pockets are shown as slate mesh. Substrates are colored in cyan. Architecture difference among ColC2, *Mm*NAT and *Sh*GdmF are highlighted in red, blue and deep teal, respectively. 4′-phosphopantetheine (Ppant) arm of ACP/CoA entry channels are indicated by arrows. **d**, Enlarged view of the acyl-binding pocket of ColC2, *Mm*NAT and *Sh*GdmF. Compound **10** (dark), The SNAC moiety of pantetheine (white) and HSCoA (grey) is shown. L73, N71, L202 in CoCl2 in are colored in salmon; Y69, Y71 and F204 in *Mm*NAT are colored in cyan; V72, Y74 in *Sh*GdmF are colored in blue. **e**, HSCoA/Ppant-binding pocket of ColC2, *Mm*NAT and *Sh*GdmF. Only the Ppant arm of HSCoA (grey) is shown. N71, L73, L202, L207, L129 and M130 in ColC2 are colored in salmon; Y69, Y71, F204, M209, G131 and G132 in *Mm*NAT are colored in cyan; V72, Y74, F2210, P130 and G131 in *Sh*GdmF are colored in blue. **f**, Analysis of the relative reactivity of ColC2 and its mutants. Error bars, standard deviation (s.d.) of three independent experiments (n=□3).

Although ColC2 utilizes a conserved catalytic mechanism, its substrate specificity and reaction type differ markedly from those of previously characterized arylamine NATs. Based on activities, arylamine NATs can be divided into three types. Type I enzymes are acyl-CoA–dependent arylamine NATs that catalyze the acylation of small aromatic amines to form simple amide products^22,23^. Type II enzymes are ACP-dependent cyclization NATs that participate in ansamycin family biosynthesis; among these, *Sh*GdmF (PDB ID: 8OOM) was only recently characterized from the geldanamycin biosynthetic pathway and catalyzes intramolecular cyclization of ACP-bound polyketide chains to generate macrolactams^24^. Type III are ColC2 and its homologues.

To gain deeper insight into the substrate-binding architecture of ColC2, we determined the crystal structure of ColC2 in complex with *trans*-2-octanoyl-SNAC (**10**) at 2.42 Å resolution (PDB ID: 8K56), which mimics the ACP-tethered “upper” chain. Attempts to obtain a complex structure with **1** were unsuccessful. The complex structure closely resembles the ligand-free structure, with a Cα RMSD of only 0.162 Å over 273 residues (Extended Data Fig. 5c). **10** is accommodated within a large, curved substrate-binding pocket, with its acyl moiety positioned in an elongated cavity surrounded by residues S46, G49, T50, P51, P52, N71, C72, L73, I190, I191, H194 and K195 (Fig. 3a). Notably, the acyl group was not transferred to the catalytic residue Cys72, likely due to crystallization conditions that impeded progression of the catalytic reaction. The distance between the thiol group of Cys72 and the carbonyl carbon of **10** was 4.2 Å and 4.5 Å in chains A and B, respectively, consistent with its proposed role in catalysis (Extended Data Fig. 5d).

To elucidate the structural basis for functional divergence among arylamine NATs, we compared the ColC2–**10** complex with the *Mm*NAT-HSCoA complex structure from *Mycobacterium marinum* (Type I, PDB ID: 2VFC, Fig. 3b) and the *Sh*GdmF-pantetheine complex structure (Type II, PDB ID: 8OOM, Fig. 3c)^24,30^. Structural alignment revealed that ColC2 more closely resembles *Mm*NAT than *Sh*GdmF (Fig. 3a–c), despite phylogenetic analyses indicating a more recent common ancestor between ColC2 and *Sh*GdmF (Extended Data Fig. 6). Although all three enzymes share a conserved arylamine NAT fold, pronounced differences were observed in helices α3, α6, and α8, including (1) *Sh*GdmF is truncated by approximately 20 amino acids and completely lacks the C-terminal helix α8, whereas *Mm*NAT retains α8 but displays a pronounced mid-helix kink that results in a significant angular deviation compared to the straight α8 observed in ColC2 (Fig. 3a–c; Extended Data Fig. 7a). (2) Relative to ColC2, helix α6 in *Mm*NAT is deflected inward toward the substrate-binding pocket, with the adjacent loop adopting a new α-like conformation (α6’). In *Sh*GdmF, this region comprises 12 unresolved residues, suggesting increased flexibility that accommodates an enlarged substrate-binding pocket (Fig. 3a–c; Extended Data Fig. 7a). (3) Helix α3 in *Sh*GdmF is interrupted between Pro47 and Arg50, producing a marked angular deviation relative to the corresponding helices in ColC2 and *Mm*NAT (Fig. 3a–c; Extended Data Fig. 7a). Collectively, these structural differences in helices α3, α6, and α8 are critical for shaping substrate-binding pocket, likely underlying the enzymes’ distinct substrate specificities and catalytic behaviors.

The substrate-binding pocket of arylamine NATs consists of two distinct regions: one that accommodates the acyl moiety and another that binds the Ppant arm of ACP/CoA and arylamine component, with the catalytic cysteine positioned between these sites. Comparative analysis revealed that ColC2 exhibits a markedly different substrate-binding mode and access channel compared to *Mm*NAT and *Sh*GdmF. In *Mm*NAT, the acyl-binding pocket is primarily formed by residues Y69, Y71, and F204, with Y69 and Y71 creating an enclosed base (Fig. 3d). In *Sh*GdmF, the corresponding residues are V72 and Y74, while the residue corresponding to F204 is part of a 12-residue segment that remains unresolved in the crystal structure (Fig. 3e). This segment is believed to be flexible, accommodating the coiled polyketide chain of geldanamycin.^24^ In ColC2, these positions correspond to N71, L73, and L202, respectively (Fig. 3c). The side chain of N71 extends outward from the pocket, significantly reducing steric hindrance. Additionally, the α6 helix in ColC2 shifts outward by 12° relative to *Mm*NAT, further enlarging the acyl-binding pocket (Extended Data Fig. 7a) and forming a solvent-accessible cavity that accommodates very long polyketide acyl chains (Fig. 3a). To assess the functional relevance of N71 and L73, they were individually mutated to tyrosine. The N71Y mutation resulted in a complete loss of enzymatic activity, while the L73Y mutation had minimal effect (Fig. 3f), confirming that N71 is a critical determinant of acyl-pocket size.

In *Mm*NAT and *Sh*GdmF, CoA or pantetheine of ACP enters the pocket via a lateral channel on the right (Fig. 3b/c). Examination of the CoA/pantetheine-binding pocket revealed that two small residues at the base of the channel in *Mm*NAT and *Sh*GdmF—Gly131/Gly132 and Pro130/Gly131, respectively—are replaced by bulkier residues Leu129/Met130 in ColC2, effectively blocking the original CoA/pantetheine channel (Fig. 3a/e). To further characterize this altered channel, as well as the arylamine-binding pocket, the ColC2 product protocolabomycin acid (**11**) was docked into the substrate-binding cavity (Fig. 4a). The docking results indicated that **11** fits well within the curved substrate pocket, with its “upper” chain occupying a position identical to the acyl moiety of **10**. The amide group forms hydrogen bonds with residues Y41 and D109, while the aromatic ring engages in inclined face-to-face interactions with Y41 and F128. Analysis of other arylamine NATs revealed that Y41 and F128 are highly conserved, occasionally substituted by tyrosine, tryptophan, or histidine (Supplementary Fig. 3). Mutation of Y41, F128, or D109 to alanine resulted in a marked reduction in ColC2 activity (Fig. 3f), confirming their pivotal roles in stabilizing the arylamine moiety.

**Fig. 4.**
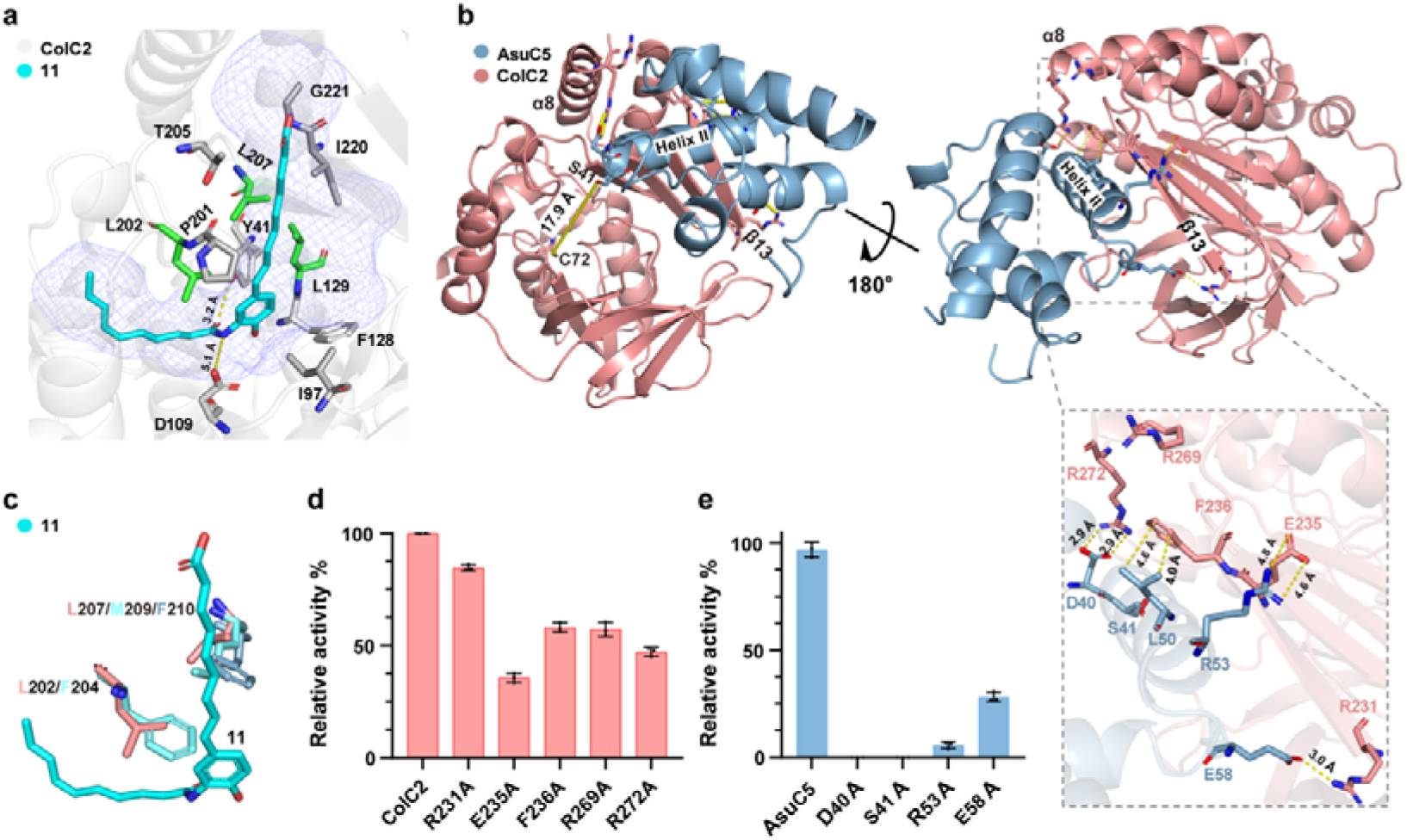
Docking model of ColC2-**11** and ColC2-AsuC5. **a**, Docking model of ColC2-**11**. The binding cavity formed by residues I97, L129, P201, L202, T205, L207, I220, and G221 is shown, with key critical residues highlighted in green. Hydrogen bonds between ColC2 and **11** are indicated by yellow dashed lines. **b**, Docking model of ColC2-AsuC5. Salt bridges, hydrophobic interaction and distance are indicated by yellow dashed lines. **c**, Enlarged view of critical residues of ColC2-**11**, *Mm*NAT and *Sh*GdmF. L202 and L207 in CoCl2-**11** are colored in salmon; F204 and M209 in *Mm*NAT are colored in cyan; F210 in *Sh*GdmF is colored in blue. **d**, Analysis of the relative reactivity of ColC2 mutants, which were predicted to be interacted with AsuC5. **e**, Analysis of the relative reactivity of AsuC5 mutants. D40, R53 and E58 were predicted to be interacted with ColC2; S41 is the conserved catalytic residues. All experiments were repeated independently three times.

As anticipated, the “lower” chain of **11** adopts a nearly perpendicular orientation relative to the acyl-binding pocket and is accommodated within a cavity created by residues I97, L129, P201, L202, T205, L207, I220, and G221 (Fig. 4a). In *Mm*NAT, this pocket is occluded by two bulky residues, F204 and M209, whereas ColC2 contains smaller residues, L202 and L207, in these positions. *Sh*GdmF also contains a large residue F210, corresponding to M209, while the residue corresponding to F204 is part of a missing 12-residue segment (Fig. 4c). These structural differences are thought to create a new pocket in ColC2 that accommodates the Ppant arm of ACP and the “lower” chain. Collectively, these structural rearrangements establish a fundamentally distinct mechanism for substrate entry and binding in ColC2 compared to other arylamine NATs.

### Elucidating the interaction between ColC2 and ACP

Based on above analysis of the upward channel, ACP should be positioned near the α8 helix of ColC2. To verify this assumption, we investigated the interaction interface between ColC2 and its cognate ACP, AsuC5, using AlphaFold2 modeling combined with molecular dynamics (MD) simulations. The AlphaFold2-predicted structure of AsuC5 revealed a three-dimensional architecture similar to that of *E. coli* AcpP (PDB ID: 7L4E; Extended Data Fig. 7b)^31^. Docking of the ColC2–AsuC5 complex using HADDOCK2.2^32^, followed by Molecular dynamics (MD) simulations and per-residue energy decomposition analyses (Supplementary Fig. 4), indicated that helices α8 and β13 of ColC2 interact primarily with helix II of AsuC5 (Fig. 4b).

The conserved residue S41 of AsuC5, located at the base of helix II, is positioned near helix α8 of ColC2 and oriented toward the catalytic residue Cys72, at a distance of approximately 17.9 Å—comparable to the ∼20 Å length of the Ppant arm (Fig. 4b). The ColC2–AsuC5 complex is stabilized by multiple salt bridges, cation–π interactions, hydrogen bonds, and electrostatic interactions (Supplementary Table 1), consistent with canonical ACP–partner protein interfaces^33^. Specifically, residues R269, R272, E235, and R231 of ColC2 form salt bridges with residues D40, R53 (helix II), and E58 of AsuC5 (Fig. 4b). In addition, F236 of ColC2 engages in hydrophobic interactions with L50 of AsuC5. To validate these interactions, alanine substitution mutants were generated for ColC2 (R231A, E235A, F236A, R269A, and R272A) and AsuC5 (D40A, R53A, and E58A). Functional assays demonstrated that amide bond formation activity was severely impaired in both ColC2 and AsuC5 mutants (Fig. 4c/d), confirming that these salt bridges and hydrophobic contacts are critical for ACP recognition. Together, these interactions enable ColC2 to recognize ACP-tethered polyketide chains and catalyze condensation and amidation with a second, free polyketide chain.

Multiple sequence alignment analysis revealed that these ColC2–AsuC5 interface residues are generally conserved among the type III arylamine NATs from other manumycin-type BGCs, albeit with some variability (Supplementary Fig. 3). In contrast, type II NATs appear to employ a distinct mode of ACP engagement; for example, residues corresponding to R231 and E235 in ColC2 are replaced by V234 and K238 in *Sh*GdmF, while residues corresponding to R269 and R272 are absent altogether. This variability in ACP-interaction motifs mirrors the diversity observed in other ACP–enzyme recognition systems.

### Evaluating the Substrate Diversity of ColC2 and Its Application in Convergent Synthesis of Manumycin-Type Compounds

The structural and mechanistic analysis described above suggest that ColC2 may possess broad substrate specificity. To validate this hypothesis, we systematically evaluated its substrate scope for the generation of non-natural proto-manumycin–type products. First, we examined whether ColC2 could utilize CoA- and SNAC-bound acyl donors. ColC2 was incubated separately with *trans*-2-octanoyl-CoA (**12**) or **10** in the presence of **1**. In both cases, ColC2 efficiently converted the substrates to compound **4** (Fig. 5a; Extended Data Fig. 8a, traces II and III; Supplementary Fig. 5a, traces II and III), demonstrating broad tolerance for acyl donors regardless of whether the thioester is linked to ACP, CoA, or the shorter SNAC moiety.

**Fig. 5.**
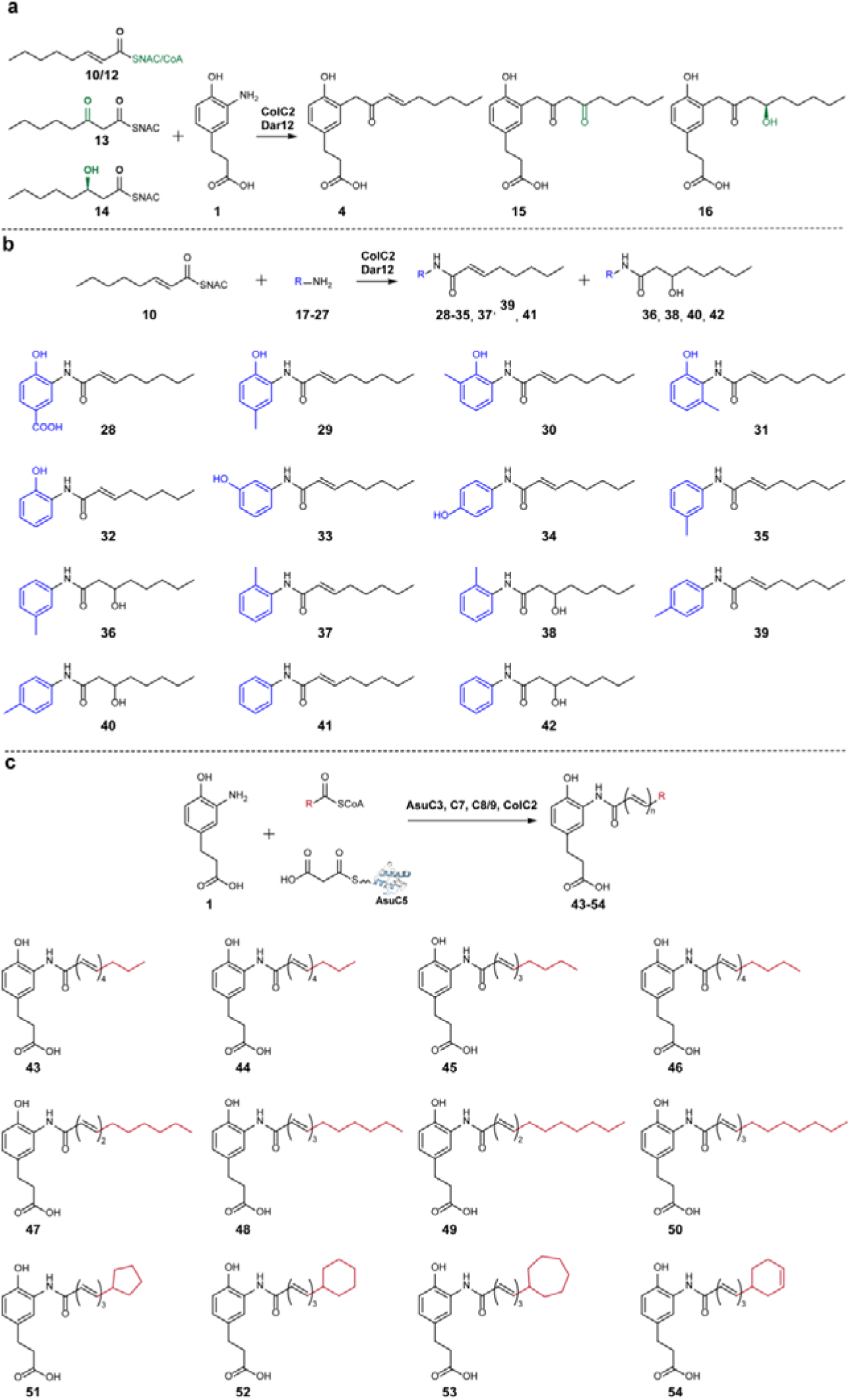
Diversity-oriented biosynthesis of unnatural manumycin-type compounds. **a**, Diversity of acyl donnors. **b**, Diversity of acyl acceptors. **c**, Practical utility of condensation system with changed starter units. Structures resulting from the changes are highlighted in green, blue and red, respectively.

Previous studies have shown that omission of ketoreductase and dehydratase domains during “upper” chain assembly leads to the accumulation of polyketone or polyhydroxylated polyketide intermediates^18^. To assess whether ColC2 can accept such intermediates, we incubated the enzyme separately with synthetic 3-oxooctanoyl-SNAC (**13**) and (*R*)-3-hydroxyoctanoyl-SNAC (**14**) in the presence of **1**. ColC2 successfully utilized both substrates as acyl donors, yielding compounds **15** and **16**, respectively (Fig. 5a; Extended Data Fig. 8a, traces V and VI; Supplementary Fig. 5a, traces V and VI).

Next, we evaluated the specificity of ColC2 toward a diverse set of arylamine acceptors. Eleven arylamines—3-amino-4-hydroxybenzoic acid (3,4-AHBA, **17**), 2-amino-4-methylphenol (**18**), 2-amino-6-methylphenol (**19**), 2-amino-3-methylphenol (**20**), 2-aminophenol (**21**), 3-aminophenol (**22**), 4-aminophenol (**23**), m-toluidine (**24**), o-toluidine (**25**), p-toluidine (**26**), and aniline (**27**)—were incubated with **10**. ColC2 accepted all tested arylamines, producing the corresponding amide products **28–42** (Fig. 5b; Extended Data Fig. 8b; Supplementary Fig. 5b). Notably, for substrates **24–27**, both the expected products (**35, 37, 39, 41**) and their spontaneous hydration derivatives (**36, 38, 40, 42**) were detected. The structures of compounds **29** and **40** were confirmed by comparison with authentic standards.

Finally, to demonstrate the practical utility of ColC2 for convergent biosynthesis, we tested whether the enzyme could utilize ACP-tethered “upper” chains derived from diverse starter units. Eight alternative starter substrates were evaluated by modifying the standard reaction used for compound **2**. Linear fatty acids (n-butyric, n-valeric, n-caproic, and n-octanoic acids) were substituted for isovaleric acid, while cyclic carboxylic acids, including cyclopentanecarboxylic, cyclohexanecarboxylic, cycloheptanecarboxylic, and 3-alkenylcyclohexenecarboxylic acids were introduced using the acyl-CoA synthetase AliA_100_^34^ in place of UkaQ^FAV^. All tested substitutions successfully yielded corresponding compound **2** derivatives (**43-54**) (Fig. 5c; Extended Data Fig. 8c).

Collectively, these results demonstrate that ColC2 exhibits remarkably relaxed specificity toward both acyl donors and arylamine acceptors. This broad substrate tolerance establishes ColC2 as a versatile biocatalyst for the convergent biosynthesis of polyketide and highlights its potential applications in chemoenzymatic synthesis.

## Discussion

Manumycin-type metabolites employ an unconventional amidative chain transfer/condensation strategy that links an ACP-tethered polyketide chain with an ACP-free polyketide chain. This mode of chain coupling has not previously been documented in polyketide biosynthesis. By reconstituting the pathway in vitro, we elucidate for the first time the mechanistic logic underlying the amidative chain transfer/condensation step in novodaryamide A biosynthesis. We demonstrate that this key transformation is catalyzed by the unique arylamine NAT Dar12, while the two polyketide precursors are independently assembled by a KASIII-mediated PKS and a highly reducing type II PKS, respectively.

Amidative chain transfer/condensation reactions involving polyketides and amino acids, peptides, or nucleosides are well documented in natural product biosynthesis. However, these reactions are typically mediated by C domains of NRPSs or by TE domains (Extended Data Fig. 9)^4,35–38^. In other cases, GCN5-related NATs have been implicated, such as GdvG in the biosynthesis of the PK/RiPP hybrid lipopeptide goadvionin^6^, Jaw2 in the polyketide–nucleoside hybrid jawsamycin^7^, and the transglutaminase homolog AdmF in andrimide biosynthesis^39^. In contrast, Dar12 represents the first example of an arylamine NAT that catalyzes direct condensation and amidation between two polyketide chains, establishing a fundamentally new paradigm for amide bond formation in polyketide assembly.

Arylamine NATs are widely distributed in humans and microorganisms, where they typically utilize acetyl-CoA (C_2_ donors) to acetylate xenobiotic substrates for detoxification.^21,22^ However, this work, together with emerging studies^23,24^, reveals that arylamine NATs can catalyze atypical acyl-transfer reactions using non–acetyl-CoA donors during natural product biosynthesis. Structural analysis of ColC2 revealed a canonical arylamine NAT fold with distinct adaptations that enable this unique function. Specifically, replacement of conserved residue with smaller amino acids (N71) and swing the α6 helix in ColC2 outward by 12°, substantially expands the acyl-binding pocket, allowing accommodation of extended polyketide chains of up to at least C14. Additionally, substitution of a conserved glycine with methionine blocks the original CoA/ACP tunnel, while replacement of bulky residues with smaller ones (L202 and L207) generates a newly evolved acyl-acceptor tunnel oriented nearly perpendicular to the canonical acyl-binding channel. This architecture contrasts with the linear lateral channel observed in other NATs, such as *Mm*NAT and *Sh*GdmF.

AlphaFold2 modeling and MD simulations of the ColC2–AsuC5 complex further revealed that the Ppant arm of the ACP accesses this newly evolved channel. Based on these structural and biochemical observations, we propose the following catalytic mechanism (Fig. 6). First, the ACP-tethered “upper” polyketide chain binds to ColC2, inserting the Ppant-linked acyl chain into the vertical channel. The catalytic dyad His110 and Asp125 activates Cys72, which nucleophilically attacks the thioester to form an acyl–enzyme intermediate. Following ACP release, the ACP-free “lower” polyketide chain enters the channel, where its aromatic amine—positioned by residues such as Tyr41 and Phe128—is activated by the same catalytic dyad to attack the acyl–enzyme intermediate, yielding the amidated condensation product.

**Fig. 6.**
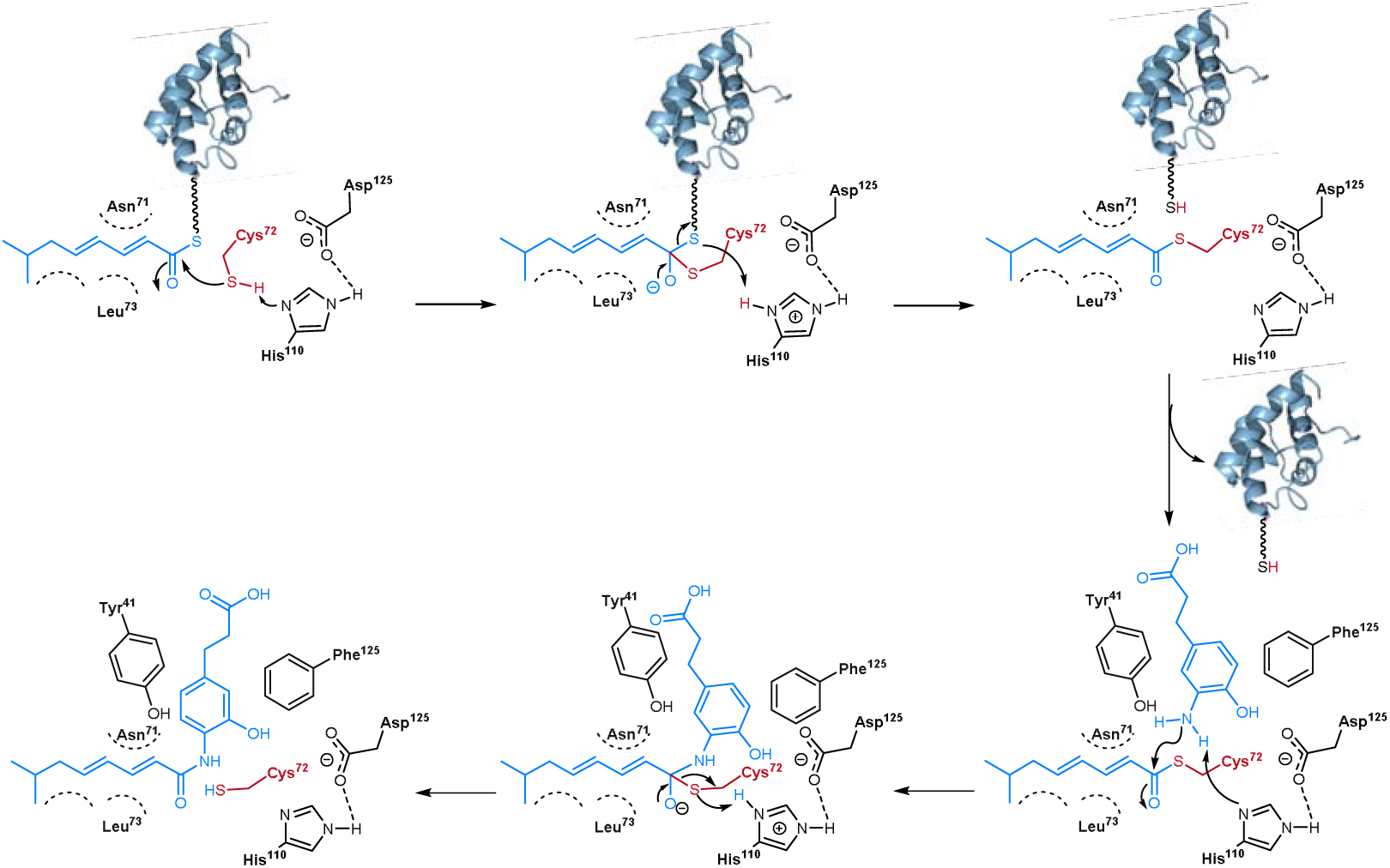
Proposed catalytic mechanism of ColC2 mediated condensation and amidation. ACP is colored in salmon, residues of ColC2 are colored in deepteal.

Notably, ColC2 and its homolog Dar12 exhibit remarkable substrate promiscuity. They efficiently recognize acyl donors tethered to ACP as well as small-molecule thioesters (CoA or SNAC), and they catalyze acyl transfer to a broad range of arylamine acceptors, including non-polyketide substrates. This catalytic flexibility highlights their potential as versatile biocatalysts for constructing non-natural manumycin-type analogs. Moreover, in contrast to the conventional stepwise assembly strategies of PKS–NRPS systems^4^, this convergent synthetic capability is far more amenable to engineering. It obviates the need to modify the specificity of individual modules or domains in the PKS for “lower” chains, which is often technically challenging and time-consuming. By combining a PKS with synthetic amine analogs, a wide range of novel hybrid polyketide scaffolds could potentially be generated. Thus, the discovery of ColC2 provides a powerful new enzymatic toolkit for polyketide engineering.

In summary, this study provides a comprehensive mechanistic understanding of the biosynthesis of manumycin-family metabolites and identifies the arylamine NAT Dar12 and its homolog ColC2 as a distinct class of polyketide chain transfer/condensation enzymes. These enzymes catalyze convergent amidation between an ACP-tethered “upper” polyketide chain and an ACP-free “lower” polyketide chain through a unique substrate-binding pocket and recognition mode. Together with extensive biochemical characterization demonstrating broad tolerance toward both acyl donors and acceptors, this work elucidates the core amide bond formation in manumycin-type biosynthesis and expands our understanding of both arylamine NAT function and polyketide scaffold assembly. These findings position Dar12 and its homologs as promising biocatalysts with potential applications in synthetic chemistry and drug discovery.

## Methods

### Gene expression and protein purification

The constructed plasmids pJTU37, pJTU40, pJTU42, pJTU44-pJTU47, pJTU117-pJTU120, pJTU122-pJTU141 for AsuC3, AsuC5, AsuC7, AsuC89, *Sc*MatB, *Sa*FabD, Sfp, Dar12, ColC2, Dar15, Dar6_N, Dar24, ColC2 mutants, and AsuC5 mutants were individually introduced into *E. coli* BL21(DE3), respectively. The seed cultures of *E. coli* BL21(DE3) harboring the plasmids were grown overnight at 37°C with shaking at 220 rpm in 5 mL LB media containing 50 μg mL^-1^ kanamycin (pET28a) or 100 μg mL^-1^ ampicillin (pET21a). The seed cultures were transferred to 800 mL LB medium containing the corresponding antibiotic in 2.5 L shaking flasks and grown at 37°C, 220 rpm, until the logarithmic growth phase was reached. Expression was induced with 0.1 mM isopropyl β-D-thiogalactopyranoside (IPTG) at 16°C, and the culture was continued at 16°C for 18 hours. The cells were then harvested by centrifugation at 4000 rpm for 20 minutes. The cell pellets were resuspended in 50 mL of buffer A (20 mM HEPES, 10% glycerol, pH 8.0, 500 mM NaCl) and lysed by sonication. The lysate was clarified by centrifugation at 12,000 rpm for 30 minutes at 4°C. The supernatant was loaded onto a 10 mL nickel affinity column (Cytiva) and eluted with buffer A containing a gradient of imidazole from 25 to 300 mM. Fractions containing pure proteins were identified by SDS-PAGE, then concentrated using centrifugal filter units (Amicon MWCO 10 kDa or 3000 Da, Millipore). The concentrated proteins were pooled and desalted by using a PD10 column (GE Healthcare) and stored in storage buffer (20 mM HEPES, 10% glycerol, pH 8.0, 100 mM NaCl) at -80°C.

### *In vitro* biochemical assay

For the Dar6_N-bound “lower” chain, a 100 μL reaction mixture A contained 1 mM **1**, 5 μM UkaQ^FAV^, 2 mM MgCl□, 5 mM ATP, 2 mM HSCoA, and 100 mM HEPES buffer (pH 8.0, 10% glycerol). The mixture was incubated at 30°C for 1 hour, then 50 μM Dar6_N and 10 μM Sfp were added, and the incubation continued at 30°C for an additional 3 hours.

For the “upper” polyketide chain, a 100 μL reaction mixture B contained 5 μM AsuC3, 50 μM holo-AsuC5 (or holo-AsuC5 mutants or apo-Dar15), 10 μM AsuC7, 10 μM AsuC8/C9, 5 μM Sfp, 5 μM UkaQ^FAV^, 5 μM *Sa*FabD, 5 μM *Sc*MatB, 5 μM GDH, 2 mM MgCl□, 5 mM ATP, 2 mM HSCoA, 3 mM malonic acid, 1 mM isovaleric acid or hexanoic acid, 1 mM NADP^+^, and 10 mM glucose in 100 mM HEPES buffer (pH 8.0, 10% glycerol). The mixture was incubated at 30°C for 4 hours.

For the assay of Dar12, ColC2, and ColC2 mutants, reaction mixture A or 1 mM **1**, 1 mM **3**, or 1 mM compound **17**-**27** were mixed with reaction mixture B or 1 mM *trans*-2-octanoyl-CoA/SNAC (**12/10**) or 3-oxa-octanoyl-SNAC (**13**). Then, 5 μM Dar12, ColC2, or ColC2 mutants were added. The reaction volume was adjusted to 200 μL using 100 mM HEPES buffer (pH 8.0, 10% glycerol), and the mixture was incubated at 30°C for 4 hours. After incubation, an equal volume of ethyl acetate (EA) was added, and the mixture was centrifuged at 12,000 rpm for 10 minutes, repeated three times. EA was removed by rotary evaporation, and the sample was dissolved in 30 μL acetonitrile for UPLC-MS and HR-ESI-MS analysis.

For Fig. 2, 3f, 4c/d; Extended Data Fig.4, 8a; Supplementary Fig. 5, products were analyzed on a Shimadzu LCMS-2020 coupled with a Shimadzu LC-40D XS UPLC equipped with a Shim-pack Scepter HD-C18 column (2.1 × 50 mm, 1.9 μm, Shimadzu). A gradient elution was applied with mobile phase A (H_2_O with 0.1% formic acid) and mobile phase B (acetonitrile with 0.1% formic acid) as follows: 5%-95% B (0-8 min), 95% B (8-9 min), 95%-5% B (9-10 min), 5% B (10-12 min). For Extended Data Fig. 8b, the gradient was set to 40% B (0-1 min), 40%-95% B (1-7 min), 95% B (7-11 min), 95%-10% B (11-13 min) with a flow rate of 0.2 mL/min.

HR-ESI-MS analysis was performed using a Thermo Scientific Orbitrap Exploris 120 equipped with a UMISIL™ C18 column (4.6 × 250 mm, 5 μm, Thermo Scientific). The gradient was set to 10% B for 1 minute, 10%-90% B for 15 minutes, 90% B for 4 minutes, and 10% B for 5 minutes with a flow rate of 1.0 mL/min. MS parameters were as follows: acquisition mass range, m/z 200−1000; TOF accumulation time, 0.1 s; collision energy, 30 eV; Ion Source Gas 1, 50 psi; Ion Source Gas 2, 50 psi; Curtain gas, 25 psi; Ion spray voltage (Floating), 5500 V; Temperature, 550°C. MS data were analyzed with TraceFinder software (Thermo Scientific). Supplementary Fig. 6 displays the mass spectrometry data of all compounds in this study.

### Feeding assay and products analysis

The plasmid pJTU121 carrying *dar24* was transformed into *E. coli* ET12567/pUZ8002 and further conjugated into *S. lividans* TK24 following the standard procedure^40^. After 7 days of cultivation, the apramycin-resistant colonies were picked and inoculated into 3 mL TSB medium (3g tryptic soya broth in 100 mL water) with 50 μg mL^-1^ apramycin. After growing at 30°C and 220 rpm for 3 days, the DNA of these candidate clones was extracted and used as templates for PCR to confirm the genotypes. For the feeding assay, a fresh seed culture of *S. lividans* TK24 containing *dar24* was inoculated into 250 mL flasks containing 50 mL fermentation medium (2 g glucose, 0.5 g bacto peptone, 25 mg K_2_HPO_4_, 25 mg MgSO_4_, 0.5 mg (NH_4_)_6_Mo_7_O_24_·4 H_2_O, 5 mg FeSO_4_·7 H_2_O, 0.5 mg CuSO_4_·5 H_2_O, 0.5 mg ZnSO_4_·7 H_2_O, 1 mg MnCl_2_·4 H_2_O in 100 mL water, pH 7.2). After 2 days, 1 mM of chemically synthesized compounds **4**, **5**, **6**, or **1** were added separately, and the cultures were further incubated for 1 day. After fermentation was completed, products were extracted with 50 mL EtOAc three times, and the organic phase was concentrated by vacuum. The residues were re-dissolved in MeCN and filtered through a 0.22 μm microporous membrane before being subjected to UPLC-MS analysis. The analytical procedures for compounds **7**, **8**, **9**, and **3** were identical to the method described for Fig. 2.

### Crystallization and crystal structure determination of ColC2

Initial purification of the protein was carried out as described above. Fractions containing pure ColC2 were identified by SDS-PAGE, pooled, concentrated using centrifugal filter units (Amicon MWCO 10 kDa, Millipore), and loaded onto a Superdex 75 Increase column (Cytiva) equilibrated with buffer B (20 mM Tris-Cl, pH 7.5, 100 mM NaCl). Fractions containing pure ColC2 were pooled and concentrated to 10 mg/mL for protein crystallization. Crystallization of ColC2 was screened with 6 × 96 conditions at 18°C using the sitting-drop vapor diffusion method. Crystals grew in the following condition: 0.1 M HEPES sodium (pH 7.5), 1.4 M lithium sulfate monohydrate. ColC2 crystals were cryoprotected in a reservoir solution containing 10% glycerol and frozen in liquid nitrogen for mounting prior to *X*-ray diffraction.

For the complex structure, crystals of *apo*-ColC2 were soaked in a reservoir solution containing 0.7 mM **10** and 10% glycerol before being frozen in liquid nitrogen for *X*-ray diffraction. *X*-ray diffraction data were collected at the Shanghai Synchrotron Radiation Facility BL02U1. The collected data were indexed and integrated using XDS, and scaled and merged using Aimless^41,42^. The structures were solved by molecular replacement using Phaser in Phenix^43^ with the atomic coordinates of PDBID: 4NV7 as a search model. The models were subjected to iterative rounds of rebuilding and refinement in Coot and Phenix^44,45^. The data collection and refinement statistics are summarized in Supplementary Table. 2. The atomic coordinates and the structure factors were deposited in the Protein Data Bank: *apo*-ColC2 (PDB: 8K51) and ColC2 complex (PDB: 8K56). The space of catalytic tunnels in the protein was analyzed using CavitOmiX (Version 1.0, 2022, Innophore GmbH), and the figures were prepared in PyMOL (Version 2.0, Schrödinger).

### Docking

For the docking of protocolabomycin acid to ColC2, the crystal structure of ColC2-**10** was used as the template. Docking was performed in Discovery Studio (Version 4.5, BIOVIA). The protein input file was prepared by adding hydrogen atoms, and the ligand, protocolabomycin acid (**11**), was built and energy-minimized using Chem3D. The binding pocket was defined as all residues within 11 Å of Cys72, based on the position of **10** in the pocket. Docking was carried out using the default LibDock parameters with 100 docking trials. For the complex structure of ColC2 and AsuC5, the HADDOCK2.2 webserver^32^ was used. Docking was performed using the ColC2 crystal structure and the AlphaFold2^46^ predicted model of *apo*-AsuC5. The docked complexes were visualized in PyMOL.

### Molecular dynamics simulations and analysis

All-atom MD simulations of the systems were conducted using GROMACS (Version 2021.1). The protein structure docked with substrates was solvated in a TIP3P cubic box, with a minimum distance of 1.0 nm between the solvate and the box edges. The system was neutralized by adding Na□ or Cl□ ions as needed. The protein was parameterized using the amberff14sb force field and parameter files.

Energy minimization was performed until the maximum force was reduced to less than 80 kJ mol^-1^, using both the steepest descent method and the conjugate gradient method. After energy minimization, a simulated annealing step was performed for 300 ps. The systems were gently heated using six 50-ps steps, incrementing the temperature by 50 K each step (0–300 K) under constant volume and periodic boundary conditions. Following simulated annealing, an NPT simulation was performed for 100 ps. Finally, individual production runs were executed for 10 ns in the NPT ensemble with random initial velocities generated to simulate different initial conditions, and a 20 ns MD run was performed (Supplementary Fig. 7).

The particle mesh Ewald method was used to evaluate the electrostatic interactions with a cut-off radius of 1.4 nm, and van der Waals interactions were calculated with a cutoff value of 1.4 nm. The bonds involving hydrogen atoms were constrained using LINCS with an order of 4. The temperature and pressure were regulated using the V-rescale method and the Parrinello-Rahman method, respectively. The trajectories were recorded every 2 fs. VMD was used to analyze the trajectory, and properties like potential energy, root mean square deviation (RMSD), minimal distance, radius of gyration (Rg), and H-bonding were calculated for the MD run. Graphs were plotted using XMGRACE software. Binding free energy prediction with decomposition and virtual alanine scanning were performed using gmx_MMPBSA^47^.

## Phylogenetic analysis of NAT

The phylogenetic tree was generated with MEGA X^48^ via the Neighbor-Joining method using the best-fit model G + F + I, according to the calculation. The tree was visualized in iTOL^49^.

## Data Availability

The structure of ligand free ColC2 and substrate complexes ColC2 have been deposited in the Protein Data Bank under ID numbers: 8K51 and 8K56, respectively.

## Acknowledgment

This work was supported by the National Natural Science Foundation of China (32425033 to X.Q., 32300044 to X.Y., 32370083 to M.L.), the China Postdoctoral Science Foundation (2023M732239 to X.Y.), National Key R&D Program of China (2025YFA0921000 to X.Q.) and LUI Che Woo Talent Development Fund (X.D.) and Shanghai Municipal Science and Technology Major Project (to X.Q. and X-D.K). We also thank the staff of beamline BL02U1 of Shanghai Synchrotron Radiation Facility (SSRF) for access and help with the X-ray data collection.

## Author contributions

X.Q., X-D.K., X.Y., G.W., Z.L., Z.D. conceived this project; X.Y. and Y.G. performed biochemistry experiment; X.Y. performed chemical synthesis; B.M. and X-D.K. solved the crystal structure; Y.G. conducted the docking and MD simulations; M.L. prepared the standard carpatamide I; Q.Z. provided helps for protein expression; X.Y. X-D.K. and X.Q. prepared the manuscript. All authors contributed to the manuscript editing.

## Additional information

**Supplementary Information** accompanies this paper at XXX.

## Competing interests

The authors declare no competing interests

---

**Extended Data Fig. 1.**
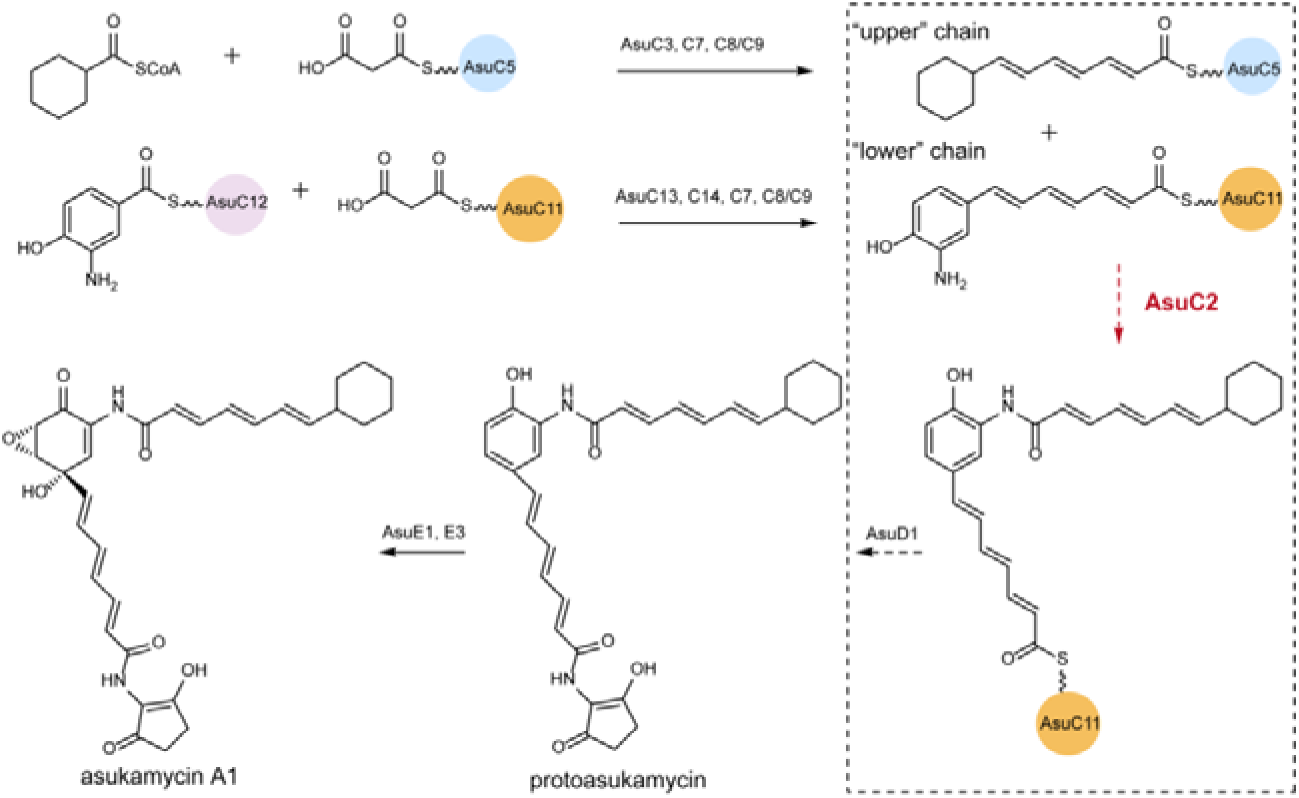
Proposed biosynthesis of asukamycin A1. The “upper” chain, bound to AsuC5, is generated by a KASIII-mediated PKS, while the “lower” chain, bound to AsuC11, is produced by HR II PKS. These two chains are proposed to be coupled by the arylamine NAT AsuC2, resulting in an elongated polyketide product tethered to AsuC11. This intermediate is then converted into asukamycin A1 by downstream tailoring enzymes. AsuC3 encodes KAS III; AsuC5 encodes ACP; AsuC7 encodes ketoreductase; AsuC8/C9 encode heterodimeric dehydratase; and AsuC13/C14 encode KSα/KS_CLF_ complex.

**Extended Data Fig. 2.**
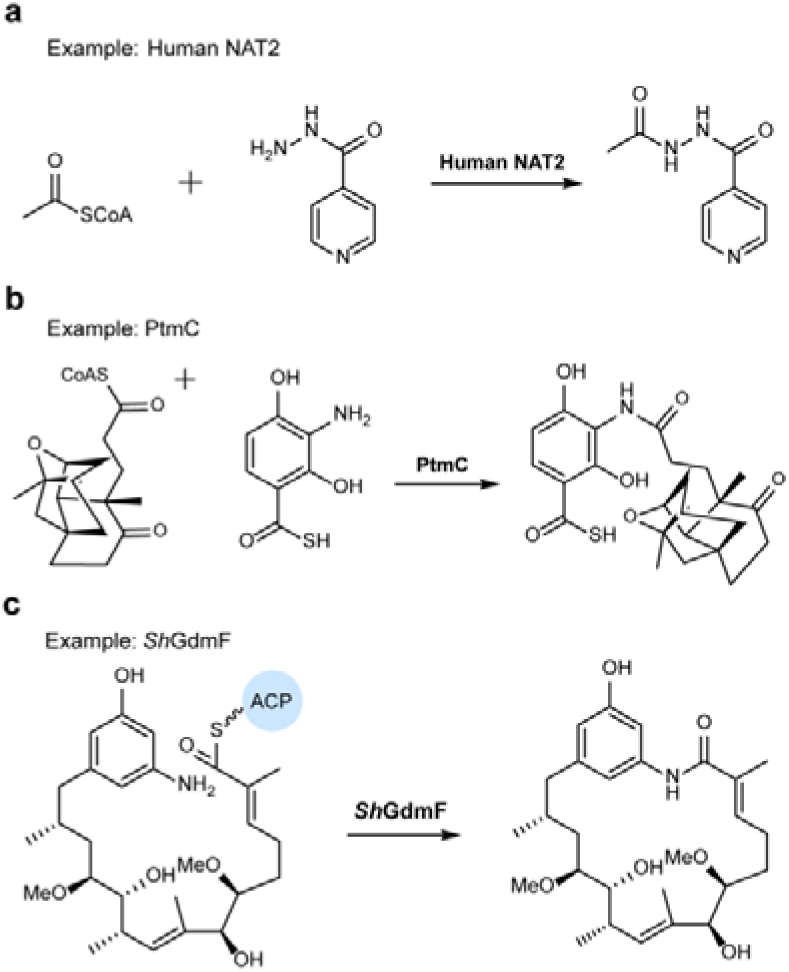
Examples of arylamine NATs involved in xenobiotic detoxification and natural products biosynthesis. **a**, Human NAT2, which is involved in the metabolism of isoniazid. **b**, PtmC catalyzes the N-acylation of arylamines using diterpene acyl-CoA in pladienolide biosynthesis. **c**, *Sh*GdmF catalyzes the lactamization of an ACP-tethered chain in geldanamycin biosynthesis.

**Extended Data Fig. 3.**
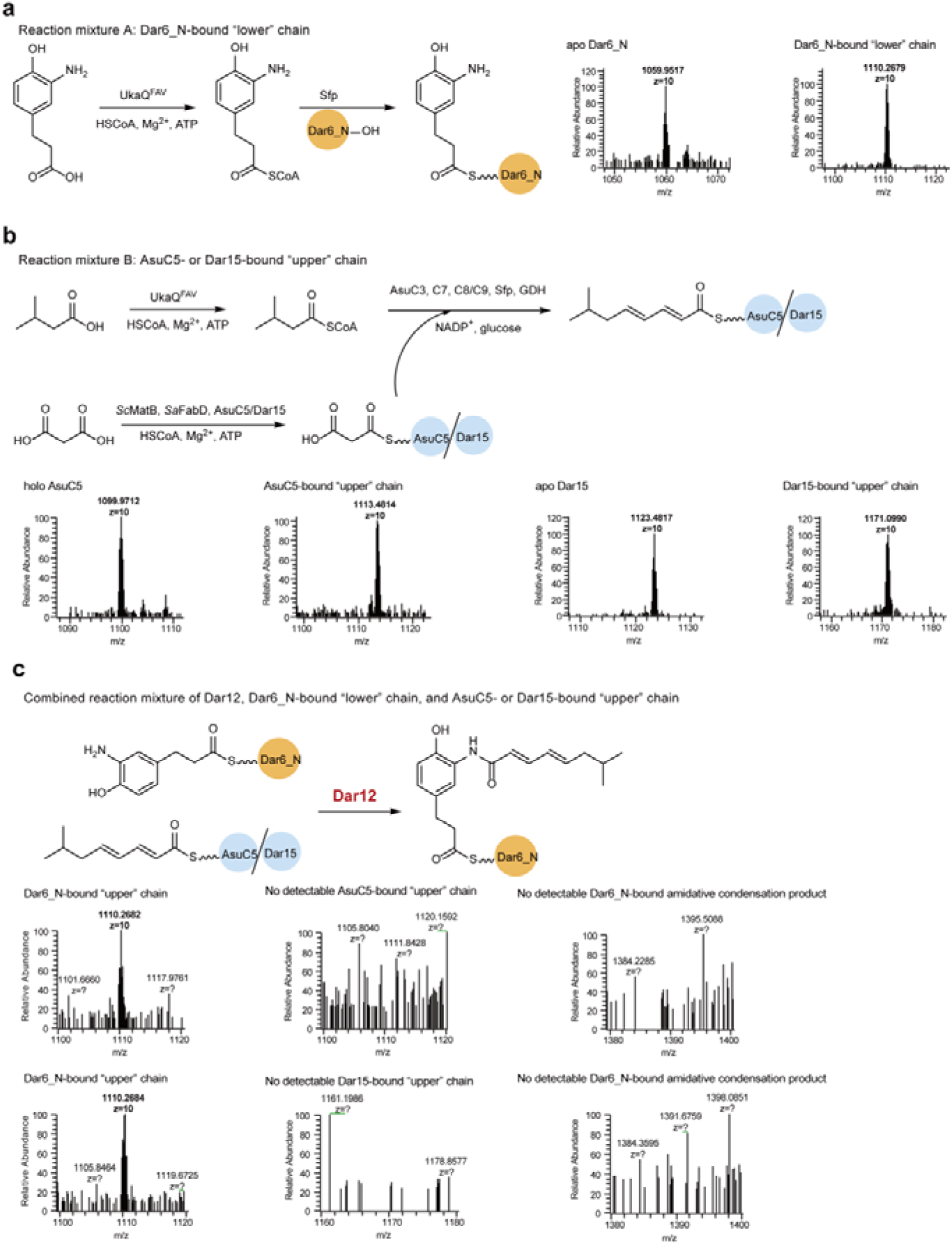
LC-HRMS analysis of reactions A and B their combined reaction. **a**, Reaction mixture A: The *m/z* values of the protein masses (molecular weight) for the +10 charge states (MS^1^) are shown. Measured (theoretical) masses *m/z* [M + 10H]^10+^ 1059.9517 (1059.9112) and 1110.2679 (1110.2618) correspond to *apo*-Dar6_N and the Dar6_N-bound “lower” chain, respectively. **b**, Reaction mixture B: Measured (theoretical) masses *m/z* [M + 10H]^10+^ values 1099.9712 (1099.9410), 1113.4814 (1113.5604), 1123.4817 (1123.4570), and 1171.0990 (1171.1095) correspond to *holo*-AsuC5, the AsuC5-bound “upper” chain, *apo*-Dar15, and the Dar15-bound “upper” chain. **c**, Combined reaction mixture of Dar12, a Dar6_N-bound “lower” chain, and an AsuC5- or Dar15-bound “upper” chain. LC–HRMS analysis showed signals at m/z [M + 10H]^10+^ 1110.2682 and 1110.2684, consistent with the theoretical mass (1110.2618) of the Dar6_N-bound “lower” chain. No detectable AsuC5- or Dar15-bound “upper” chain or Dar6_N-bound amidative condensation product (m/z [M + 10H]^10+^ 1393.3187, theoretical) was observed.

**Extended Data Fig. 4.**
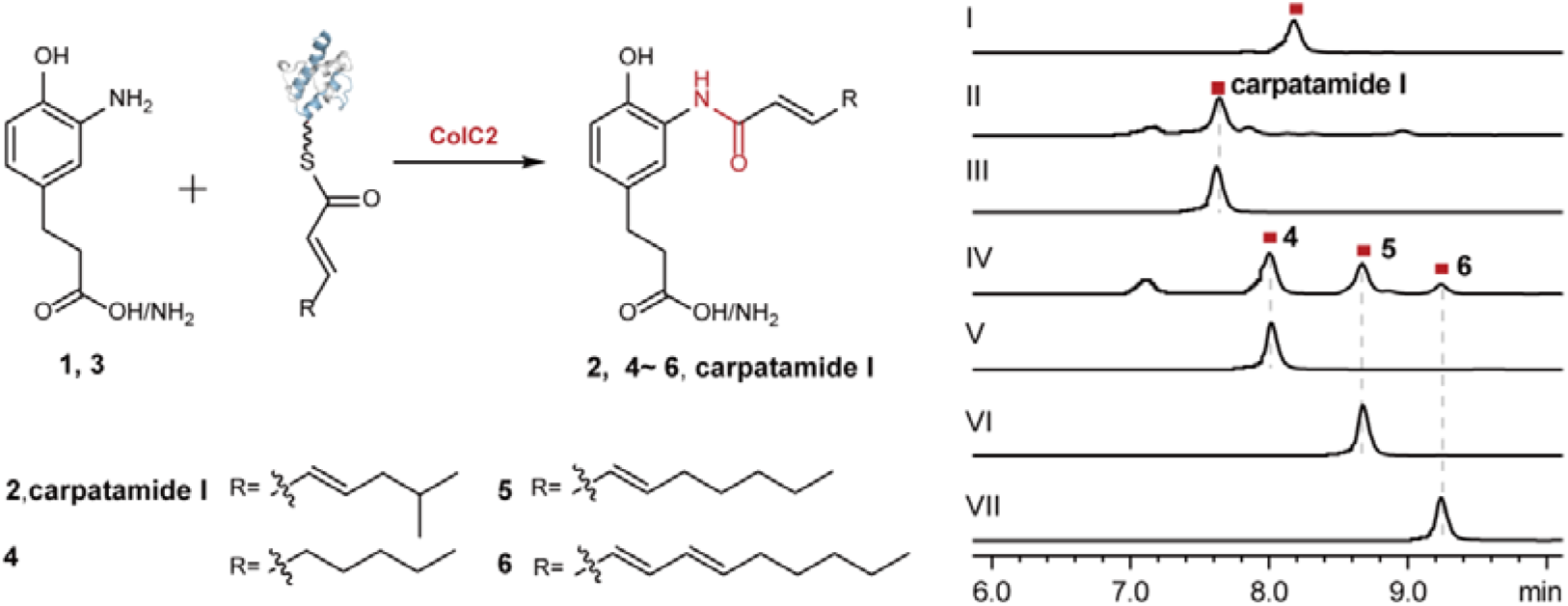
UPLC analysis of the *in vitro* enzymatic reactions of ColC2. (I) Compound **2** was biosynthesized by combining **1** with reaction mixture B (isovaleric acid, HSCoA, ATP, UkaQ^FAV^, malonic acid, *Sc*MatB, *Sa*FabD, AsuC3, AsuC5, AsuC7, AsuC8/C9) and ColC2. (II) Carpatamide I was biosynthesized by combining **3**, reaction mixture B, and ColC2. (III) Carpatamide I standard. (IV) Compounds **4**, **5**, and **6** were biosynthesized by combining **1** with reaction mixture B (where isovaleric acid was replaced with hexanoic acid) and ColC2. (V-VII) Standards **4**, **5**, and **6**. All reactions were detected at λ = 254 nm.

**Extended Data Fig. 5.**
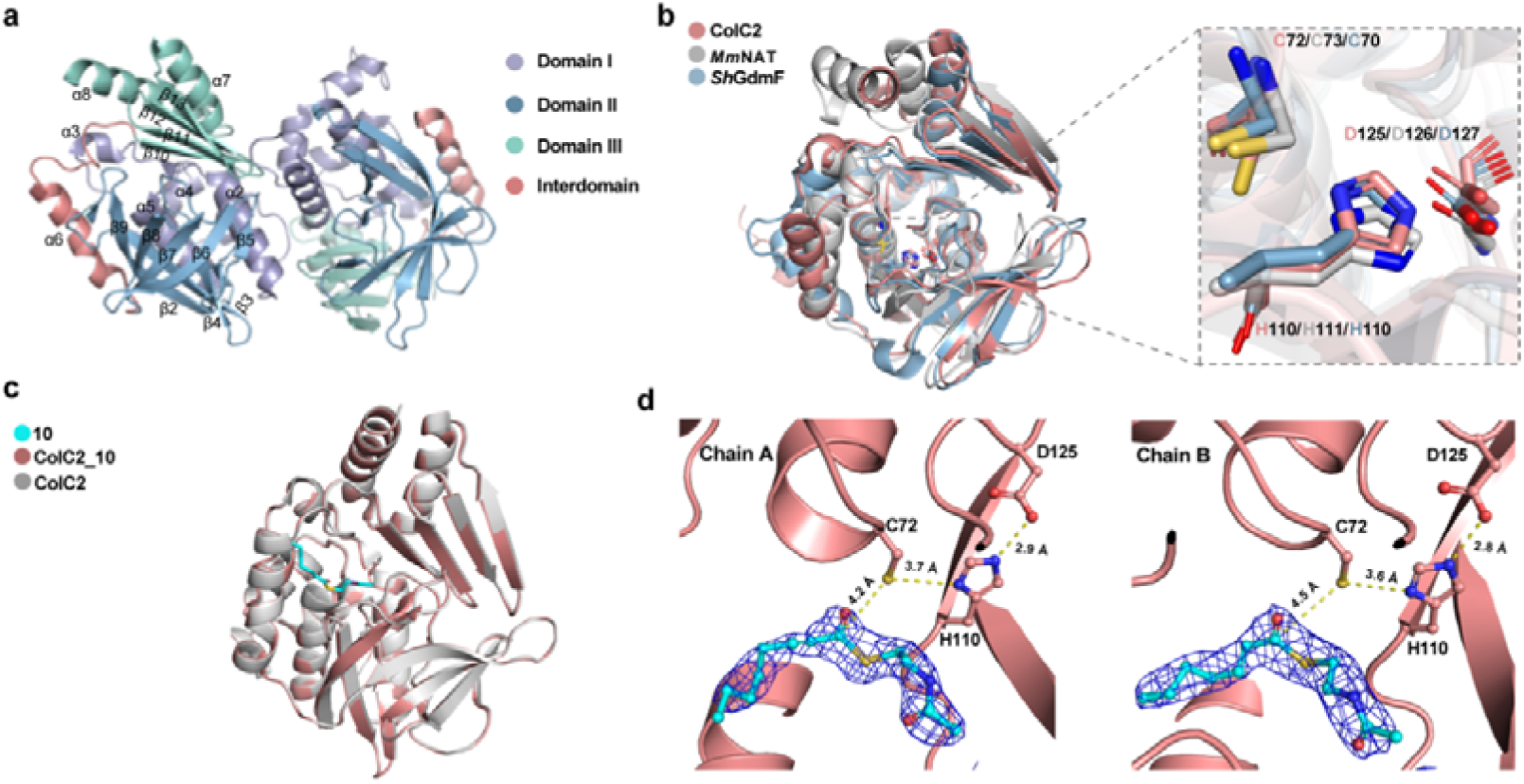
Structural analysis of ColC2. **a**, Overall structure of ligand-free ColC2 (PDB ID: 8K51) in a single cell. **b**, Structure alignment of ligand-free ColC2 (PDB ID: 8K51), *Sh*GdmF (PDB ID: 8OOM), and *Mm*NAT (PDB ID: 2VFC). The conserved catalytic triads are highlighted. **c**, Structure alignment of ligand-free ColC2 (PDB ID: 8K51) and the ColC2_**10** complex (PDB ID: 8K56). **d**, Binding conformation of **10** in Chain A and Chain B of the ColC2 complex structure. The *2Fo*-*Fc* electron density map of **10** is contoured at a 1.0 σ level. The catalytic residues of ColC2 are shown as ball-and-stick models.

**Extended Data Fig. 6.**
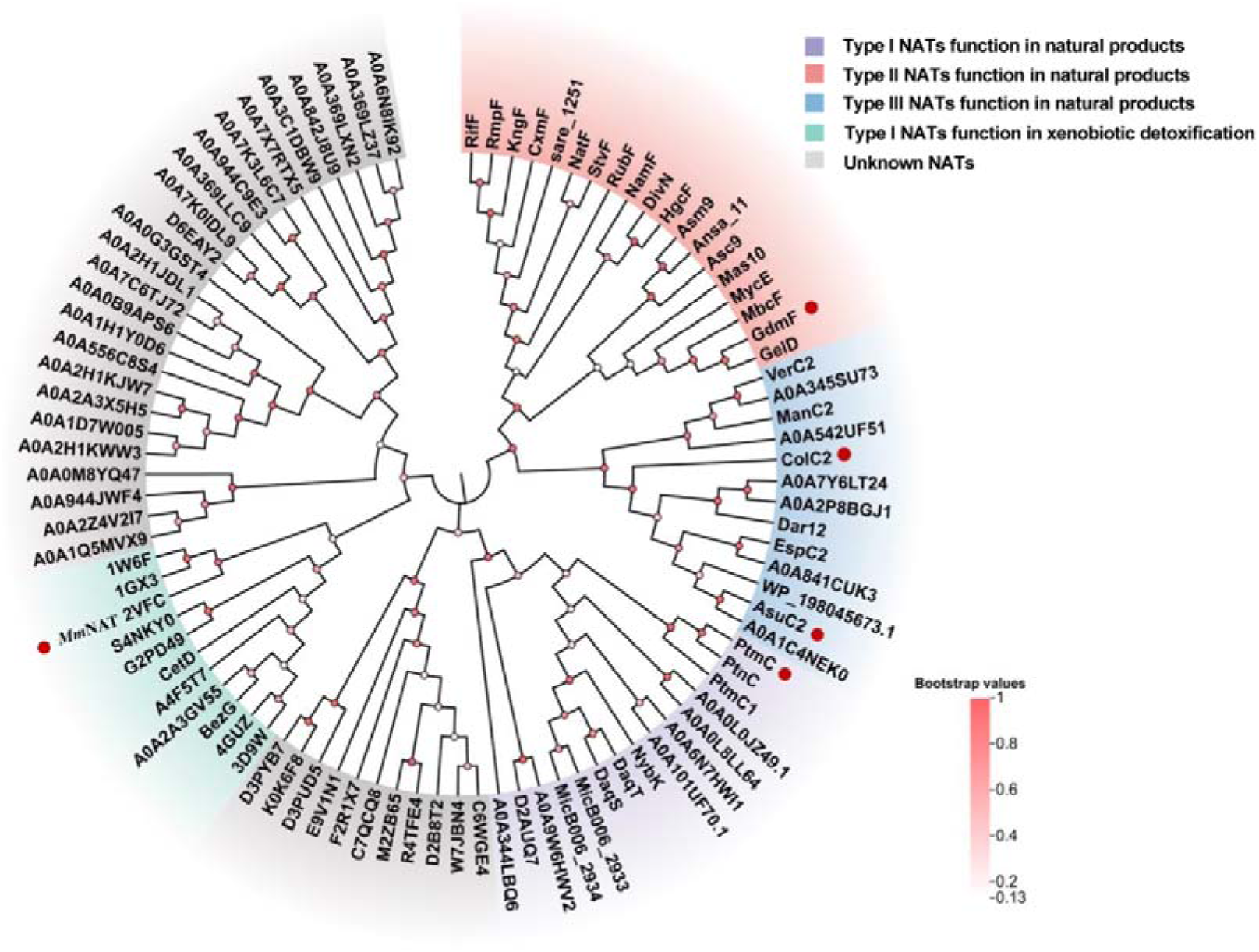
Phylogenetic relationships of arylamine NATs. The tree was constructed using the Neighbor-Joining method. Color coding was performed in iTOL. Representative NATs are marked with red circles.

**Extended Data Fig. 7.**
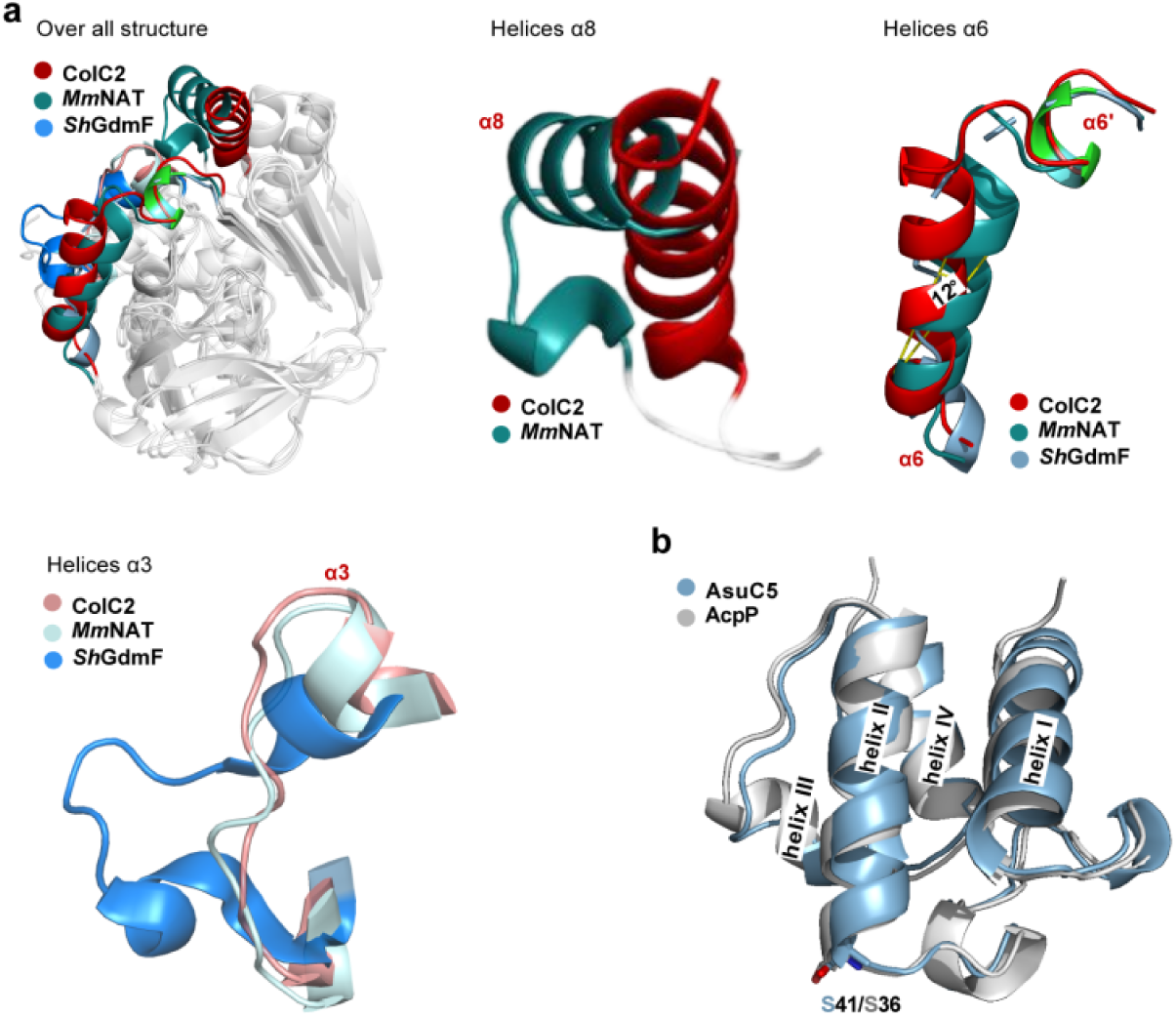
Architecture differences between ColC2, *Sh*GdmF, and *Mm*NAT. **a**, Alignment of the overall structure and an enlarged view of the architectural differences between ColC2 (PDB ID: 8K56, highlighted in red), *Sh*GdmF (PDB ID: 8OOM, highlighted in blue), and *Mm*NAT (PDB ID: 2VFC, highlighted in deep teal) in helices α8, α6, and α3. **b**, Structural alignment of AcpP (PDB ID: 7L4E) and AsuC5. The structural model of AsuC5 was predicted using AlphaFold2.

**Extended Data Fig. 8.**
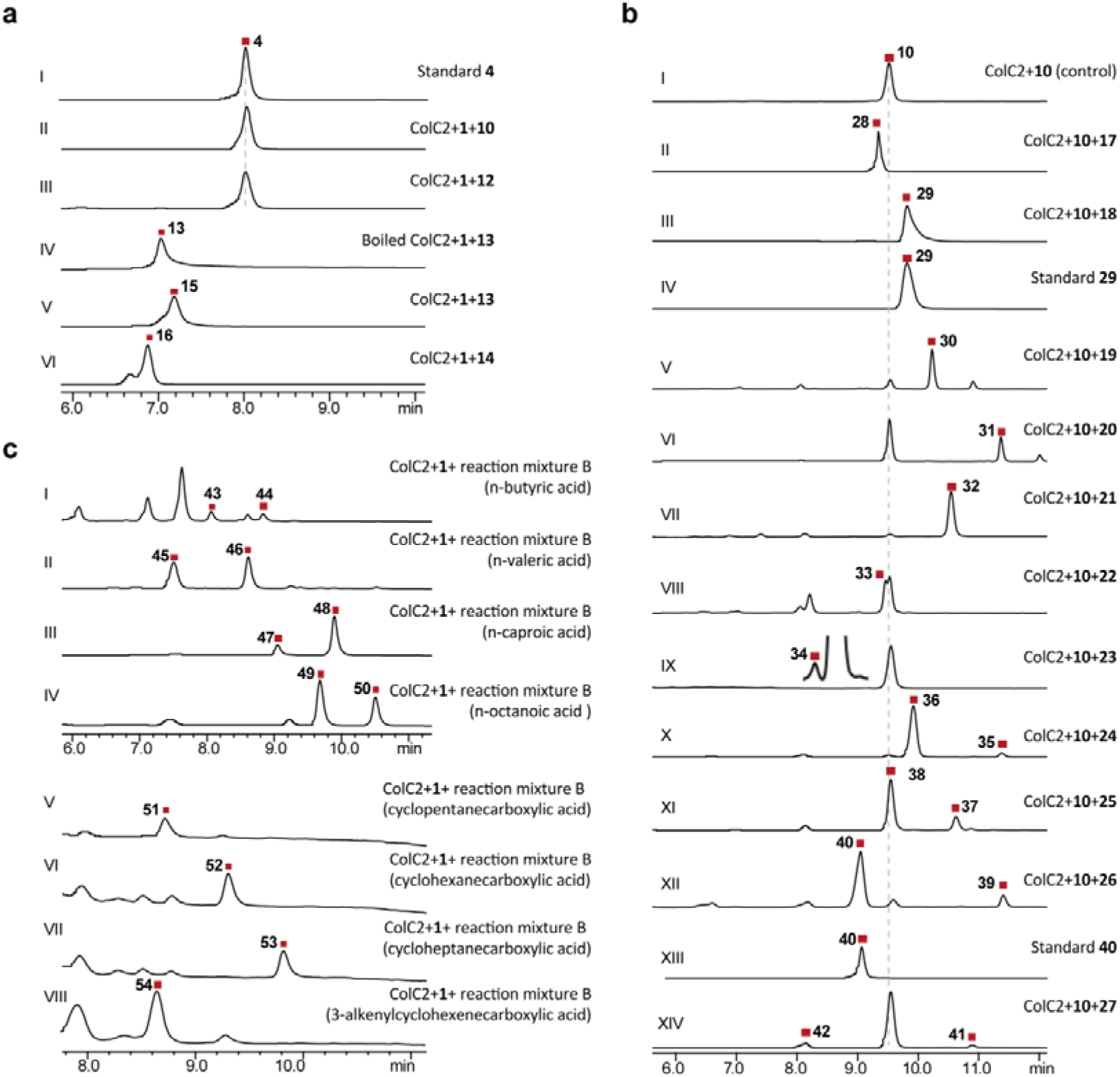
Substrate scope of ColC2. **a**, Evaluation of acyl donor promiscuity. Reactions were conducted with ColC2, substrate **1**, and various acyl donors, as indicated in each trace. **b**, Evaluation of acyl acceptor promiscuity. Reactions were conducted with ColC2, substrate **10**, and different acyl acceptors, as indicated in each trace. Acyl acceptors include 3,4-AHBA (**17**), 2-amino-4-methylphenol (**18**), 2-amino-6-methylphenol (**19**), 2-amino-3-methylphenol (**20**), 2-aminophenol (**21**), 3-aminophenol (**22**), 4-aminophenol (**23**), m-toluidine (**24**), o-toluidine (**25**), p-toluidine (**26**), and aniline (**27**). **c**, Convergent amidative condensation between aryamine substrate **1** and the polyketide upper chain synthesized using the KAIII-PKS system with various starter units. Reactions were conducted with ColC2, substrate **1**, and reaction mixture B, where isovaleric acid was replaced with the indicated acids in each trace. Note that UkaQ^FAV^ was substituted with AliA_100_ in reactions (V-VIII). All reactions were detected at λ = 254 nm.

**Extended Data Fig. 9.**
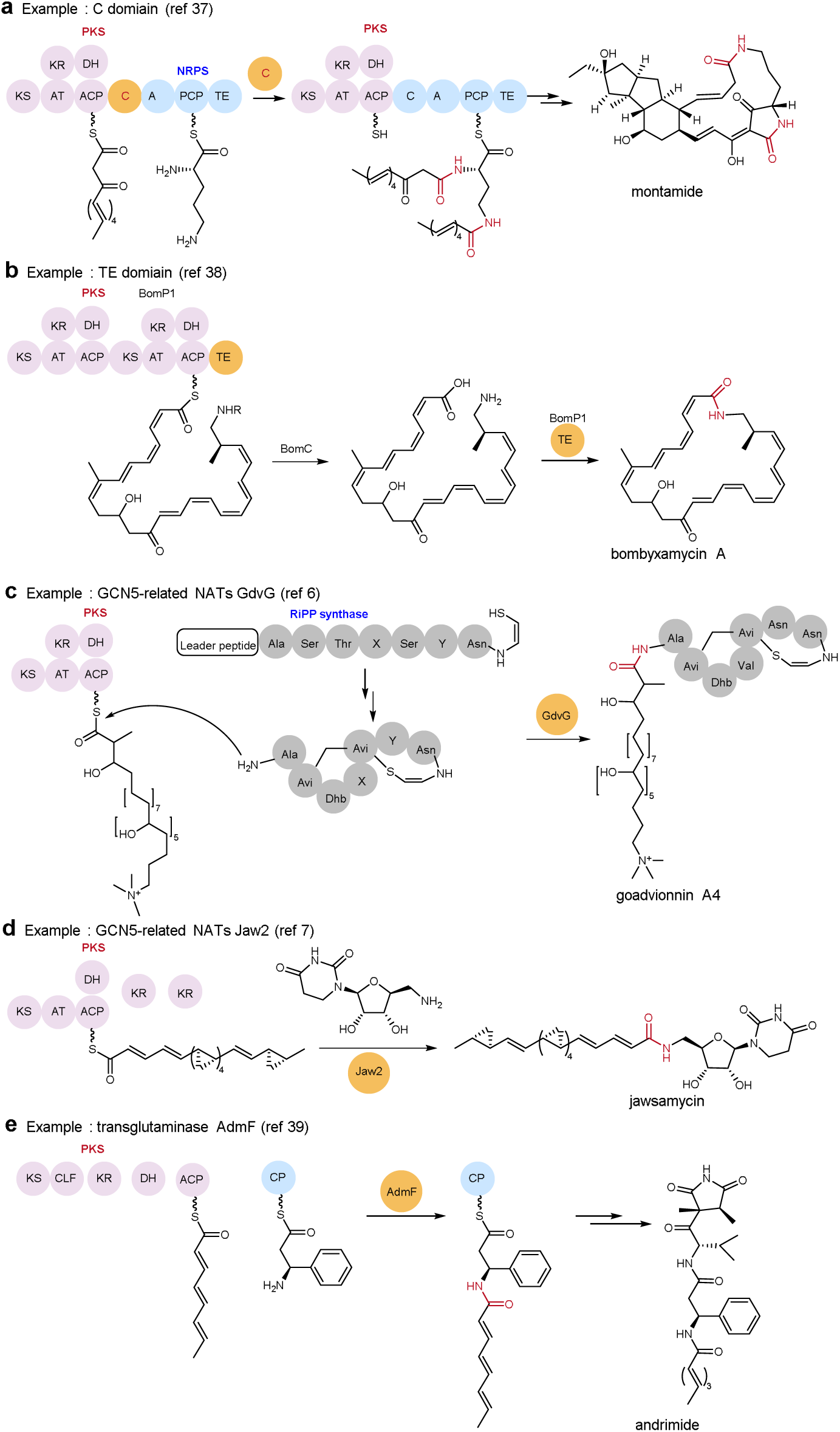
Enzymes responsible for amidative condensation. **a**, NRPS-C domain in montamide biosynthesis. **b**, TE domain in bombyxamycin A biosynthesis. **c**, GdvG in goadvionnin biosynthesis. **d**, Jaw2 from the jawsamycin biosynthesis. **e**, AdmF in andrimide biosynthesis.

## Notes

### Competing Interest Statement

The authors have declared no competing interest.

