## Supporting figure for "A Divergent Class of Arylamine N-Acetyltransferases Catalyzes Convergent Amidative Condensation of Polyketides in Manumycins Biosynthesis"

**Table of Contents**

1. Gerneral Information

1.1 Materials

1.2 Sequence of optimized Dar12, Dar15, Dar6_N, ColC2 and Dar24

1.3 Construction of the expression plasmids

1.4 Site-directed mutagenesis

1.5 Chemical synthesis

2. Supplementary Tables

Table 1. Protein-protein interaction (PPI) analysis of ColC2 and AsuC5

Table 2. *X*-ray data collection and structure refinement statistics for ColC2

Table 3. Bacterial strains and plasmids used in this study

Table 4. Primers used in this study

Table 5. Accession number of the protein sequences used for the phylogenetic analysis and sequence alignment

3. Supplementary Figures

Fig. 1. Structural alignment and comparative modeling of ACPs

Fig. 2. UPLC-MS analysis of feeding assays with compound **1**

Fig. 3. Sequence alignment of ColC2 with representative arylamine NATs

Fig. 4. MM/PBSA binding energy analysis of the ColC2–AsuC5 complex

Fig. 5. Substrate specificity and biocatalytic scope of Dar12

Fig. 6. HR-ESI-MS analysis of synthetic and biosynthetic polyketide amides

Fig. 7. Molecular dynamics simulations of the ColC2–AsuC5 complex

Fig. 8. SDS-PAGE analysis of purified recombinant enzymes and mutants

Fig. 9. ^1^H (500 MHz) NMR spectra of **Int-1** in CDCl_3_

Fig. 10. ^13^C (126 MHz) NMR spectra of **Int-1** in CDCl_3_

Fig. 11. ^1^H (500 MHz) NMR spectra of compound **3** in D_2_O

Fig. 12. ^13^C (126 MHz) NMR spectra of compound **3** in D_2_O

Fig. 13. ^1^H (500 MHz) NMR spectra of compound **4** in CD_3_CN

Fig. 14. ^13^C (126 MHz) NMR spectra of compound **4** in CD_3_CN

Fig. 15. ^1^H NMR (500 MHz) spectra of (2*E*,4*E*)-deca-2,4-dienoic acid in CDCl_3_

Fig. 16. ^1^H (500 MHz) NMR spectra of compound **5** in acetone-*d_6_*

Fig. 17. ^13^C (126 MHz) NMR spectra of compound **5** in acetone-*d_6_*

Fig. 18. ^1^H NMR (500 MHz) spectra of (2*E*,4*E*,6*E*)-dodeca-2,4,6-trienoic acid in CDCl_3_

Fig. 19. ^1^H (500 MHz) NMR spectra of compound **6** in acetone-*d_6_*

Fig. 20. ^13^C (126 MHz) NMR spectra of compound **6** in acetone-*d_6_*

Fig. 21. ^1^H (500 MHz) NMR spectra of compound **7** in acetone-*d_6_*

Fig. 22. ^13^C (126 MHz) NMR spectra of compound **7** in acetone-*d_6_*

Fig. 23. ^1^H (600M) NMR spectra of compound **10** in CDCl_3_

Fig. 24. ^1^H (500M) NMR spectra of compound **12** in D_2_O

Fig. 25. ^1^H (600M) NMR spectra of compound **13** in CDCl_3_

Fig. 26. ^1^H (500 MHz) NMR spectra of compound **29** in CDCl_3_

Fig. 27. ^13^C (126 MHz) NMR spectra of compound **29** in CDCl_3_

Fig. 28. ^1^H (500 MHz) NMR spectra of compound **40** in CD_3_CN

Fig. 29. ^13^C (126 MHz) NMR spectra of compound **40** in CD_3_CN

4. Supplementary References

**1. General Information**

**1.1 Materials**

Strains, plasmids, and polymerase chain reaction (PCR) primers used in this study were listed in Tables 3 and 4. DNA of Dar12, Dar15, Dar6_N, ColC2 and Dar24 were synthesized and codon-optimized at Jiutian Gene Technology Co., Ltd. (Tianjing, China). Plasmid extraction, DNA purification and gel extraction were carried out using commercial kits (Sangon Biotech Co., Ltd., Shanghai, China). Primers synthesis and DNA sequencing were performed at Tsingke Biotech Co., Ltd. (Shanghai, China). *Escherichia coli* strains DH5α and BL21(DE3) were used as hosts for DNA manipulation and recombinant protein expression, respectively. *Streptomyces* *lividans* TK24 was used as hosts for expression of *dar24* and feeding assay. LB and tryptic soy broth (TSB) broth were used as media for cultivating *E. coli* and *S.* *lividans* TK24. Restriction enzymes, DNA polymerases (Taq and PrimeSTAR), DNA maker, and Protein maker were purchased from New England Biolabs Co., Ltd. (Beijing, China) and Takara Biotechnology Co., Ltd. (China). All chemicals and reagents were purchased from commercial sources unless noted otherwise. ESI-APCI-MS were measured on a Shimadzu LCMS-2020 coupled with a Shimadzu LC-40D XS UPLC equipped with a Shim-pack Scepter HD-C18 column (2.1 × 50 mm, 1.9 μm, Shimadzu). HR-ESI-MS were measured on Thermo Scientific Orbitrap Exploris 120 equipped with a UMISIL™ C18 column (4.6 × 250 mm, 5 μm, Thermo Scientific). Semipreparative HPLC was performed on Shimadzu LC20-AT with SPM-20A detector, equipped with an Agelar Technologies Venusil MP C18(2) column (250 × 10 mm). NMR experiments were conducted on a Bruker AVANCE 500/600 spectrometer at 500/600 MHz for ^1^H and 126 MHz for ^13^C.

**1.2 Sequence of optimized Dar12, Dar15, Dar6_N, ColC2 and Dar24**

In order to express in *E. coli* and *S. lividans* TK24, Dar12, Dar15, Dar6_N, ColC2 and Dar24 genes were codon optimized and synthesized by Jiutian Gene. Sequence see below. Restriction sites for cloning are underlined.

***dar12*:** cat**ATG**ACCGCCCGTCTGCATCATCCGACCGCCGCCGGTACCCGTTGGGAACTGCCGACCGCCGAATATCTGCGTCGCCTGGGCTATAGTGGTCCGGTTGAACCGGGCGCAGATTGTCTGCGTGCCCTGTGCGAAGCCCATCTGCTGGCCGTGCCGTATGAAATGCTGGATGCCCTGGAAGGTCGCCGCCCGTGTCTGGAACTGCCGGGTGTGTTTGATAAACTGGTGTATCGTCGCTGCGGCAGCACCTGCCTGGAAGCAACCCCGCTGTTTGGTCGTTTTCTGCGTGATGTTGGCTTTCGTGTTCGTCTGGTGGCAGCCCAGGCATGGCGTGTGAATGGTCGCTGGGCCCCGCGCTGGGATCATCTGCTGCTGCTGGTTGAAGCCGAAGGTGCAGAATGGGTGGTGGATGTGAGTTTTCTGATGCTGACCGTTCTGCGTCCGCTGCGTACCGGCGGTGAACCGCAGGAACATAGTGGCTGGACCCATCGCGTTGGTGAAGTTGATGGCCATCGCGCCGTGCTGCGCCGTATGCCTGGTGGTGATTGGGTTCCGGTTTATCGTTTTGCCGAACGTGCCCTGGAAATTGATGATTATGCCTGGATTGTGGATTATCACATGGATGCCGATGATAGCCCGCTGACCGGTAGTCTGCTGTGTAGCCGCGTTGTGCCGGGTGGCAAACTGATTGTGATGCGTGATAATTTTGTTCGTGCAGAAGATGGTCGTGAAAGCGTTGATTTTCTGGCCACCGCAGCCGAAGCAGAAAAAGCCTTTGCCGAAGTTTTTCAGGATCATGGTCATCTGGTGGAAAAAGCACTGGGCCTGTGGGAAAAAACCCGTCGCAGCCATCGTGGCCCGCTGCCTGGTATTGGT**Taa**gct

***dar15*:** cat**ATG**AGCTCTCGTGACGACATCTTGATCACTCTGCGTGAAATTCTGGACGAAGTTGCGGAAATTCCGGGTGAACAGGTTACCTTGGACGCATCTTTCGCAGGCGATCTGGAACTGGATAGCCTGGCGGTGGTTTCCGTGTTCGTTCTGGTTCAGCGTCGTTTCGGCATCGAAGTTCCGAACGAAGATGCTGACCGTCTGACCACCGTACGTGAAGCTGTTGACTACCTGGAGAAACGTTCCGCGGCGGTG**TAA**gaattc

***dar6_N*:** cat**ATG**CGTACCGCAGAAACCGATCGTATTACCGAATTTGTTCTGAAAGTGGTGCGTGATATGCTGAATGCCCCGCTGCCGCCGAGTACCCCGCCTGGTATTCCGCTGGGCCCGGGTGGTCTGGAACTGGAAAGTCTGAGCCTGGTGGAACTGACCGTTCATGTTGAACGTGAATATGGTATTCGTTTTCCGGATGAAGCAGTTGATGGCCTGGGCACCATGACCCTGGGCGAACTGGTGGCAGATATTCGTAGTCGTGCACCGGGCGAAGGCGGCACCGGTGAACCTCCTACC**TGA**gaattc

***colC2***: cat**ATG**GAAACCGAAAGCCTGCCGTGGCGCGAATATCTGGAACGTATTGGCTATCAGGGCCTGCTGAATAATAGTCTGGAATGTCTGCGTGAACTGTATACCGCCCATCTGCGTAGTGTGCCGTATGAAATGCTGGATAGTTTTGATGGCACCCCGCCGGTTCTGGGCCATGCCGAAAGTTTTGCCAAACTGGTGCATCGTCGCCGCGGCGGCAATTGTCTGGAAAGCACCCCGCTGTTTGGTGAATTTCTGCGTCAGGCAGGCTTTGAAGTTCGCCTGGTGCCGGCACAGATTTGGAAAGTTAGCGGTGAATGGTGGGATGCATGGGATCATCTGCTGCTGATTGTTACCGTGGATGGTGAAGATTGGCTGCTGGATGTTGGCTTTCTGATGCTGACCTTTGCAGAACCGCTGAAAGTGGCCGAAGGCCCGCAGGAACAGAGCGGTTGGCGCTTTCGCGTTGCAGAAGAAGAAGGCTTTCCGACCGTGAGCCATCAGGGCCCGGATGGTACCTGGACCGCAGTTTATCGTTATCGCGATGAACCGCAGCAGCGTGCAGATTATGAATGGATTATTGATTTTCACAAGAGCGCCGAAGATAGTCCGCTGGTGGGTACCCTGCTGTGCAGTCGCAATGTTCCGGATGGCAAACTGATTATGATTGGCGAAAATCTGCTGCATGCACGTAATGGCCGTGTTAGCGCAGAATTTATTGAAACCACCAGTCGCGCCGAAGAACTGCTGCGTGTGATTTTTGCAGGCCATGAACACATGGTGGAAAGTGCCGTTCGCACCTGGGAAAAAGCACGTGCAGATCGCAGTACCCGTCGCAGCCTGGTTTATCGTAAAGCAGAACAG**Taa**gct

***dar24***: cat**ATG**TGCGGTATAACCGGATGGGTTTCGTTCGACCGTGAGCTGACAGCCCGCCAGGACGTCCTGGACGCCATGACGGCGACCATGTCCTGTCGCGGACCGGACGCCAGCGGCGTCTGGACCTCGCGCCACGCGGCGGTCGGCCACAGCCGGCTGGCGATCATCGACCTGGAGGGCGGGCGCCAGCCCATGTCCACGGCGGTGGGCGGCGCGGAGATCGTCATCACCTACAGCGGCGAGGTCTACAACTTCCAAGAACTGCGTGGGTCCCTGACGGCCAAGGGGCACGAATTCACGACCCGTAGCGACACGGAGGTGGTGCTGCGCGCATACATCGAGTGGGGCGATGAATTCGTCGAAAAGCTGAACGGCATGTTCGCCTTCGCCGTCTGGGACTCCCGCTCGGAGCGGATGCTCCTGGTGCGCGACCGCGCCGGCGTGAAGCCCCTTTTCTACCAGCCCCTGCCCGACGGTGTGCTGTTCGGTTCCGAGCCGAAGGCCGTCCTCGCCCATCCGGAGGGGGACCGCAGGGTCGACGGTGAGGGGCTGCGGGAGATGTTCTCCTTCACACGCACCCCGGGACACGCGGTGTGGAGAGGGATGCGGGAAGTGAGACCGGGCACCGTGGTCGCCGTGGACCGCAACGGTGTTCGCGAGCGCACGTACTGGCGTCTGGAAGCGCGGGCGCACACCGACGGCCTGACCGCGACGACGACACGGATCCGCGAGCTGCTGGAGGACATCGCGGCCCGGCAGATGGTCTCGGACGTCCCTCGCTGCGTGCTGCTCTCCGGAGGCCTCGACTCAAGCGCCCTGACGGCTCTGGCCGCAGCCGGACTCCGCAGAGGGAGAGACGACTCGCCGGTGCGCACGTTCGCCGTCGACTTCGCGAACGGTGCCGAGACCTTCCGGGCAACCCGGCTGCGTGAAACCCCCGACGCCCCCTACATCCGGGATGTGGTCACCCACGTCGGCACCGATCACCACGACATCGTGCTCGACTCCCGCGACATGGCGGCACCGCAGGTGAGACGCCGCACCGTGAGAGCGCGGGATCTCCCCGGGGTCCACGGTGACATGGACCTCTCTCTGCACCTGCTCTTCCGTGCGGTCAGCCGGGAGTCGACGGTCGCGCTGTCCGGCGAGGCGGCCGACGAACTGTTCGGCGGGTACCTGTGGTTCCACGACCCGGCCGCCCAGCGAGCTGAGACTTTCCCCTGGACGGCCGTCTTCGGGGACGCCGAGCAGGAGGCCCTCAAGCTGCTGCGCCCTGAGGTCCGGCAGCGTCTGGGACTGGCCGAGTACGCGAGACAGCGGTACTCCGAGGCCGTCGCCGAGGTCCCCAGACTGCCGGGCGAGGAGGGTCTGGAGCTGCGGATGCGGCGCATCGGCTATCTCAACCTCACCCGGTGGATGCCCTCCCTGCTGGAGCGGAAGGACCGGATGAGCATGGCCACGGGTCTGGAGGTCCGGGTACCGTTCTGCGACCACCGCCTCACCGAGTACGTCTTCAACACCCCCTGGAGCATGAAGACGTTCGACGGCCGGGAGAAGAGCCTGTTGCGGGCAGCCGTACGGGATCTGCTTCCGGCGTCGGTGGCCGACCGCAGGAAGAGCCCCTACCCCTCGACCCAGGACCCTTTCTACCAGCAGGAGTTGCAGCAGCAGGCCAAGCAGGTGCTGGCCGACGCGGATGACACGCTGTTCGGCCTGGTCAGCAGGGAATGGCTGGCCGGGGCCGTCGGTGGCGATCCGGGGACCATGAGCATCTCCGTGCGGCACGGCCTGGAGCGGGTGCTGGAGTTGTCGGCGTGGATCGATCTGTACAGTCCGGACATCACCCTGGCC**TGA**gaattc

**1.3 Construction of the expression plasmids**

For Dar12 and ColC2, codon-optimized genes were synthesized and used directly in plasmids pJTU117 and pJTU118, respectively (Table 3). For Dar15 and Dar6_N, synthesized genes served as templates for PCR amplification using the primer pairs Dar15-pET28a-F/R and Dar6-pET28a-F/R. The resulting fragments were cloned into the *Nde*I–*Eco*RI sites of pET28a with an N-terminal MBP tag and a TEV protease cleavage site from pSJ8^1^ to generate plasmids pJTU119 and pJTU120. For Dar24, the *dar24* gene was amplified using primers Dar24-pWY45-F/R and cloned into the *Nde*I–*Eco*RI sites of pWY45^2^, yielding plasmid pJTU121. Plasmids for AsuC3, AsuC5, AsuC7, AsuC89, AliA_100_, Sfp, *Sc*MatB, and *Sa*FabD (pJTU37, pJTU40, pJTU42, and pJTU44–pJTU47) were obtained from previous studies^3^.

**1.4 Site-directed mutagenesis**

Single missense mutations were introduced into pJTU118 (pET28a-ColC2) and pJTU40 (pET21a-AsuC5) using a PCR-based site-directed mutagenesis protocol. For each mutation, a forward primer was designed to incorporate the desired nucleotide substitutions, flanked by an approximately 30-nucleotide sequence downstream of the target site. The corresponding reverse primer was designed to bind to an approximately 30-nucleotide sequence immediately upstream of the mutation site (Table 4). Whole-plasmid PCR amplification was performed using these mutation-specific primer pairs with pJTU118 or pJTU40 as the template. Following amplification, the parental (methylated) template DNA was digested with *Dpn*I (New England Biolabs). The resulting nicked circular mutant plasmids were transformed into *E. coli* DH5α. All mutant plasmids (pJTU122–pJTU141) were verified by DNA sequencing (Tsingke, China) and subsequently transformed into *E. coli* BL21(DE3) for protein expression.

**1.5 Chemical Synthesis**

Synthesis of 3-(3-amino-4-hydroxyphenyl)-propanamide (**3**)

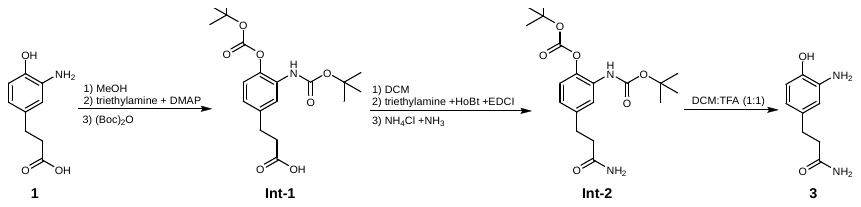

To a solution of 3-(3-amino-4-hydroxyphenyl)-propanoic acid (**1**) (36 mg, 1 eq.) in dry MeOH (5 mL), was added (Boc)_2_O (4 eq.), triethylamine (3 eq.), DMAP (0.1 eq.) at ambient temperature. After 10 h, the reaction mixture was evaporated *in vacuo* and then purified by HPLC using an Agelar Technologies Venusil MP C18(2) column (250 × 10 mm) on Shimadzu LC20-AT with SPD (SPM-20A) detector, with mobile phase B (100% MeCN): A (100% H_2_O containing 0.1% FA) 67: 33 within 0-15 min, at a flow rate of 2 mL min^-1^ to obtain (3-(tert-butoxycarbonyl)amino)-4-(tert-butoxycarbonyl)hydroxy)phenyl) propanoic acid (**Int-1**, 27 mg, yield 35%). ^1^H NMR (500 MHz, CDCl_3_) δ 7.09 (d, *J* = 8.3 Hz, 1H), 6.88 (dd, *J* = 8.4, 2.1 Hz, 1H), 6.72 (s, 1H), 2.93 (t, *J* = 7.9 Hz, 2H), 2.67 (t, *J* = 7.9 Hz, 2H), 1.56 (s, 9H), 1.52 (s, 9H). ^13^C NMR (126 MHz, CDCl_3_) δ 178.33, 152.56, 151.12, 138.52, 138.42, 130.25, 122.76, 121.56, 120.05, 84.41, 80.97, 35.55, 30.42, 28.31, 27.63. MS (ESI-APCI) *m/z* [M + H]^+^ calcd. for C_19_H_28_NO_7_ 382.19, found 382.30.

To a solution of **Int-1** (27 mg) in anhydrous DCM (3 mL) was added triethylamine (4 eq.), HoBt (1.2 eq.), EDCI (2 eq.). The reaction mixture was stirred at ambient temperature for 1 h, then NH_4_Cl (4 eq.) solid and NH_3_ solution (7 N in methanol, 5 eq.) were dropwise added. The reaction mixture was stirred for 5 h and then concentrated *in vacuo*, the residue was dissolved in ethyl acetate (5 mL), washed with brine (2 × 2 mL), dried over anhydrous Na_2_SO_4_ and evaporated *in vacuo.* The residue was then dissolved in 0.6 mL DCM: TFA (1:1). The reaction mixture was stirred at ambient temperature for 1 h and then concentrated *in vacuo*. The residue was purified by HPLC using a YMC-Triart C18 column (250 × 10 mm) on Shimadzu LC20-AT with SPD (SPM-20A) detector, with mobile phase B (100% MeCN): A (100% H_2_O containing 0.1% FA) 0%-16% B (0-20 min), at a flow rate of 2 mL min^-1^. 3-(3-amino-4-hydroxyphenyl)-propanamide (**3**), brown solid. ^1^H NMR (500 MHz, D_2_O) δ 6.70 (d, *J* = 8.1 Hz, 1H), 6.66 (d, *J* = 2.1 Hz, 1H), 6.54 (dd, *J* = 8.1, 2.1 Hz, 1H), 2.70 (t, *J* = 7.4 Hz, 2H), 2.43 (t, *J* = 7.4 Hz, 2H). ^13^C NMR (126 MHz, D_2_O) δ 179.07, 142.72, 134.46, 133.27, 119.63, 117.22, 115.60, 36.91, 30.43. MS (ESI-APCI) *m/z* [M + H]^+^ calcd. for C_9_H_13_N_2_O_2_ 181.10, found 181.30.

Synthesis of compound **4**

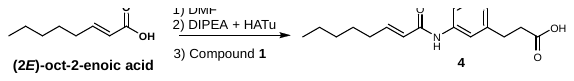

Synthetic procedures were referred to reported method with slight changes^4^. Briefly, (2*E*)-oct-2-enoic acid (0.1 mmol, 1 eq.) was dissolved in DMF (500 µL) and the solution was cooled to 0 °C, then DIPEA (0.2 mmol, 2 eq.) and HATu (0.105 mmol, 1.05 eq.) were added in order. The mixture was stirred at 0 °C for 5 min, and then **1** (0.12 mmol, 1.2 eq.) was added. The reaction was further stirred at 0 °C for 30 min. After the reaction, the solution was extracted by EtOAc. The organic layer was dried over *in vacuo* and compound **4** was purified by HPLC using an Agelar Technologies Venusil MP C18(2) column (250 × 10 mm) on Agilent 1260 infinity II with DAD HS detector and with mobile phase B (100% MeCN): A (100% H_2_O containing 0.1% FA) 70:30 within 0-15 min, at a flow rate of 2.5 mL min^-1^. Compound **4**, white solid. ^1^H NMR (500 MHz, CD_3_CN) δ 7.15 (d, *J* = 2.2 Hz, 1H), 7.03 – 6.94 (m, 2H), 6.85 (d, *J* = 8.3 Hz, 1H), 6.16 (dt, *J* = 15.2, 1.6 Hz, 1H), 2.81 (t, *J* = 7.6 Hz, 2H), 2.57 (t, *J* = 7.6 Hz, 2H), 2.27 (m, 2H), 1.50 (m, 2H), 1.39 – 1.27 (m, 4H), 0.93 (t, *J* = 6.9 Hz, 3H). ^13^C NMR (126 MHz, CD_3_CN) δ 174.39, 166.35, 148.18, 147.62, 133.26, 127.00, 126.50, 123.05, 122.84, 119.01, 117.95, 35.63, 32.26, 31.66, 30.06, 28.13, 22.74, 13.86. MS (ESI-APCI) *m/z* [M + H]^+^ calcd. for C_17_H_24_NO_4_ 306.17, found 306.30.

Synthesis of compound **5**

a) Synthesis of (*2E*,*4E*)-deca-2,4-dienoic acid

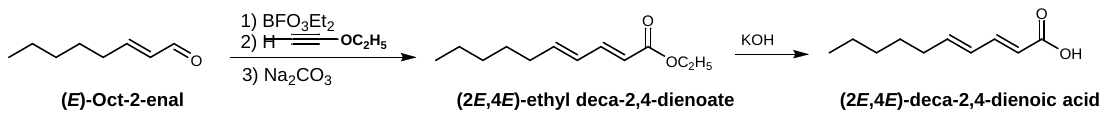

Synthetic procedures were referred to reported method^5^. (*E*)-Oct-2-enal (1 mmol, 1 eq.) was dissolved in ether (20 mL) under nitrogen, and the solution was cooled to -30 °C. A solution of boron trifluoride diethyl etherate (1.5 eq.) in 10 mL of ether cooled to -30 °C was added. The mixture was stirred at -30 °C for 30 min, and then cooled ethoxyacetylene (40% solution in hexane, 1.5 eq.) in 10 mL of ether was added slowly. The reaction was further stirred at -30 °C for 1.5 h. Then, a saturated aqueous sodium carbonate solution was added to quench the reaction, and the mixture was extracted with chloroform. The organic layer was dried over by anhydrous Na_2_SO_4_. After filtration, the solvent was removed *in vacuo*. The resulting (2*E*,4*E*)-ethyl deca-2,4-dienoate was used for the next step without further purification. (2*E*,4*E*)-ethyl deca-2,4-dienoate was dissolved in acetone (5 mL). After KOH solution (350 mM, 2 mL) was added, the solution was stirred at room temperature for 2 h and acidified by 6 M HCl, followed by EtOAc extraction. The organic layer was dried over *in vacuo*. the (2*E*,4*E*)-deca-2,4-dienoic acid was purified by HPLC using an Agelar Technologies Venusil MP C18(2) column (250 × 10 mm) on Shimadzu LC20-AT with SPD detector SPM20A, and with mobile phase B (100% MeCN): A (100% H_2_O containing 0.1% FA) 85:15 within 0-14 min, at a flow rate of 2.0 mL min^-1^. (2*E*,4*E*)-deca-2,4-dienoic acid, white solid. ^1^H NMR (500 MHz, CDCl_3_): δ 7.35 (m, 1H), 6.19 (m, 2H), 5.78 (d, *J* = 15.3 Hz, 1H), 2.18 (m, 2H), 1.44 (m, 2H), 1.31 (m, 4H), 0.89 (t, *J* = 6.9 Hz, 3H). MS (ESI-APCI) *m/z* [M + H]^+^ calcd. for C_10_H_17_O_2_ 169.24, found 169.30.

b) Synthesis of compound **5**

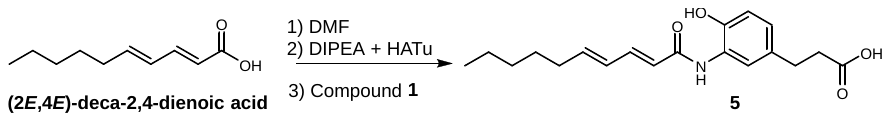

Compound **5** was synthesized in the same method as compound **4**. Compound **5**, light yellow solid. ^1^H NMR (500 MHz, Acetone-*d_6_*) δ 7.33 (dd, *J* = 14.9, 10.3 Hz, 1H), 7.25 (d, *J* = 2.1 Hz, 1H), 6.96 (dd, *J* = 8.2, 2.1 Hz, 1H), 6.83 (d, *J* = 8.2 Hz, 1H), 6.34 – 6.20 (m, 3H), 2.81 (t, *J* = 7.6 Hz, 2H), 2.56 (t, *J* = 7.6 Hz, 2H), 2.20 (m, 2H), 1.46 (m, 2H), 1.38 – 1.26 (m, 4H), 0.90 (t, *J* = 6.9 Hz, 3H). ^13^C NMR (126 MHz, Acetone-*d_6_*) δ 174.86, 167.46, 148.91, 145.98, 144.69, 134.25, 130.29, 128.15, 127.79, 123.70, 123.06, 120.15, 37.01, 34.48, 33.02, 31.55, 30.14, 24.03, 15.18. MS (ESI-APCI) *m/z* [M + H]^+^ calcd. for C_19_H_26_NO_4_ 332.19, found 332.30.

Synthesis of compound **6**

a) Synthesis of (2*E*,4*E*,6*E*)-dodeca-2,4,6-trienoic acid

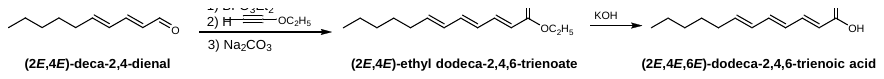

(2*E*,4*E*,6*E*)-dodeca-2,4,6-trienoic acid was synthesized in the same method as (2*E*,4*E*)-deca-2,4-dienoic acid. (2*E*,4*E*,6*E*)-dodeca-2,4,6-trienoic acid, white solid. ^1^H NMR (500 MHz, CDCl_3_) δ 7.39 (dd, *J* = 15.1, 11.4 Hz, 1H), 6.57 (dd, *J* = 14.8, 10.7 Hz, 1H), 6.24 (dd, *J* = 14.9, 11.3 Hz, 1H), 6.14 (m, 1H), 5.97 (dt, *J* = 14.7, 7.1 Hz, 1H), 5.84 (d, *J* = 15.2 Hz, 1H), 2.14 (m, 2H), 1.41 (m, 2H), 1.30 (m, 4H), 0.89 (t, *J* = 7.0 Hz, 3H). MS (ESI-APCI) *m/z* [M + H]^+^ calcd. for C_12_H_19_O_2_ 195.28, found 195.30.

b) Synthesis of compound **6**

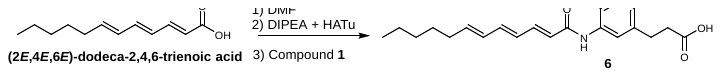

Compound **6** was synthesized in the same method as compound **4**. Compound **6**, yellow solid. ^1^H NMR (500 MHz, Acetone-*d_6_*) ^1^H NMR (500 MHz, Acetone-*d_6_*) δ 7.38 (dd, *J* = 14.8, 11.4, 1H), 7.26 (s, 1H), 6.96 (dd, *J* = 8.2, 2.2 Hz, 1H), 6.83 (d, *J* = 8.2 Hz, 1H), 6.69 (dd, *J* = 14.8, 11.4 1H), 6.37 (m, 2H), 6.24 (dd, *J* = 15.1, 10.7 Hz, 1H), 6.01 (m, 1H), 2.81 (t, *J* = 7.6 Hz, 2H), 2.56 (t, *J* = 7.6 Hz, 2H), 2.16 (m, 2H), 1.44 (m, 2H), 1.37~1.28 (m, 4H), 0.88 (t, *J* = 7.0 Hz, 3H). ^13^C NMR (126 MHz, Acetone-*d_6_*) δ 174.87, 167.32, 144.46, 142.80, 141.78, 134.23, 131.96, 129.82, 128.15, 127.78, 123.93, 123.75, 123.67, 120.12, 37.01, 34.45, 33.03, 31.55, 30.35, 24.04, 15.18. MS (ESI-APCI) *m/z* [M + H]^+^ calcd. for C_21_H_28_NO_4_ 358.20, found 358.30.

Synthesis of compound 7

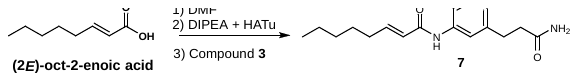

Compound **7** was synthesized in the same method as compound **4**. Compound **7**, white solid. ^1^H NMR (500 MHz, DMSO-*d_6_*) δ 9.41 (s, 1H), 7.56 (s, 1H), 7.27 (s, 1H), 6.83 – 6.75 (m, 3H), 6.73 (s, 1H), 6.33 (d, *J* = 15.3 Hz, 1H), 2.68 (t, *J* = 7.6 Hz, 2H), 2.29 (m, *J* = 7.6 Hz, 2H), 2.20 (q, *J* = 7.0 Hz, 2H), 1.45 (m, 2H), 1.30 (m, 4H), 0.89 (t, *J* = 6.9 Hz, 3H). ^13^C NMR (126 MHz, DMSO-*d_6_*) δ 173.47, 164.09, 146.10, 144.82, 131.94, 126.05, 124.55, 124.21, 122.08, 116.00, 37.08, 31.26, 30.79, 30.34, 27.36, 21.90, 13.86. MS (ESI-APCI) *m/z* [M + H]^+^ calcd. for C_17_H_25_N_2_O_3_ 305.19, found 305.30.

Synthesis of *trans*-2-octenoyl-SNAC (**10**)^5^

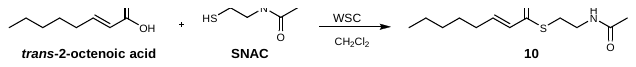

A mixture of *trans*-2-octanoic acid (10 mM, 1 eq.), DMAP (0.25 eq.) and WSC (2 eq.) was dissolved in dry CH_2_Cl_2_ (10 mL) under nitrogen gas and cooled to 0 °C. The mixture was stirred at 0 °C for 20 min. After *N*-acetylcysteamine (SNAC, 1.2 eq.) was added, the mixture was warmed to room temperature and stirred overnight. The reaction was quenched with water. The organic layer was washed with brine and dried over anhydrous Na_2_SO_4_. After filtration, the solvent was removed *in vacuo* and the compound was purified by HPLC using an Agelar Technologies Venusil MP C18(2) column (250 × 10 mm) on Shimadzu LC20-AT with SPD (SPM-20A) detector, with mobile phase B (100% MeCN): A (100% H_2_O containing 0.1% FA) 80:20 within 0-15 min, at a flow rate of 2 mL min^-1^. *Trans*-2-octenoyl-SNAC (**10**), white solid. ^1^H NMR (600 MHz, CDCl_3_) δ 6.85 (dt, *J* = 15.6, 6.9 Hz, 1H), 6.59 (brs, 1H), 6.05 (d, *J* = 15.6 Hz, 1H), 3.37 (q, *J* = 6.7 Hz, 2H), 3.02 (t, *J* = 6.7 Hz, 2H), 2.13 (m, 2H), 1.90 (s, 3H), 1.40 (p, *J* = 7.4 Hz, 2H), 1.24 (tp, *J* = 7.4, 4.3, 3.6 Hz, 4H), 0.82 (t, *J* = 6.9 Hz, 3H). MS (ESI-APCI) *m/z* [M + H]^+^ calcd. for C_12_H_22_NO_2_S 244.37, found 244.30.

Synthesis of *trans*-2-octanoyl-CoA (**12**)

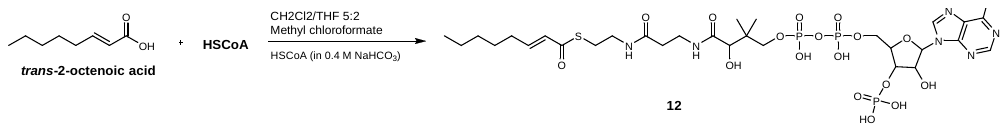

According to our previous chemical approach^3^, trimethylamine (3.0 μL, 30 μmol) was added to a solution of *trans*-2-octenoic acid (27 μmol) dissolved in the mixed solvent of CH_2_Cl_2_/THF (v: v= 5:2, 1.4 mL). The reaction system was stirred for 10 min at room temperature, then methyl chloroformate (2.9 μL, 30 μmol) was added and stirred for an additional 1 h at the same temperature. The solvents were evaporated under reduced pressure, and then 0.5 mL (CH_3_)_3_COH were added to the container with residues. Coenzyme A free acid (11.5 mg, 15 μmol dissolved in 0.25 mL of 0.4 M NaHCO_3_) was added to the solution, and the mixture was stirred for 0.5 h. After the reaction finished, 1 mL H_2_O was added and then subjected to HPLC for analysis and purification. Elution procedure for purification of *trans*-2-octanoyl-CoA (**12**) (90% yield) was performed on Shimadzu LC20-AT with SPM-20A detector, fitted with an Agelar Technologies Venusil MP C18(2) column (250 × 10 mm) with a linear gradient mobile phase B (100% MeCN) and A (100% H_2_O containing 10 mM ammonium acetate): 5%-60% B ( 0-10 min), 60%-5% B (10-13 min), 5% B (13-15 min) at a flow rate of 2.5 mL min^-1^. *trans*-2-octanoyl-CoA (**12**), white solid, ^1^H NMR (500 MHz, Deuterium Oxide) δ 8.34 (s, 1H), 8.02 (s, 1H), 6.71 (p, *J* = 8.3, 7.3 Hz, 1H), 6.00- 5.88 (m, 2H), 4.65 (m, 2H), 4.41 (s, 1H), 4.07 (s, 2H), 3.84 (s, 1H), 3.67 (m, 1H), 3.39 (m, 1H), 3.26 (s, 2H), 3.16 (s, 2H), 2.84 (d, *J* = 6.2 Hz, 2H), 2.24 (t, *J* = 6.4 Hz, 2H), 1.95 (t, *J* = 6.9 Hz, 2H), 1.29-1.15 (m, 2H), 1.03 (m, 4H), 0.71 (s, 3H), 0.63 (m, 3H), 0.57 (s, 3H). MS (ESI-APCI) *m/z* [M + H]^+^ calcd. for C_29_H_49_N_7_O_17_P_3_S 892.72, found 892.50.

Synthesis of 3-oxooctanoyl-SNAC (**13**)^5^

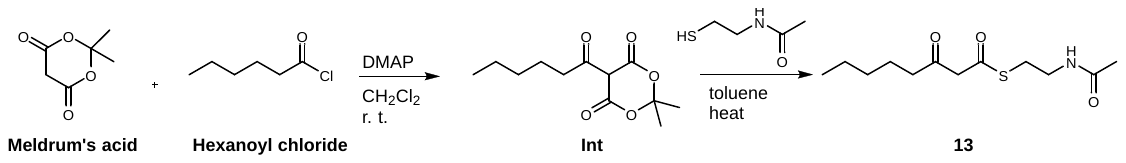

Under nitrogen gas, meldrum’s acid (20 mM, 1 eq.) and DMAP (2 eq.) were dissolved in dry CH_2_Cl_2_ (10 mL) and stirred at room temperature for 10 min. The reaction was then cooled to 0 °C and hexanoyl chloride (1 eq.) was added drop-wise. The mixture was warmed to room temperature and stirred overnight. The reaction was quenched with 1 M HCl. The organic layer was washed with brine and dried over anhydrous Na_2_SO_4_, filtered, and concentrated to yield a crude yellow oil, which was used for the next step without purification. This synthetic intermediate (20 mM) and SNAC (1 eq.) were dissolved in toluene (10 mL) and heated to 100 °C. The mixture was stirred overnight. The solvent was then removed *in vacuo* and the compound was purified by HPLC using an Agelar Technologies Venusil MP C18(2) column (250 × 10 mm) on Shimadzu LC20-AT with SPD (SPM-20A) detector, with a linear gradient mobile phase B (100% MeCN) and A (100% H_2_O containing 0.1% FA): 50%-100% B ( 0-12 min), 100%-50% B (12-15 min), 50% B (15-20 min) at a flow rate of 2.0 mL min^-1^. **3**-oxooctanoyl-SNAC (**13**), white solid. ^1^ H NMR (600 MHz, CDCl_3_): δ 6.79 (brs, 1H), 3.60 (s, 2H), 3.32 (m, 2H), 2.97 (td, *J* = 6.7, 4.4 Hz, 2H), 2.43 (t, *J* = 7.4 Hz, 2H), 1.87 (s, 3H), 1.47 (p, *J* = 7.4 Hz, 2H), 1.18 (m, 4H), 0.78 (t, *J* = 7.5 Hz, 3H). MS (ESI-APCI) *m/z* [M + H]^+^ calcd. for C_12_H_22_NO_3_S 260.37, found 260.30.

Synthesis of compound **29**

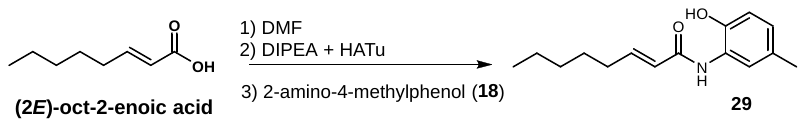

Compound **29** was synthesized in the same method as compound **4.** Compound **29**, white solid. ^1^H NMR (500 MHz, CDCl_3_) δ 8.41 (s, 1H), 6.98 (m, 2H), 6.86 (s, 2H), 6.04 (d, *J* = 15.2 Hz, 1H), 2.16 (d, *J* = 15.1 Hz, 5H), 1.40 (p, *J* = 7.3 Hz, 2H), 1.27 (m, 4H), 0.88 (t, *J* = 6.9 Hz, 3H).^13^C NMR (126 MHz, CDCl_3_) δ 165.91, 148.45, 146.05, 129.88, 127.49, 125.50, 122.73, 122.43, 118.83, 77.37, 77.12, 76.86, 32.21, 31.37, 27.81, 22.45, 20.39, 13.98. MS (ESI-APCI) *m/z* [M + H]^+^ calcd. for C_15_H_22_NO_2_ 248.35, found 248.30.

Synthesis of compound **40**

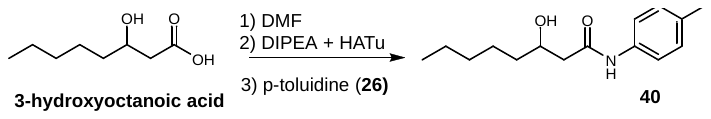

Compound **40** was synthesized in the same method as compound **4.** Compound **40**, yellow solid. ^1^H NMR (500 MHz, acetonitrile-*d_3_*) δ 7.42 (d, *J* = 8.4 Hz, 2H), 7.12 (d, *J* = 8.2 Hz, 2H), 3.95 (m, 1H), 2.44 (dd, *J* = 14.8, 3.6 Hz, 1H), 2.33 (dd, *J* = 14.8, 8.6 Hz, 1H), 2.28 (s, 3H), 1.44 (m, 3H), 1.39 – 1.22 (m, 5H), 0.89 (t, *J* = 6.9 Hz, 3H). ^13^C NMR (126 MHz, acetonitrile-*d_3_*) δ 171.16, 136.84, 133.85, 129.77, 129.77, 120.21, 120.21, 68.88, 44.45, 37.43, 32.12, 25.54, 22.93, 20.42, 13.91. MS (ESI-APCI) *m/z* [M + H]^+^ calcd. for C_15_H_24_NO_2_ 250.18, found 250.30.

**2. Supplementary Tables**

**Table 1.** Protein-protein interaction (PPI) analysis of ColC2 and AsuC5.

| Residues in ColC2 | Residues in AsuC5 | Categrory | Type |
| --- | --- | --- | --- |
| Arg231 | Glu58 | Hydrogen Bond; Electrostatic | Salt Bridge; Attractive Charge |
| Arg272 | Asp40 | Hydrogen Bond; Electrostatic | Salt Bridge; Attractive Charge |
| Glu235 | Arg53 | Electrostatic | Attractive Charge |
| Asn223 | Ser46 | Hydrogen Bond | Conventional Hydrogen Bond; Carbon Hydrogen Bond |
| Ala234 | Arg53 | Hydrogen Bond | Carbon Hydrogen Bond |
| Ala234 | Val49 | Hydrophobic | Alkyl |
| Phe236 | Val19 | Hydrophobic | Pi-Alkyl |
| Phe236 | Leu50 | Hydrophobic | Pi-Alkyl |
| Val100 | Phe64 | Hydrophobic | Pi-Alkyl |

**Table 2.** *X*-ray data collection and structure refinement statistics for ColC2.

| Name | *apo*-olC2 (8K51) | ColC2_**10** (8K56) |
| --- | --- | --- |
| Data collection |  |  |
| Space group | P2_1_2_1_2_1_ | P2_1_2_1_2_1_ |
| Cell dimensions |  |  |
| a, b, c (Å) | 72.41, 88.85, 127.32 | 72.45, 89.29, 127.24 |
| α, β, γ (°) | 90.00, 90.00, 90.00 | 90.00, 90.00, 90.00 |
| Wavelength (Å) | 0.9792 | 0.9792 |
| Resolution (Å) | 56.13-2.15  (2.21-2.15)* | 62.96-2.42  (2.55-2.42) |
| CC1/2 | 0.998 (0.762) | 0.996 (0.681) |
| *R*_merge_ | 0.246 (1.563) | 0.197 (1.317) |
| Average I / σ (I) | 13.3 (2.2) | 10.5 (2.3) |
| Completeness (%) | 100.0 (100.0) | 100.0 (100.0) |
| Multiplicity | 13.4 (13.7) | 8.3 (8.0) |
| Refinement |  |  |
| Resolution (Å) | 51.75-2.15  (2.22-2.15) | 47.81-2.42  (2.51-2.42) |
| No. unique reflections | 45393 | 32143 |
| *R*_work_ / *R*_free_ (%) | 19.45/23.15 | 18.37/22.19 |
| No. atoms |  |  |
| Protein | 4418 | 4434 |
| Ligand / ion | 30 | 114 |
| Water | 249 | 175 |
| B-factors (Å^2^) |  |  |
| Protein | 36.21 | 39.75 |
| Ligand / ion | 64.84 | 68.68 |
| Water | 37.72 | 38.40 |
| R.m.s deviations |  |  |
| Bond lengths (Å) | 0.008 | 0.008 |
| Bond angles (°) | 0.873 | 0.974 |

*Values in parentheses are for the highest-resolution shell.

**Table 3.** Bacterial strains and plasmids used in this study

| Strains | Description | Source |
| --- | --- | --- |
| E. coli DH5α | General cloning host | Invitrogen |
| *E. coli* BL21(DE3) | Host for protein expression | Invitrogen |
| S. lividans TK24 | Host for protein expression | Laboratory storage |
| Plasmids | Description | Source |
| pET28a | protein expression vector | Invitrogen |
| pET21a | protein expression vector | Invitrogen |
| pWY45 | protein expression vector | Laboratory storage |
| pJTU117 | pET28a derived plasmid for Dar12 | This study |
| pJTU118 | pET28a derived plasmid for ColC2 | This study |
| pJTU119 | pET28a derived plasmid for Dar15 | This study |
| pJTU120 | pET28a derived plasmid for Dar6_N | This study |
| pJTU121 | Pwy45 derived plasmid for Dar24 | This study |
| pJTU37 | pET28a derived plasmid for AsuC3 | Laboratory storage |
| pJTU40 | pET21a derived plasmid for AsuC5 | Laboratory storage |
| pJTU42 | pET28a derived plasmid for AsuC7 | Laboratory storage |
| pJTU44 | pET28a derived plasmid for AsuC89 | Laboratory storage |
| pJTU45 | pET28a derived plasmid for ScMatB | Laboratory storage |
| pJTU46 | pET28a derived plasmid for SaFabD | Laboratory storage |
| pJTU47 | pET28a derived plasmid for Sfp | Laboratory storage |
| pJTU122 | pET28a derived plasmid for ColC2-Y41A | This study |
| pJTU123 | pET28a derived plasmid for ColC2-M43A | This study |
| pJTU124 | pET28a derived plasmid for ColC2-C72A | This study |
| pJTU125 | pET28a derived plasmid for ColC2-N71Y | This study |
| pJTU126 | pET28a derived plasmid for ColC2-L73Y | This study |
| pJTU127 | pET28a derived plasmid for ColC2-K99A | This study |
| pJTU128 | pET28a derived plasmid for ColC2-V100A | This study |
| pJTU129 | pET28a derived plasmid for ColC2-D109A | This study |
| pJTU130 | pET28a derived plasmid for ColC2-H110A | This study |
| pJTU131 | pET28a derived plasmid for ColC2-D125A | This study |
| pJTU132 | pET28a derived plasmid for ColC2-F128A | This study |
| pJTU133 | pET28a derived plasmid for ColC2-R231A | This study |
| pJTU134 | pET28a derived plasmid for ColC2-E234A | This study |
| pJTU135 | pET28a derived plasmid for ColC2-F236A | This study |
| pJTU136 | pET28a derived plasmid for ColC2-R269A | This study |
| pJTU137 | pET28a derived plasmid for ColC2-R272A | This study |
| pJTU138 | pET21a derived plasmid for AsuC5-D40A | This study |
| pJTU139 | pET21a derived plasmid for AsuC5-S41A | This study |
| pJTU140 | pET21a derived plasmid for AsuC5-D53A | This study |
| pJTU141 | pET21a derived plasmid for AsuC5-E58A | This study |

**Table 4.** Primers used in this study.

| Primer | Oligonucleotide sequence (5’ to 3’) |
| --- | --- |
| Dar15-pET28a-F | tttcagggatccCAtATGAGCTCTCGTGACGACATC |
| Dar15-pET28a-R | TCGACGGAGCTCGAATTCTTACACCGCCGCGGAACG |
| Dar6-pET28a-F | ttttcagggatccCAtATGCGTACCGCAGAAACCGATC |
| Dar6-pET28a-R | GTCGACGGAGCTCGAATTCTCAGGTAGGAGGTTCACCG |
| Dar24-pWY45-F | TGGTGCCGCGCGGCAGCCATATGTGCGGTATAACCGGA |
| Dar24-pWY45-R | GGTACCGAGCTCGAATTCtcaGGCCAGGGTGATGTCCGG |
| For site directed mutagenesis | |
| ColC2-Y41A-F | GTGCCGGCCGAAATGCTGGATAGTTTT |
| ColC2-Y41A-R | CATTTCGGCCGGCACACTACGCAGATG |
| ColC2-M43A-F | GTATGAGCAGCTGGATAGTTTTGATGGC |
| ColC2-M43A-R | ATCCAGTGCTTCATACGGCACACTACG |
| ColC2-N71Y-F | GGCGGCTACTGTCTGGAAAGCACCCCG |
| ColC2-N71Y-R | CAGACAGTAGCCGCCGCGGCGACGATG |
| ColC2-C72A-F | GGCAATGCCCTGGAAAGCACCCCGCT |
| ColC2-C72A-R | TTCCAGGGCATTGCCGCCGCGGCGAC |
| ColC2-L73Y-F | AATTGTTACGAAAGCACCCCGCTGTTT |
| ColC2-L73Y-R | GCTTTCGTAACAATTGCCGCCGCGGCG |
| ColC2-K99A-F | ATTTGGGCAGTTAGCGGTGAATGGTGG |
| ColC2-K99A-R | GCTAACTGCCCAAATCTGTGCCGGCACC |
| ColC2-V100A-F | TGGAAAGCAAGCGGTGAATGGTGGGAT |
| ColC2-V100A-R | ACCGCTTGCTTTCCAAATCTGTGCCGG |
| ColC2-D109A-F | GCATGGGCCCATCTGCTGCTGATTGT |
| ColC2-D109A-R | CAGATGGGCCCATGCATCCCACCATTC |
| ColC2-H110A-F | ATGGGATGCCCTGCTGCTGATTGTTAC |
| ColC2-H110A-R | CAGCAGGGCATCCCATGCATCCCACC |
| ColC2-D125A-F | CTGCTGGCCGTTGGCTTTCTGATGCT |
| ColC2-D125A-R | GCCAACGGCCAGCAGCCAATCTTCAC |
| ColC2-F128A-F | GTTGGCGCACTGATGCTGACCTTTGCA |
| ColC2-F128A-R | CATCAGTGCGCCAACATCCAGCAGCCA |
| ColC2-R231A-F | AATGGCGCCGTTAGCGCAGAATTTATT |
| ColC2-R231A-R | GCTAACGGCGCCATTACGTGCATGCAG |
| ColC2-E235A-F | AGCGCAGCATTTATTGAAACCACCAGT |
| ColC2-E235A-R | AATAAATGCTGCGCTAACACGGCCATT |
| ColC2-F236A-F | GCAGAAGCAATTGAAACCACCAGTCGC |
| ColC2-F236A-R | TTCAATTGCTTCTGCGCTAACACGGCC |
| ColC2-R269A-F | AAAGCAGCAGCAGATCGCAGTACCCGT |
| ColC2-R269A-R | ATCTGCTGCTGCTTTTTCCCAGGTGCG |
| ColC2-R272A-F | GCAGATGCAAGTACCCGTCGCAGCCTG |
| ColC2-R272A-R | GGTACTTGCATCTGCACGTGCTTTTTC |
| AsuC5-D40A-F | GAACTCGCATCGCTGTCGATCGTGTCC |
| AsuC5-D40A-R | CAGCGATGCGAGTTCCAGGTCGTCGAC |
| AsuC5-S41A-F | CTCGACGCACTGTCGATCGTGTCCGTG |
| AsuC5-S41A-R | CGACAGTGCGTCGAGTTCCAGGTCGTC |
| AsuC5-D53A-F | GTCCAGGCACGCTTCGGCGTCGAGGTG |
| AsuC5-D53A-R | GAAGCGTGCCTGGACCAGGACGAACAC |
| AsuC5-E58A-F | GGCGTCGCCGTGCCGAACGAGATCTTC |
| AsuC5-E58A-R | CGGCACGGCGACGCCGAAGCGGCGCTG |

**Table 5.** Accession number of the protein sequences used for the phylogenetic analysis and sequence alignment.

| Type | Name | Access. No. | a.a. | Strain |
| --- | --- | --- | --- | --- |
| Type I  (Function in xenobiotic detoxification by transferring acetyl-CoA to arylamine) | 1W6F | 1W6F | 278 | *Mycolicibacterium smegmatis* |
|  | 1GX3 | 1GX3 | 284 | *Mycolicibacterium smegmatis* |
|  | 2VFC | 2VFC | 280 | *Mycobacterium marinum* |
|  | 3D9W | 3D9W | 293 | *Nocardia farcinica* |
|  | 4GUZ | 4GUZ | 284 | *Mycobacteroides abscessus* ATCC 19977 |
|  | CetD | CetD | 288 | *Actinomyces* sp. Lu 9419 |
|  | BezG | BezG | 282 | *Streptomyces* sp. RI18 |
|  | 4GUZ | 4GUZ | 284 | [*Mycobacterium abscessus*](https://www.ncbi.nlm.nih.gov/Structure/mmdb/mmdbsrv.cgi?uid=114463) |
| Type I  (Transfer acyl-CoA to arylamine) | PtmC | AIW55569.2 | 291 | *Streptomyces platensis* |
|  | PtnC | ADD82996.1 | 291 | *Streptomyces platensis* |
|  | PtmC1 | ACO31290.1 | 291 | *Streptomyces platensis* MA7327 |
|  | A0A0L0JZ49.1 | A0A0L0JZ49.1 | 263 | [*Streptomyces acidiscabies*](https://www.uniprot.org/taxonomy/42234) |
|  | A0A0L8LL64 | A0A0L8LL64 | 265 | [*Streptomyces decoyicus*](https://www.uniprot.org/taxonomy/249567) |
|  | A0A101UF70.1 | A0A101UF70.1 | 268 | [*Streptomyces* sp. DSM 15324](https://www.uniprot.org/taxonomy/1739111) |
|  | NybK | UZQ45017.1 | 266 | *Streptomyces albus subsp. chlorinus* |
|  | DaqT | RZB16697.1 | 270 | *Streptomyces* sp. F001 |
|  | DadS | RZB16698.1 | 270 | *Streptomyces* sp. F001 |
|  | MicB006_2933 | MicB006_2933 | 269 | *Micromonospora* |
|  | MicB006_2934 | MicB006_2934 | 279 | *Micromonospora* |
|  | A0A9W6HWV2 | A0A9W6HWV2 | 258 | *Micromonospora* |
|  | D2AUQ7 | D2AUQ7 | 254 | *Streptosporangium roseum* |
|  | A0A344LBQ6 | A0A344LBQ6 | 249 | *Amycolatopsis albispora* |
| Type II  (catalyze macrolactamization of ACP-tethered polyketide) | RifF | AAC01715.1 | 260 | *Amycolatopsis mediterranei* S699 |
|  | RmpF | AWH12663.1 | 260 | *Amycolatopsis* sp. |
|  | KngF | WP_004559807.1 | 258 | *Amycolatopsis vancoresmycina* |
|  | CxmF | CQR60492.1 | 269 | *Streptomyces leeuwenhoekii* |
|  | Sare_1251 | ABV97156.1 | 258 | *Salinispora arenicola* CNS-205 |
|  | NatF | ADM46361.1 | 289 | *Streptomyces* sp. CS |
|  | StvF | ASZ00152.1 | 264 | *Streptomyces* *spectabilis* |
|  | RubF | CAI94702.1 | 264 | *Streptomyces rubradiris* |
|  | NamF | AID50087.1 | 256 | *Streptomyces* sp. LZ35 |
|  | DivN | CCP20052.1 | 251 | *Streptomyces* sp. HKI0576 |
|  | HgcF | AFV30252.1 | 294 | *Streptomyces* sp. LZ35 |
|  | Asm9 | AAM54087.1 | 259 | *Actinosynnema pretiosum subsp. auranticum* |
|  | Ansa_11 | AQZ37096.1 | 259 | *Actinosynnema pretiosum subsp. pretiosum* |
|  | Asc9 | XOI27400.1 | 262 | *Amycolatopsis alba* DSM 44262 |
|  | Mas10 | ATY46593.1 | 260 | *Micromonospora* sp. HK160111 |
|  | MycE | AFG19424.1 | 260 | *Streptomyces flaveolus* |
|  | MbcF | ACF35448.1 | 258 | *Actinosynnema pretiosum subsp. pretiosum* |
|  | GdmF | ABI93780.1 | 257 | *Streptomyces hygroscopicus* |
|  | GelD | ABB86411.1 | 257 | *Streptomyces hygroscopicus subsp. duamyceticus* |
| Type III | VerC2 | QTW52588.1 | 297 | *Streptomyces verdensis* |
| (CoCl2-like enzymes) | A0A345SU73 | A0A345SU73 | 285 | *Peterkaempfera bronchialis* |
|  | ManC2 | QTW52592.1 | 288 | *Streptomyces parvulus* |
|  | A0A542UF51 | A0A542UF51 | 303 | *Streptomyces puniciscabiei* |
|  | ColC2 | AIL50169.1 | 285 | *Streptomyces aureus* SOK1/5-04 |
|  | A0A7Y6LT24 | A0A7Y6LT24 | 290 | *Streptomyces* sp. CAI 127 |
|  | A0A2P8BGJ1 | A0A2P8BGJ1 | 304 | *Streptomyces* sp. CS149 |
|  | Dar12 | MCI0386312.1 | 291 | *Streptomyces sp. CNQ-085* |
|  | EspC2 | WP_051075499.1 | 305 | *Saccharothrix espanaensis* DSM44229 |
|  | A0A841CUK3 | A0A841CUK3 | 290 | *Saccharothrix tamanrassetensis* |
|  | WP_198045673.1 | WP_198045673.1 | 282 | *Kitasatospora mediocidica* |
|  | AsuC2 | D7P5V8 | 269 | *Streptomyces nodosus subsp. asukaensis* |
|  | Pac17 | WP_018222380.1 | 293 | *Salinisporapacifica* CNT-855 |
|  | A0A1C4NEK0 | A0A1C4NEK0 | 269 | unclassified *Streptomyces* |
| Unknown NAT | A0A7C6TJ72 | A0A7C6TJ72 | 304 | *Brevibacterium sp.* |
|  | A0A0G3GST4 | A0A0G3GST4 | 277 | *Corynebacterium epidermidicanis* |
|  | A0A1H1Y0D6 | A0A1H1Y0D6 | 304 | *Brevibacterium siliguriense* |
|  | A0A2H1JDL1 | A0A2H1JDL1 | 304 | *Brevibacterium linens* |
|  | A0A2H1KWW3 | A0A2H1KWW3 | 304 | *Brevibacterium aurantiacum* |
|  | A0A0B9APS6 | A0A0B9APS6 | 304 | *Brevibacterium linens* |
|  | A0A2A3X5H5 | A0A2A3X5H5 | 335 | *Brevibacterium aurantiacum* |
|  | A0A1D7W005 | A0A1D7W005 | 304 | *Brevibacterium* |
|  | A0A2H1KJW7 | A0A2H1KJW7 | 304 | *Brevibacterium* |
|  | A0A556C8S4 | A0A556C8S4 | 304 | *Brevibacterium aurantiacum* |
|  | K0K6F8 | K0K6F8 | 292 | *Pseudonocardiaceae* |
|  | D2B8T2 | D2B8T2 | 312 | *Streptosporangium roseum* |
|  | C7QCQ8 | C7QCQ8 | 288 | *Catenulispora acidiphila* |
|  | C6WGE4 | C6WGE4 | 284 | *Actinosynnema mirum* |
|  | D3PUD5 | D3PUD5 | 283 | *Stackebrandtia nassauensis* |
|  | D3PYB7 | D3PYB7 | 274 | *Stackebrandtia nassauensis* |
|  | R4TFE4 | R4TFE4 | 300 | *Amycolatopsis* |
|  | M2ZB65 | M2ZB65 | 290 | *Amycolatopsis decaplanina* |
|  | W7JBN4 | W7JBN4 | 299 | *Actinokineospora spheciospongiae* |
|  | F2R1X7 | F2R1X7 | 288 | *Streptomyces* |
|  | E9V1N1 | E9V1N1 | 281 | *Nocardioidaceae bacterium* Broad-1 |
|  | A0A944JWF4 | A0A944JWF4 | 271 | *Streptomyces* sp*.* ISL-94 |
|  | A0A1Q5MVX9 | A0A1Q5MVX9 | 281 | *Streptomyces* sp. CB00455 |
|  | A0A2Z4V2I7 | A0A2Z4V2I7 | 341 | unclassified *Streptomyces* |
|  | A0A0M8YQ47 | A0A0M8YQ47 | 272 | unclassified *Streptomyces* |

**3. Supplementary Figures**

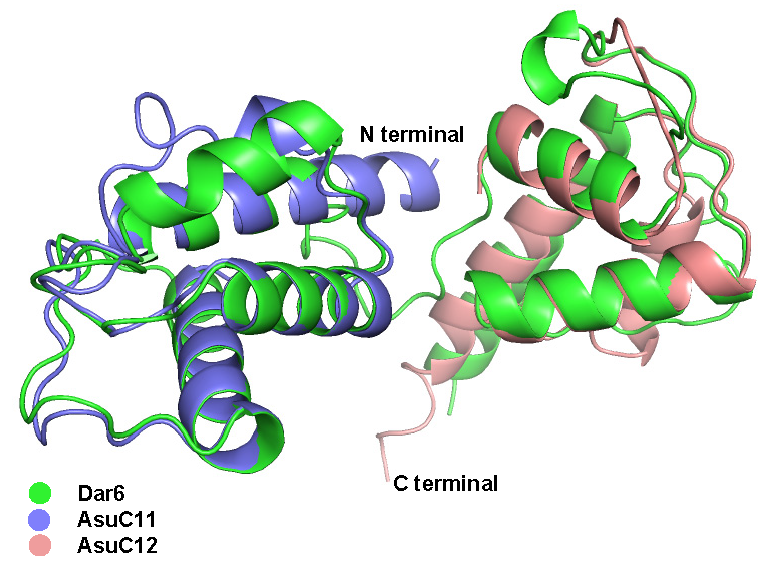

**Fig. 1 |** **Structural alignment and comparative modeling of ACPs.** Structural superposition of the ACP homologs Dar6 (green), AsuC11 (blue), and AsuC12 (magenta). Dar6 is shown in its unique didomain architecture, while AsuC11 and AsuC12 represent standalone ACPs. All protein models were generated using AlphaFold2. The alignment highlights the conserved helical bundle motif characteristic of the phosphopantetheine-binding fold.

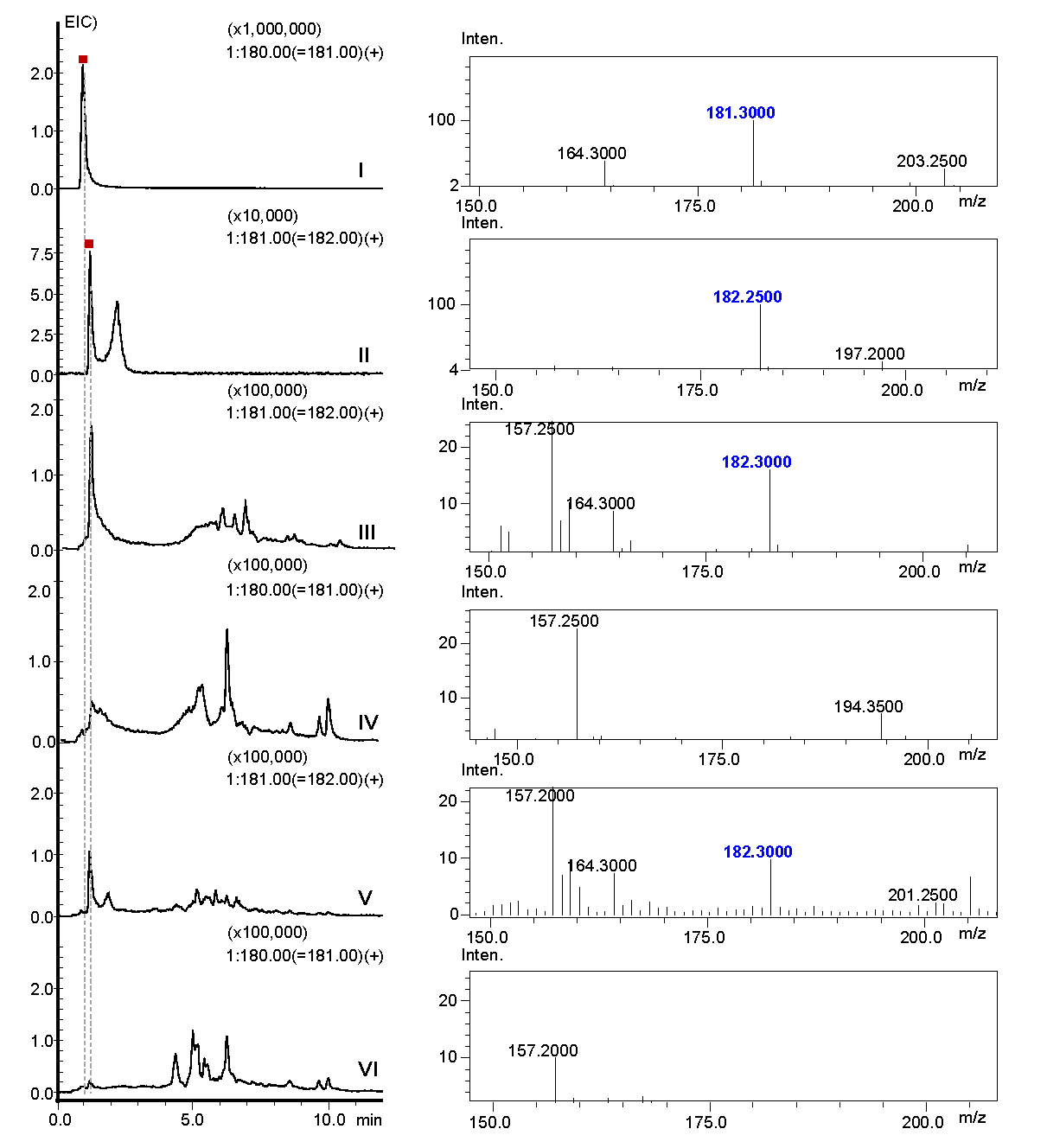

**Fig. 2 |** **UPLC-MS analysis of feeding assays with compound 1.** Extracted ion chromatograms (EICs) at *m*/*z*=181 and 182 showing the metabolic profile of *S. lividans* TK24 following the administration of **1**. (I) Synthetic standard of **3**, *m*/*z* 181. (II) Synthetic standard of **1**, *m*/*z* 182. (III–IV) *S. lividans* TK24 expressing Dar24 fed with **1**; (III) and (IV) indicate the presence of the substrate **1** (EIC at *m*/*z* =182) but the absence of product **3** (EIC at *m*/*z* =181) respectively. (V–VI) Negative control using wild-type *S. lividans* TK24 fed with **1**; (V) and (VI) indicate the presence of the substrate **1** (EIC at *m*/*z* =182) but the absence of product **3** (EIC at *m*/*z* =181) respectively confirm no endogenous conversion to **3**.

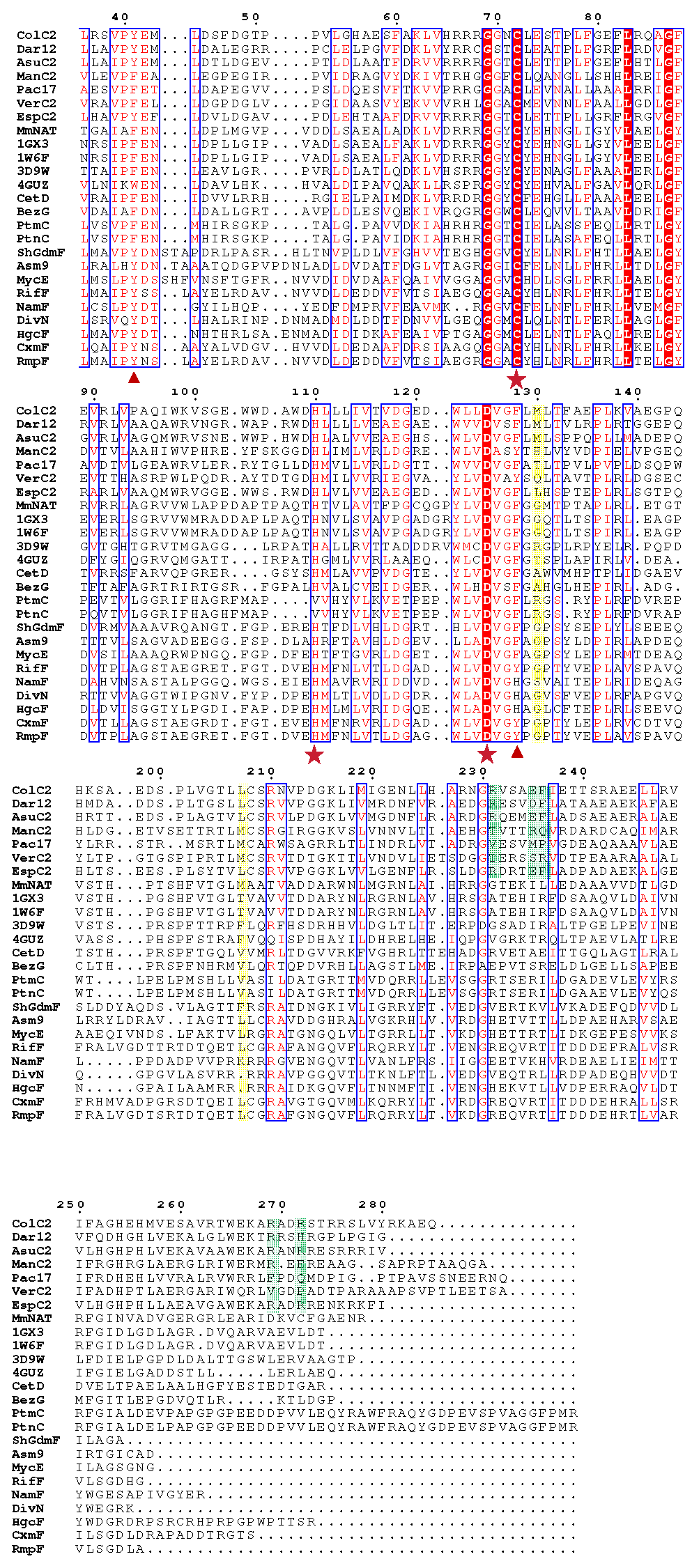

**Fig. 3 | Sequence alignment of ColC2 with representative arylamine NATs.** Multiple sequence alignment highlighting conserved and functional residues across the arylamine NAT family. The catalytic Cys-His-Asp triad is denoted by red stars (★). Residues participating in arylamine binding are marked with red triangles (▲). Amino acids forming the architecture of the unique substrate-binding pocket in ColC2 are highlighted in yellow, while those proposed to facilitate interactions with the ACP are shaded in green. Sequence accession numbers and the full list of aligned NATs are provided in Table 5.

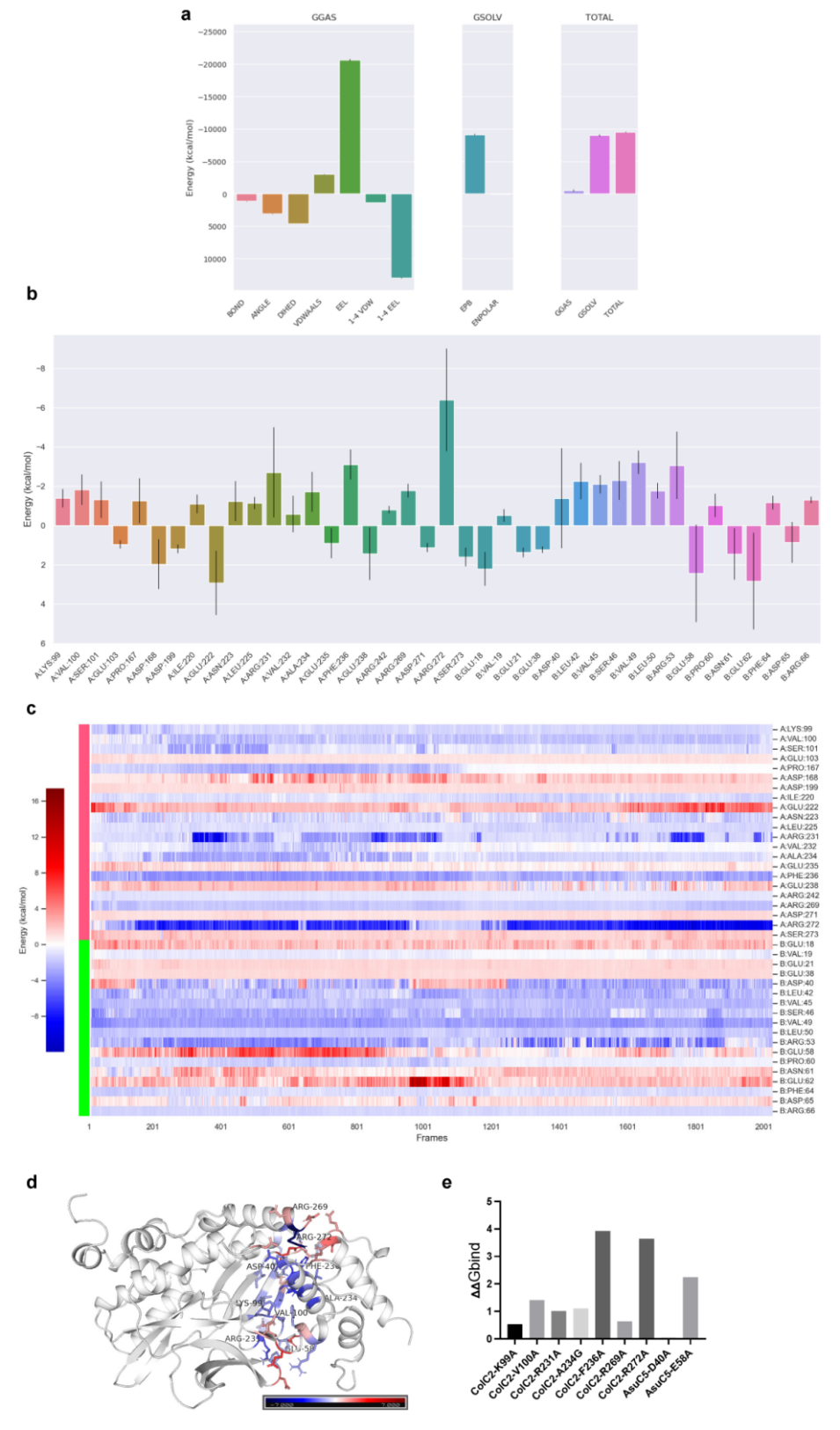

**Fig. 4 | MM/PBSA binding energy analysis of the ColC2–AsuC5 complex. a**, Predicted total binding free energies (*ΔG_bind_*) for the ColC2–AsuC5 protein–protein complex. **b**, Per-residue energy decomposition identifying the major energetic contributors to complex formation. **c**, Evolution of individual residue energy contributions monitored over a 20 ns molecular dynamics (MD) simulation trajectory. **d**, Structural mapping of energy decomposition results onto the complex interface; residues are colored by binding energy, where red indicates unfavorable (positive binding energy) and blue indicates favorable (negative binding energy) contributions. Color intensity is proportional to the absolute energy value. **e**, In silico alanine scanning mutagenesis. *ΔΔG_bind_* represents the change in binding free energy between the wild-type (WT) and mutant complexes, highlighting critical residues for the protein–protein interaction.

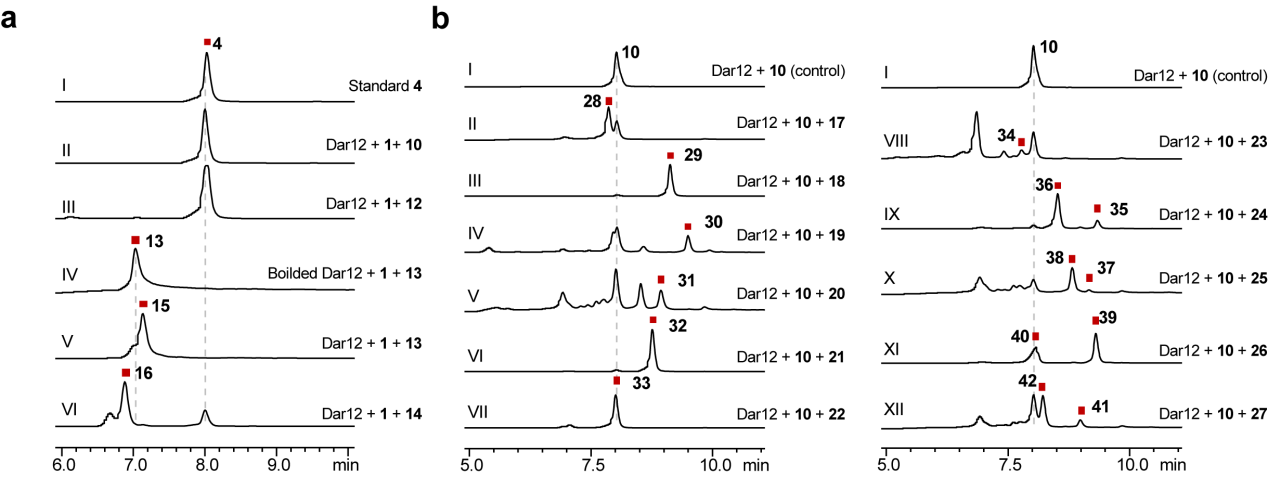

**Fig. 5 | Substrate specificity and biocatalytic scope of Dar12.** **a**, Evaluation of acyl donor promiscuity. HPLC traces show the conversion of acyl acceptor substrate **1** when reacted with Dar12 in the presence of various synthetic acyl-thioesters and acyl-CoA donors. **b**, Evaluation of acyl acceptor promiscuity. HPLC analysis of reactions containing Dar12, the "upper" chain acyl donor **10**, and a library of arylamine acceptors, including 3,4-AHBA (**17**), 2-amino-4-methylphenol (**18**), 2-amino-6-methylphenol (**19**), 2-amino-3-methylphenol (**20**), 2-aminophenol (**21**), 3-aminophenol (**22**), 4-aminophenol (**23**), m-toluidine (**24**), o-toluidine (**25**), p-toluidine (**26**), and aniline (**27**). Trace labels correspond to the specific acceptor used in each assay; peaks were verified by HR-MS (Fig. 6).

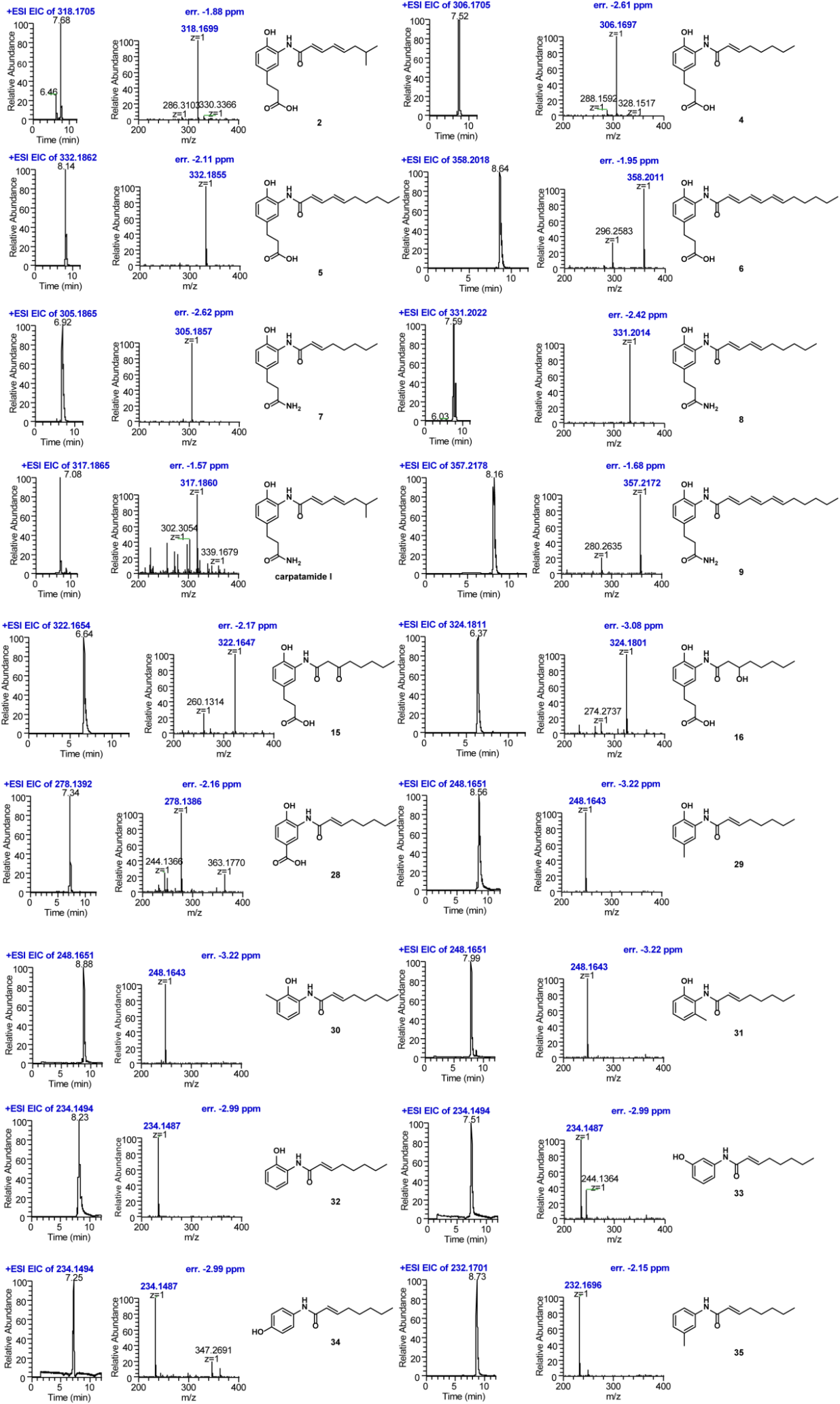

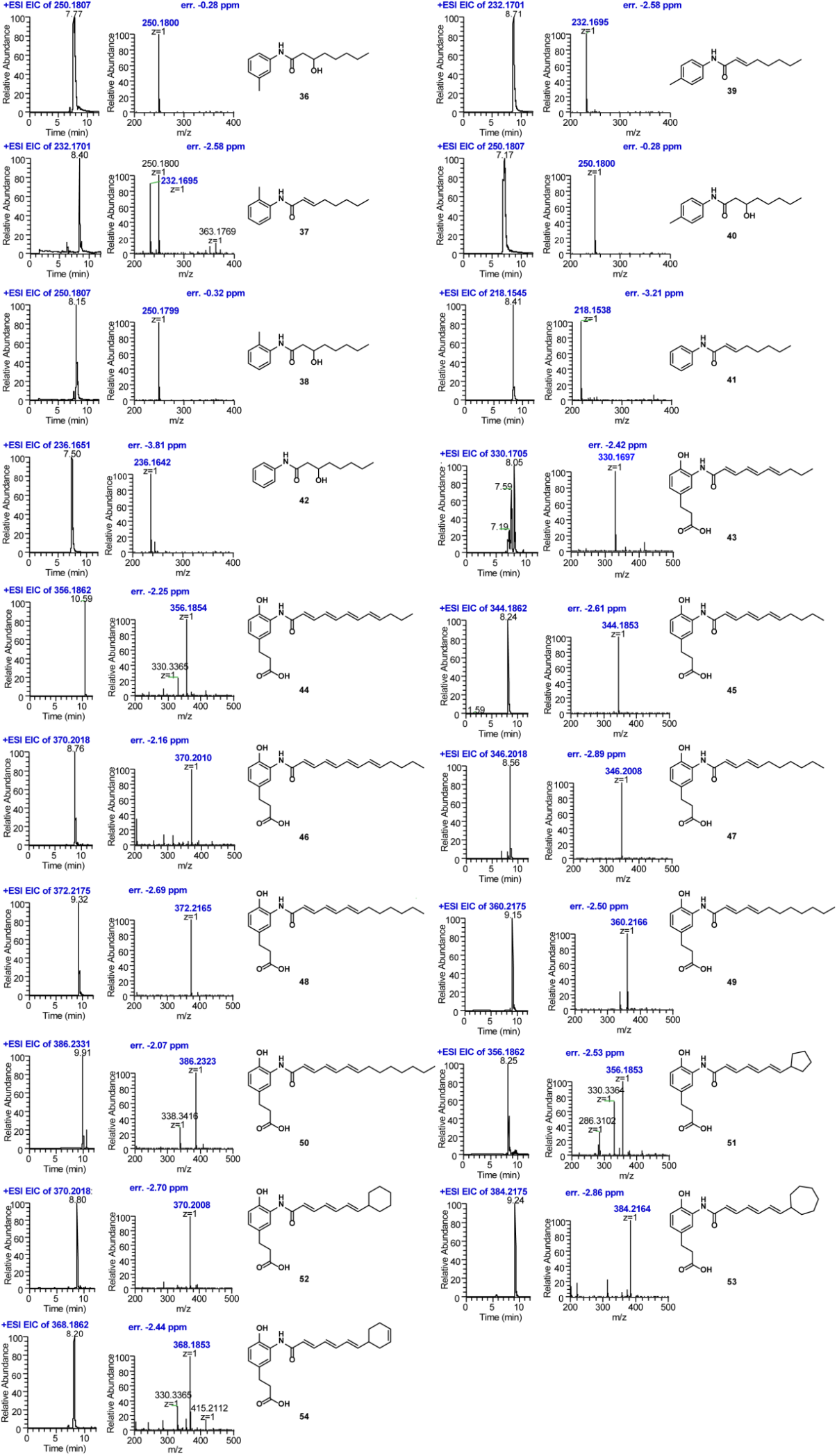

**Fig. 6 | HR-ESI-MS analysis of synthetic and biosynthetic polyketide amides.** High-resolution electrospray ionization mass spectrometry (HR-ESI-MS) spectra for compounds **2**, **4–9**, **15**, **16**, **28–54**, and carpatamide I. Observed *m/z* values for each compound are consistent with the calculated [M + H]^+^ ions (error < 5 ppm), confirming the molecular formulas and successful enzymatic conversion across a diverse range of acyl donors and acceptors. Detailed mass spectra and fragmentation patterns for all compounds are provided in the Supplementary Information.

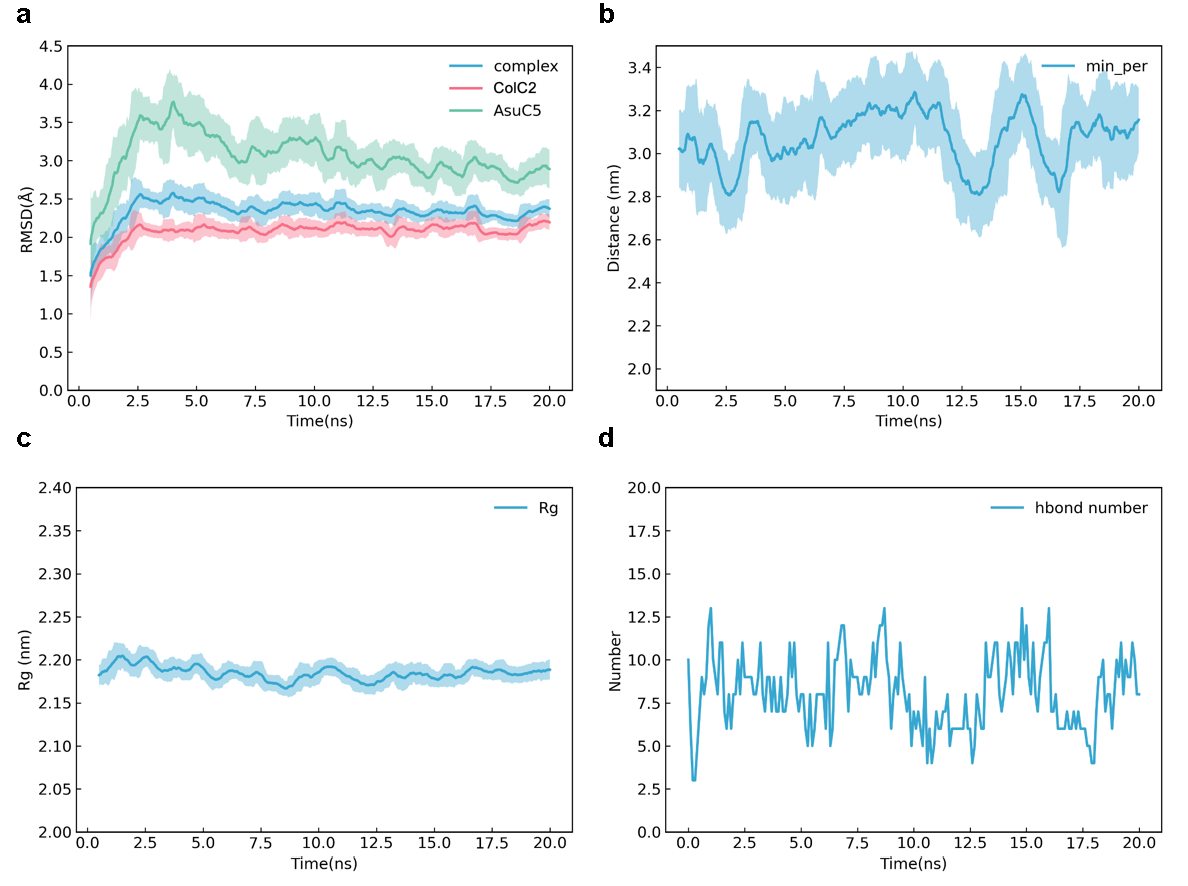

**Fig. 7 | Molecular dynamics simulations of the ColC2–AsuC5 complex.** **a**, Root-mean-square deviation (RMSD) profiles over a 20 ns MD trajectory. The plots show the structural stability of the total complex (green), the ColC2 enzyme (red), and the AsuC5 carrier protein (blue). **b**, Minimum distance between ColC2 and AsuC5 throughout the 20 ns simulation, indicating the maintenance of a stable protein–protein interface. **c**, Radius of gyration (*R_g_*) for the ColC2–AsuC5 complex, reflecting the compactness of the heterodimer during the simulation. **d**, Time-dependent analysis of the number of interfacial hydrogen bonds formed between ColC2 and AsuC5.

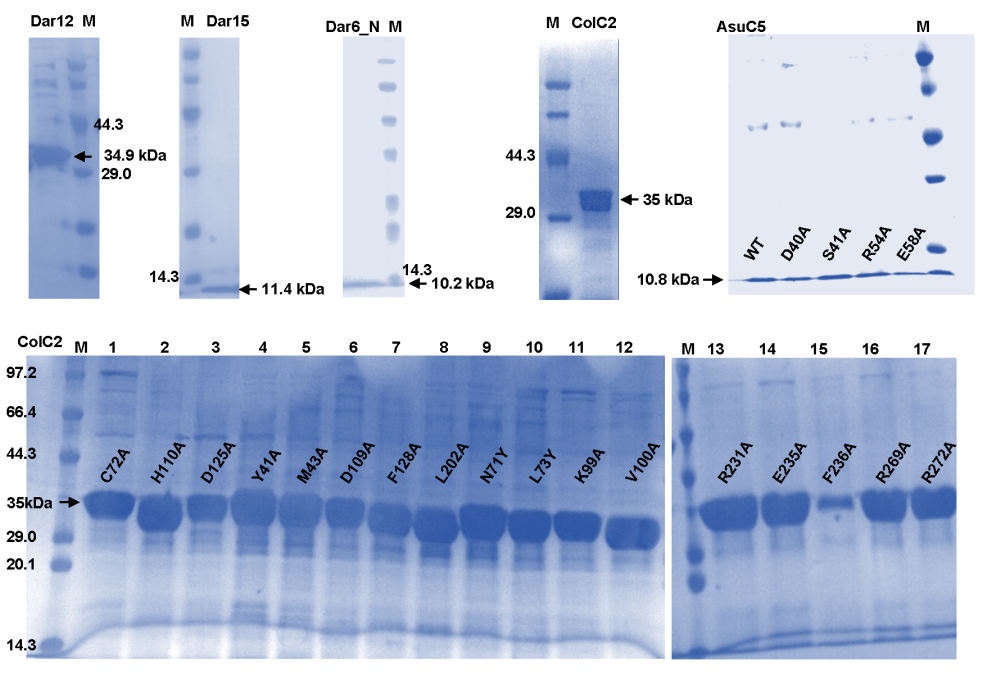

**Fig. 8 | SDS-PAGE analysis of purified recombinant enzymes and mutants.** Denaturing SDS-PAGE (12.5% polyacrylamide) showing the homogeneity and molecular weight of purified proteins used in this study, including **Dar12**, **Dar15**, **Dar6_N**, **ColC2**, and the carrier protein **AsuC5**. Panels also display the representative purification of **AsuC5** and **ColC2** site-directed mutants. Molecular weight markers (kDa) are indicated on the left. All proteins were purified via Ni-NTA affinity chromatography.

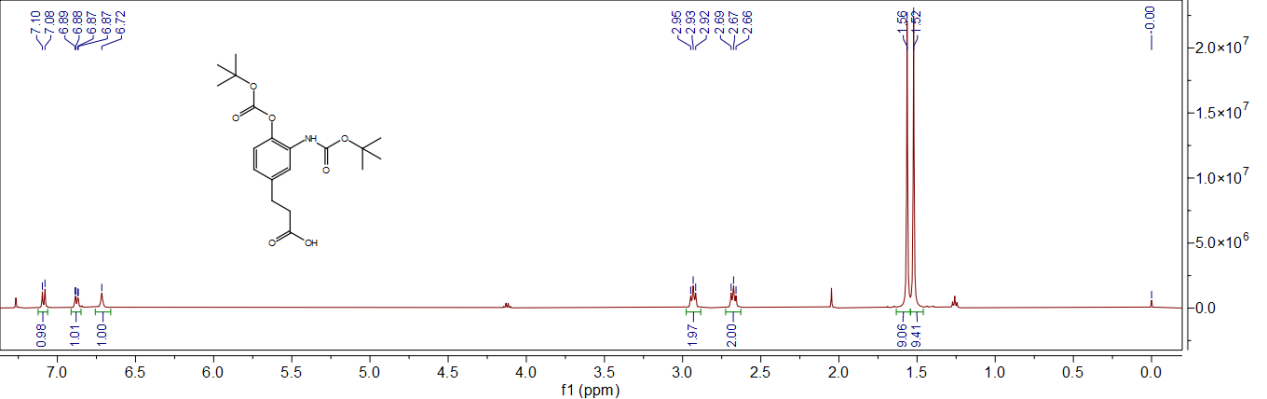
 **Fig. 9** **| ^1^H (500 MHz) NMR spectra of Int-1 in CDCl_3_.**

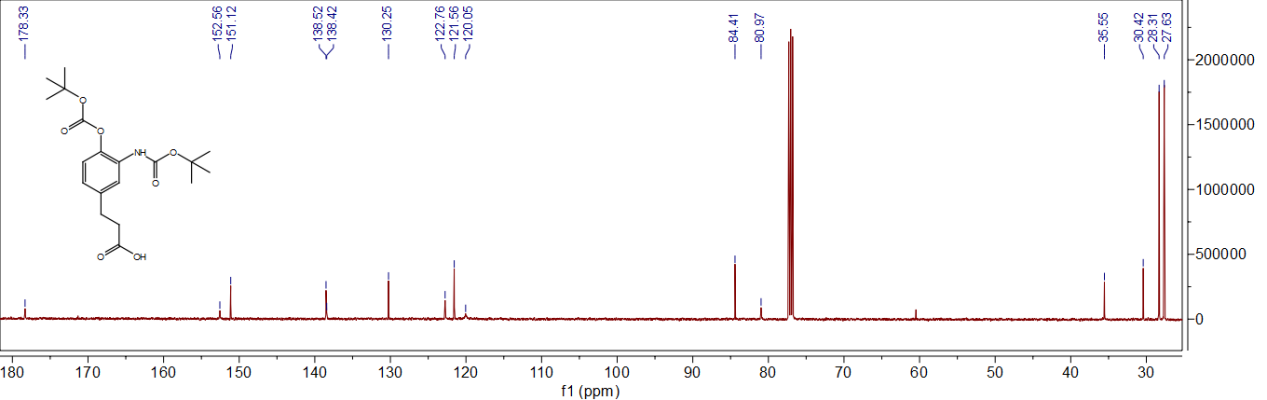

**Fig. 10 | ^13^C (126 MHz) NMR spectra of Int-1 in CDCl_3_.**

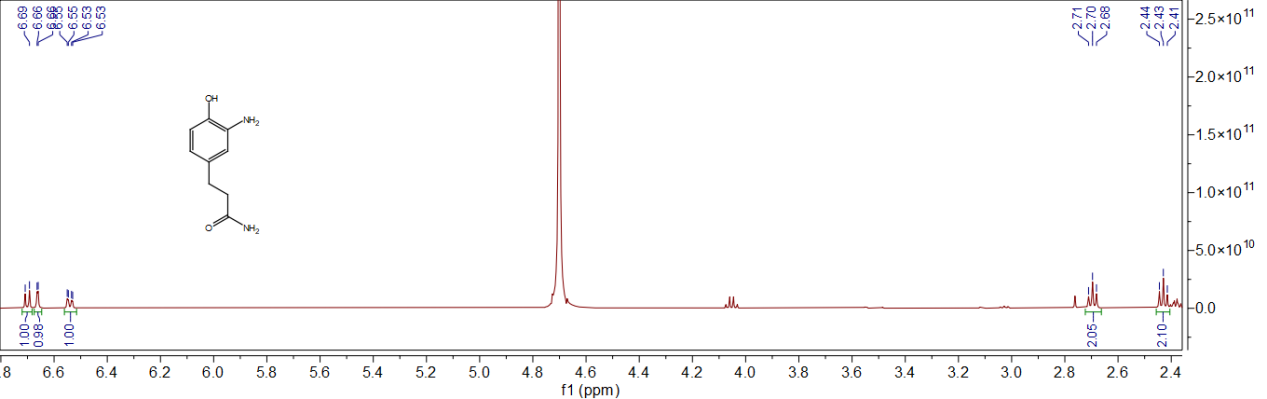

**Fig. 11 | ^1^H (500 MHz) NMR spectra of compound 3 in D_2_O.**

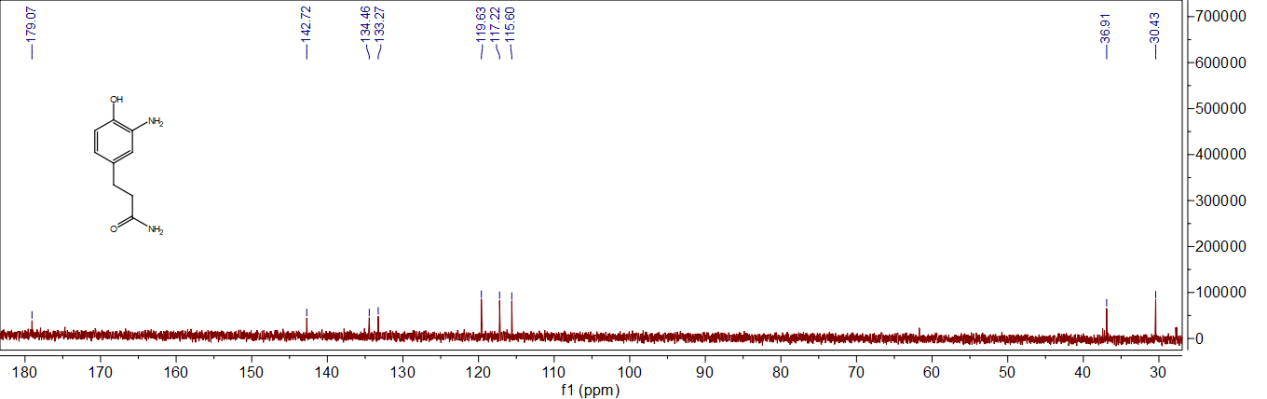

**Fig. S12 | ^13^C (126 MHz) NMR spectra of compound 3 in D_2_O.**

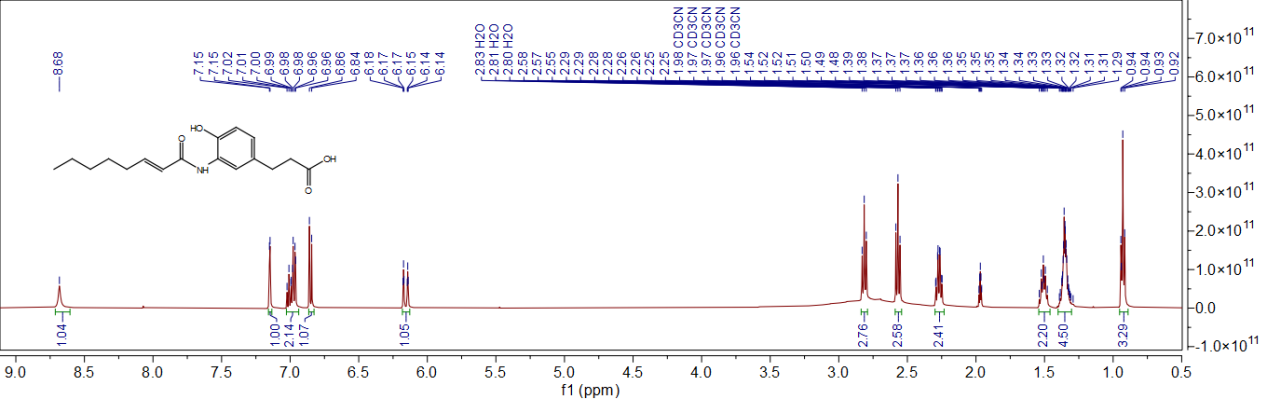

**Fig. 13 | ^1^H (500 MHz) NMR spectra of compound 4 in CD_3_CN.**

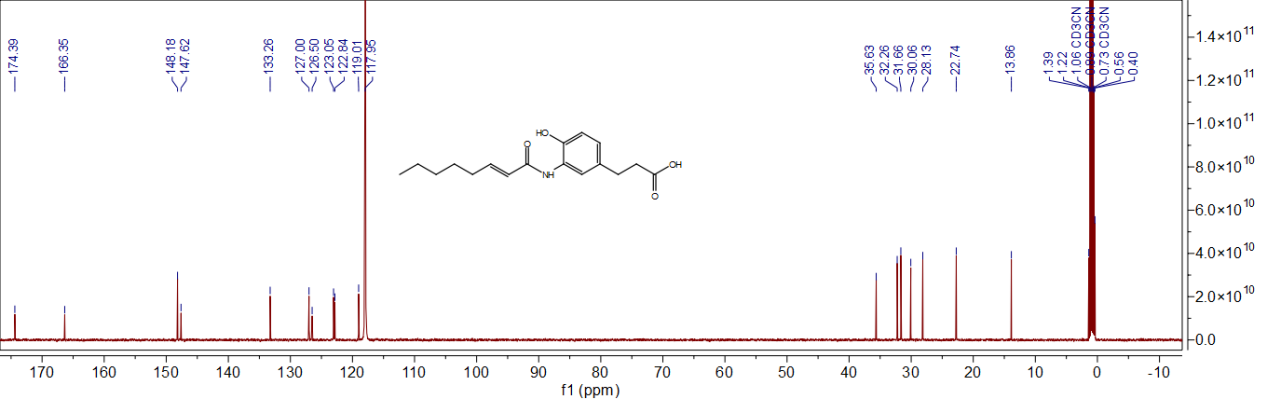
**Fig. 14 | ^13^C (126 MHz) NMR spectra of compound 4 in CD_3_CN.**

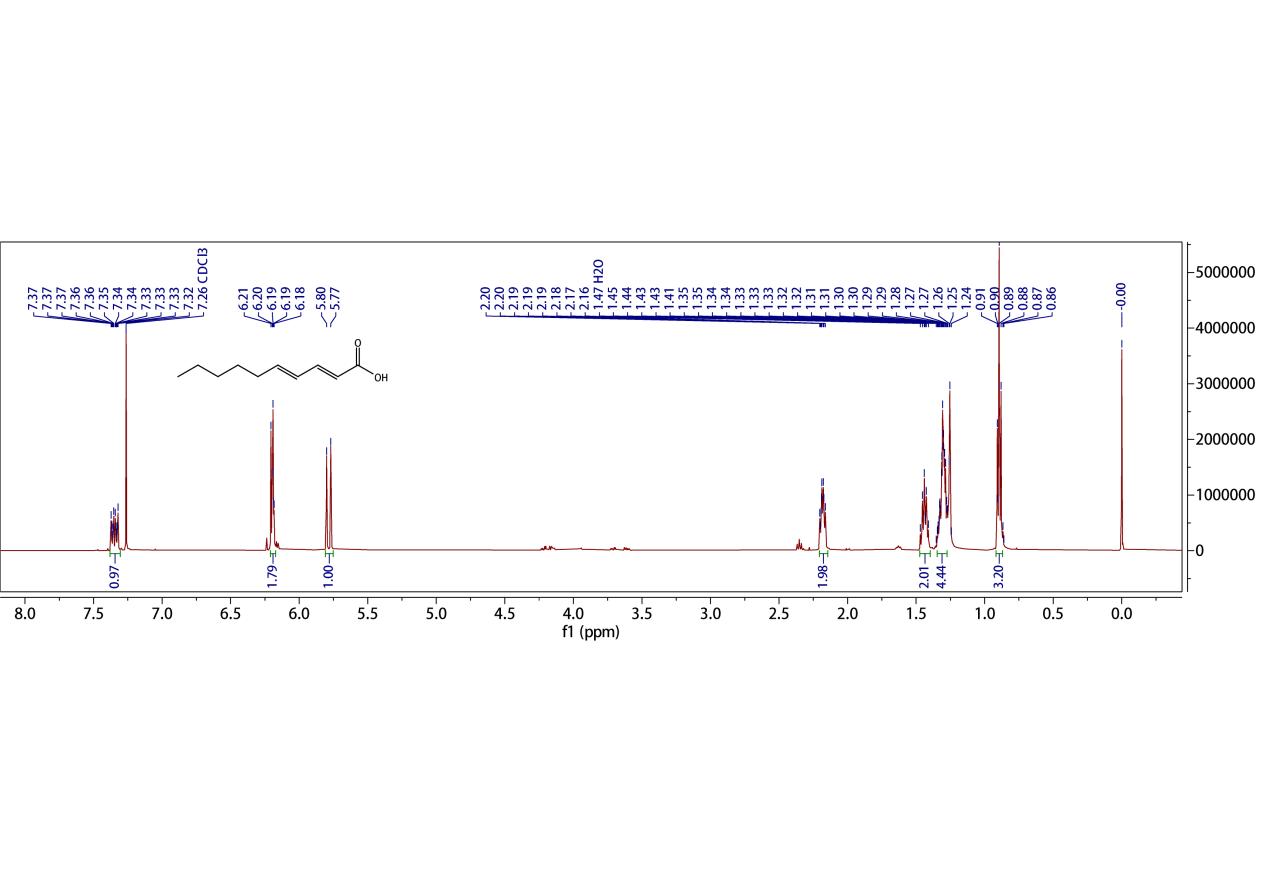

**Fig. 15 | ^1^H NMR (500 MHz) spectra of (2*E*,4*E*)-deca-2,4-dienoic acid in CDCl_3._**

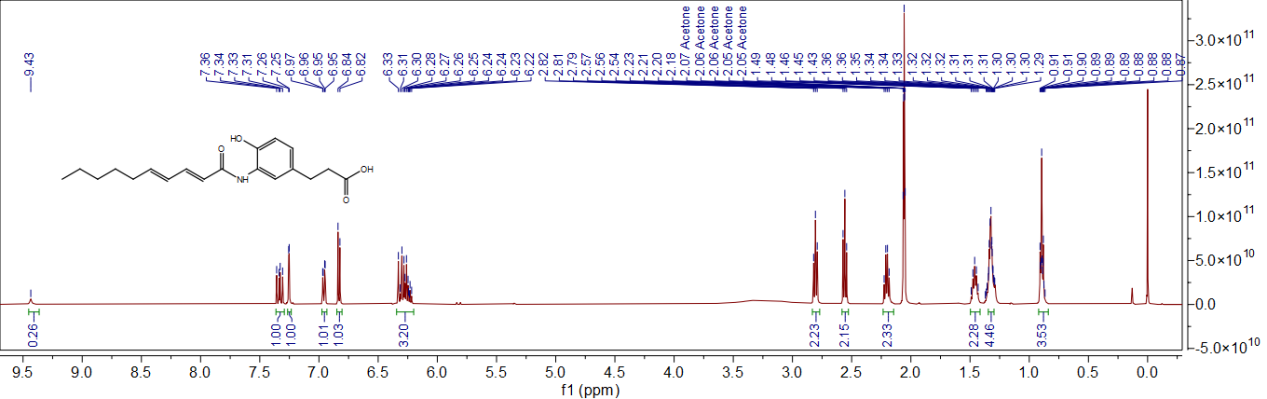

**Fig. 16 | ^1^H (500 MHz) NMR spectra of compound 5 in acetone-*d_6_*.**

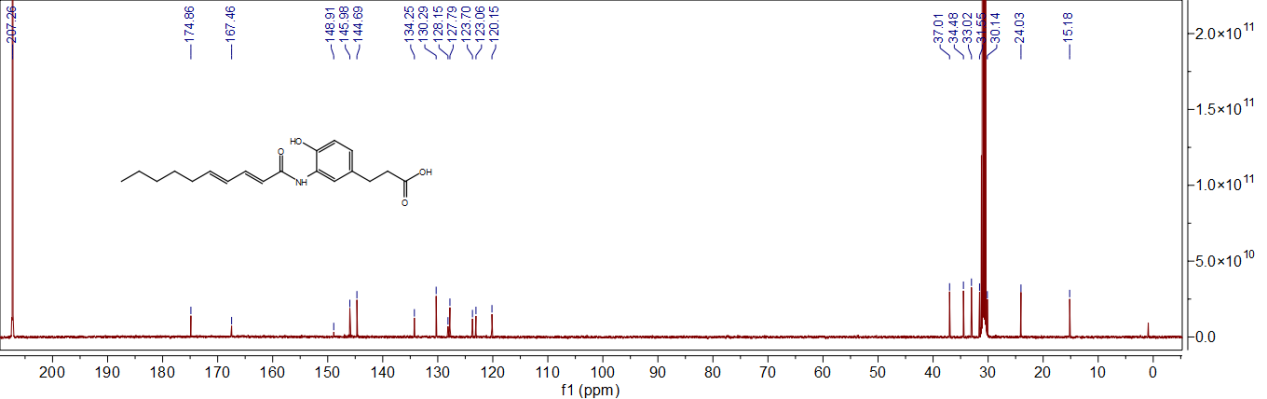

**Fig. 17 | ^13^C (126 MHz) NMR spectra of compound 5 in acetone-*d_6_*.**

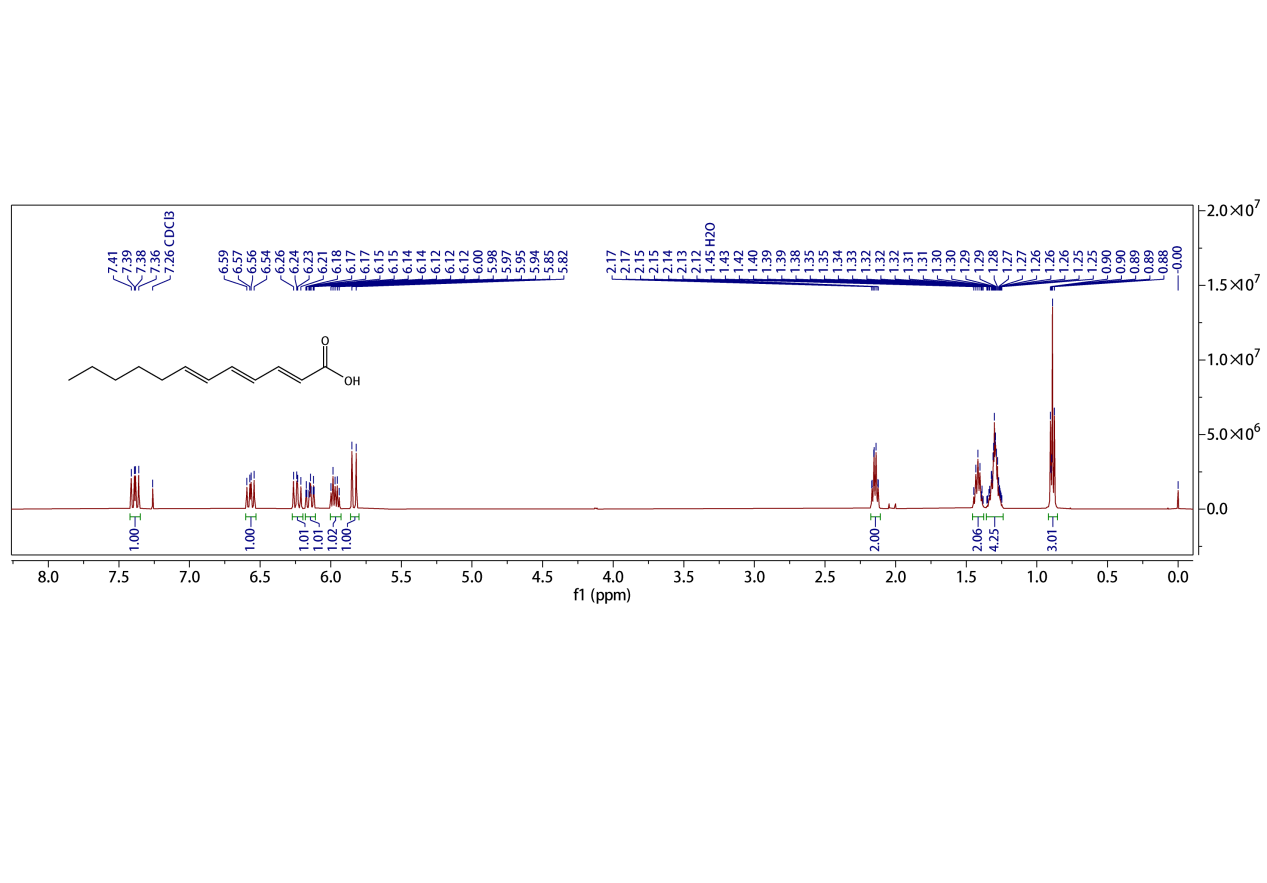

**Fig. 18 | ^1^H NMR (500 MHz) spectra of (2*E*,4*E*,6*E*)-dodeca-2,4,6-trienoic acid in CDCl_3._**

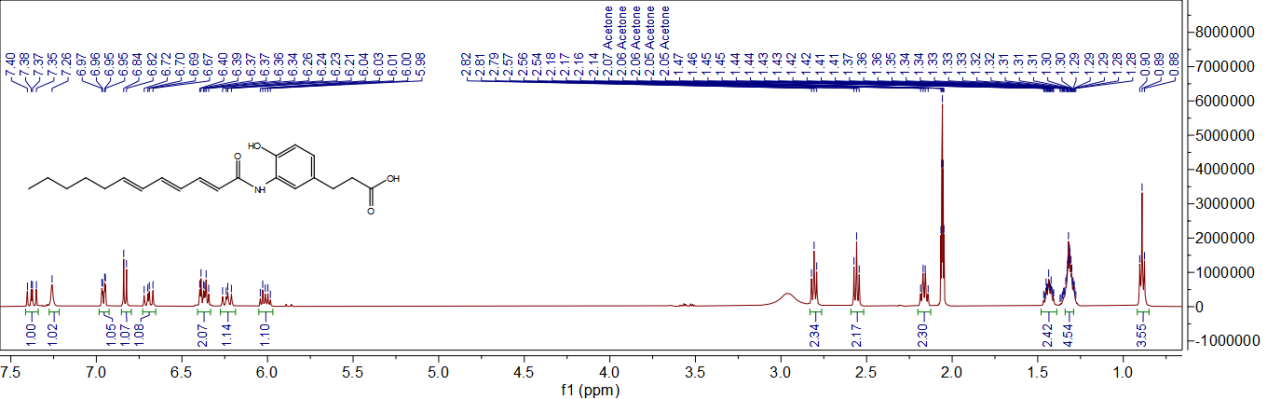

**Fig. 19 | ^1^H (500 MHz) NMR spectra of compound 6 in acetone-*d_6_*.**

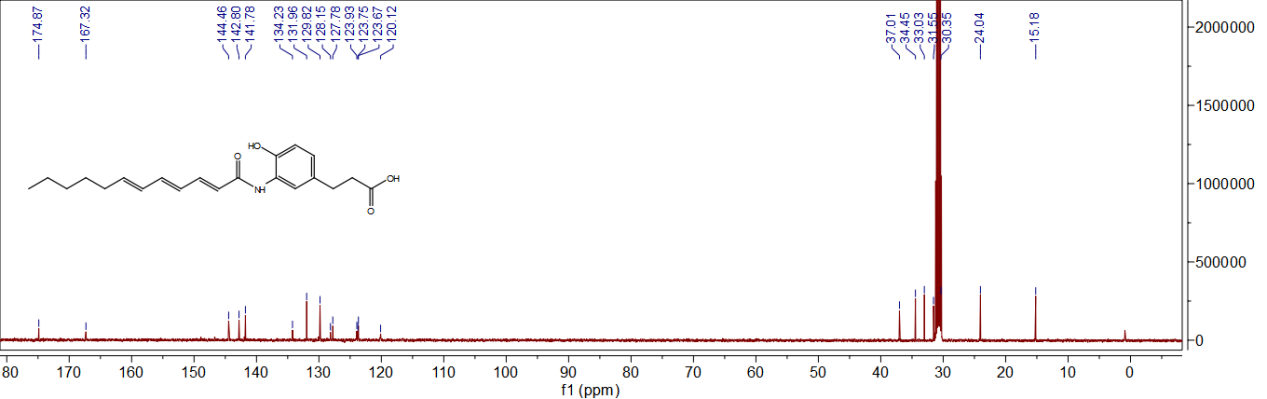

**Fig. 20 | ^13^C (126 MHz) NMR spectra of compound 6 in acetone-*d_6_*.**

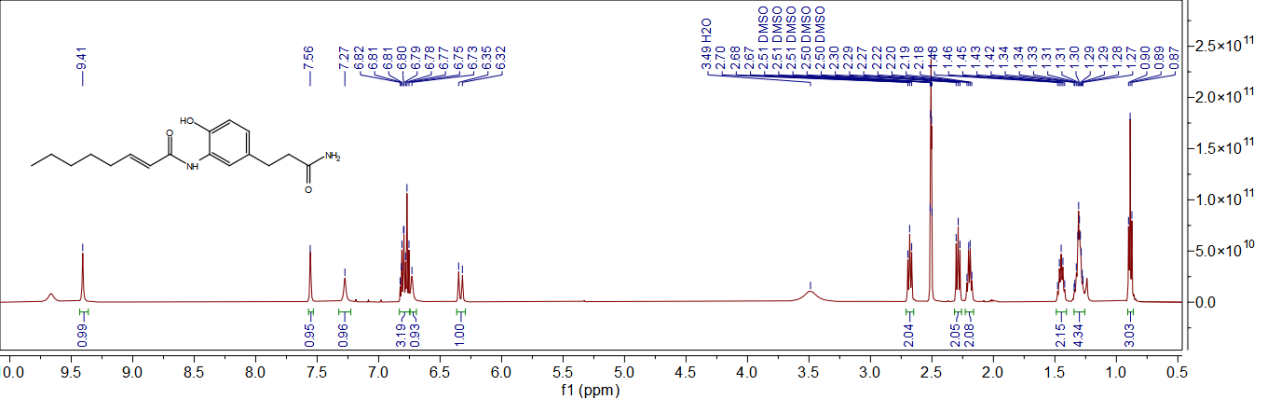

**Fig. 21 | ^1^H (500 MHz) NMR spectra of compound 7 in acetone-*d_6_*.**

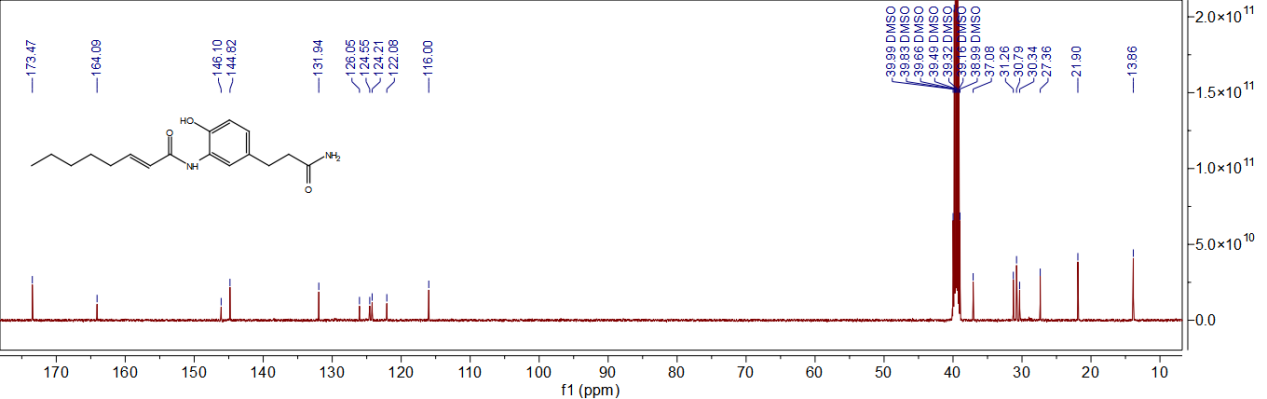

**Fig. 22 | ^13^C (126 MHz) NMR spectra of compound 7 in acetone-*d_6_*.**

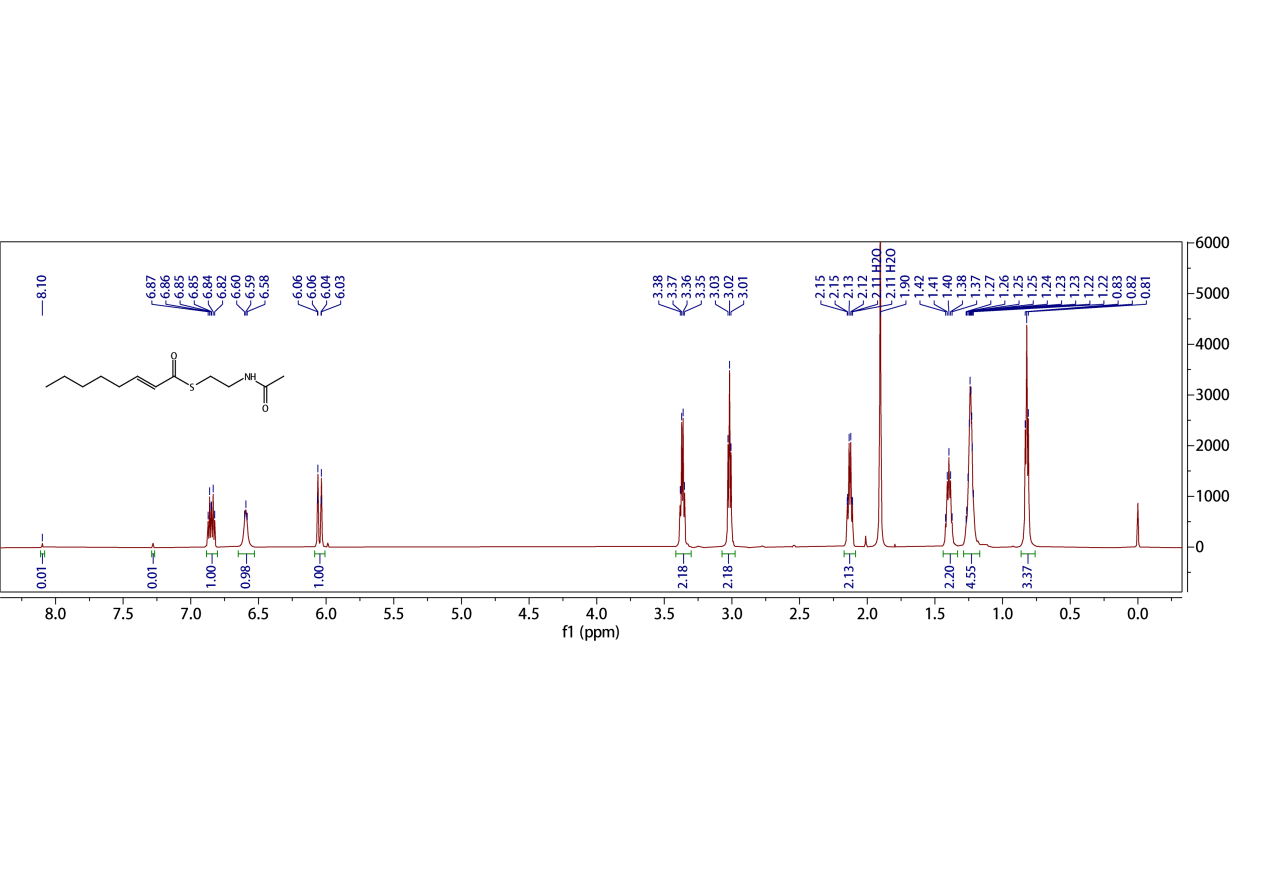

**Fig. 23 | ^1^H (600M) NMR spectra of compound 10 in CDCl_3_.**

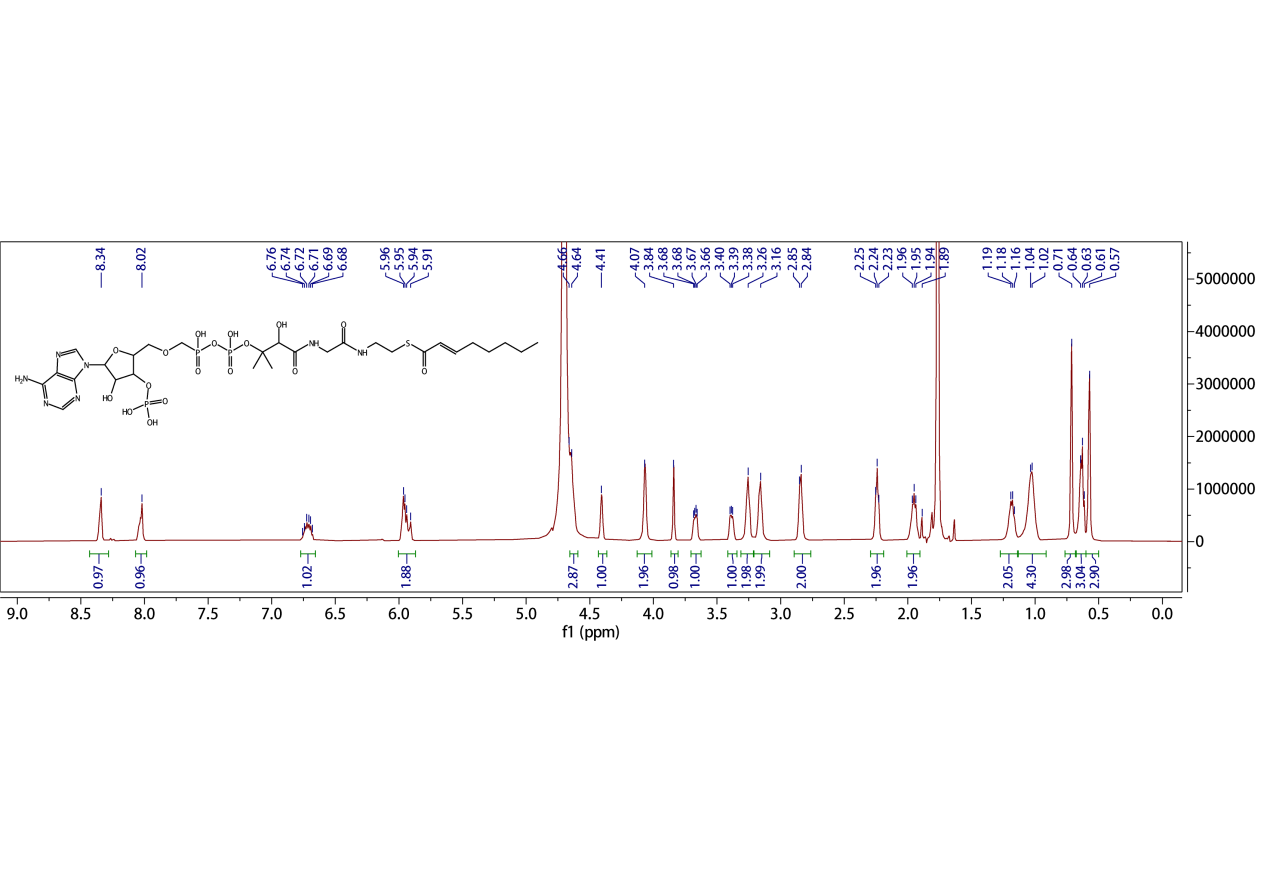

**Fig. 24 | ^1^H (500M) NMR spectra of compound 12 in D_2_O.**

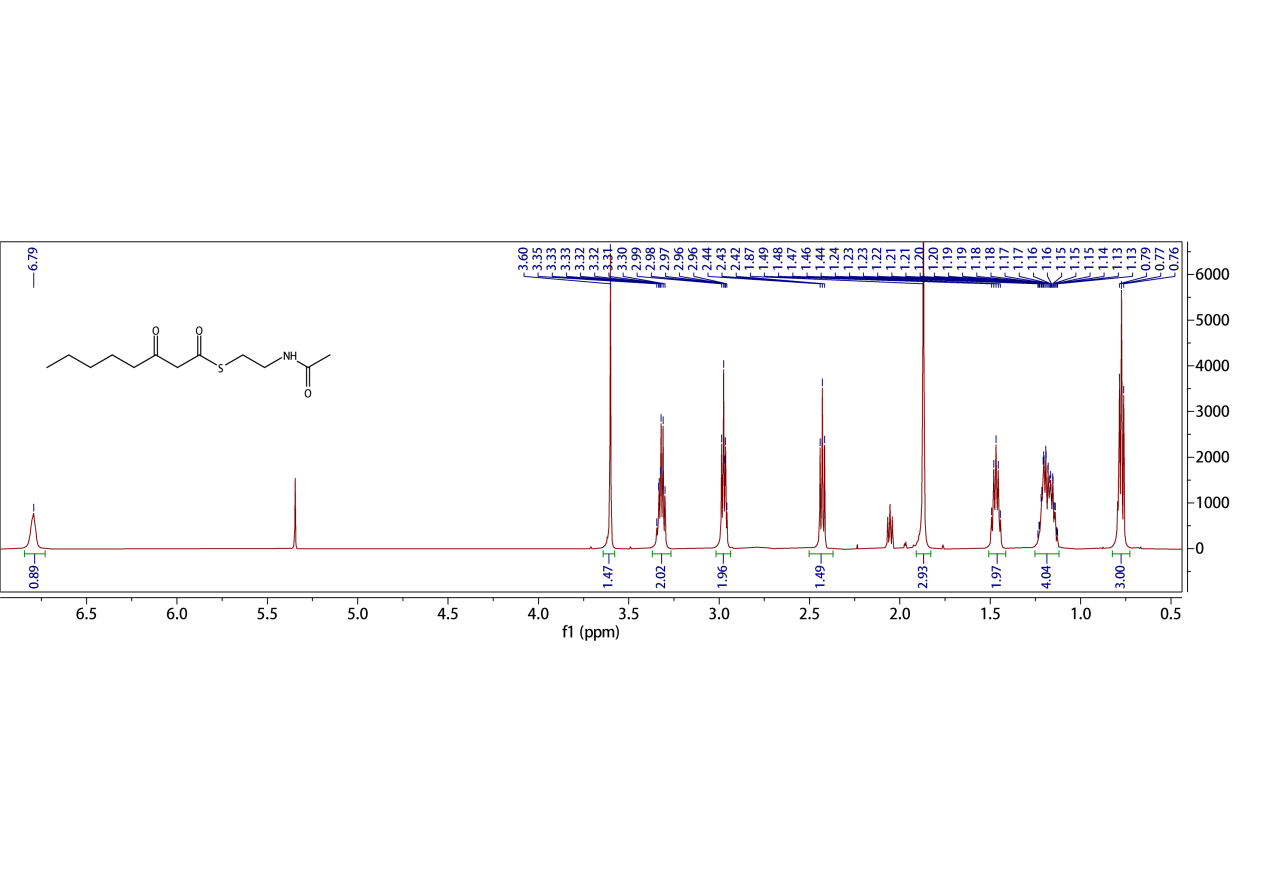

**Fig. 25 | ^1^H (600M) NMR spectra of compound 13 in CDCl_3_.**

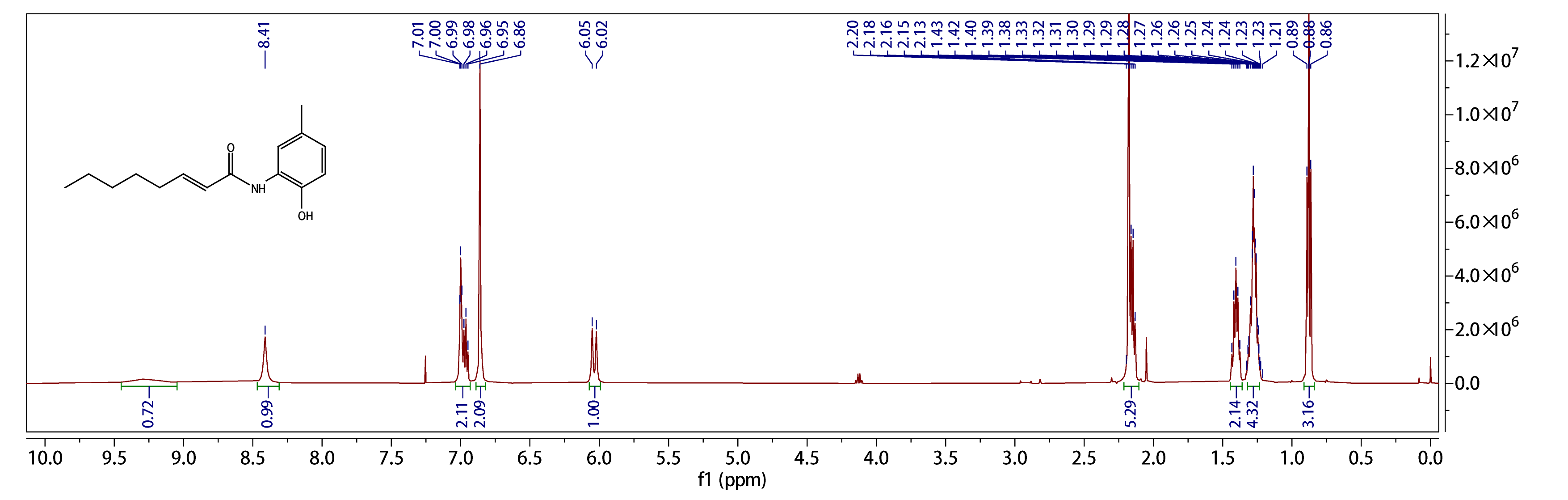

**Fig. 26 | ^1^H NMR (500 MHz) spectra of compound 29 in CDCl_3._**

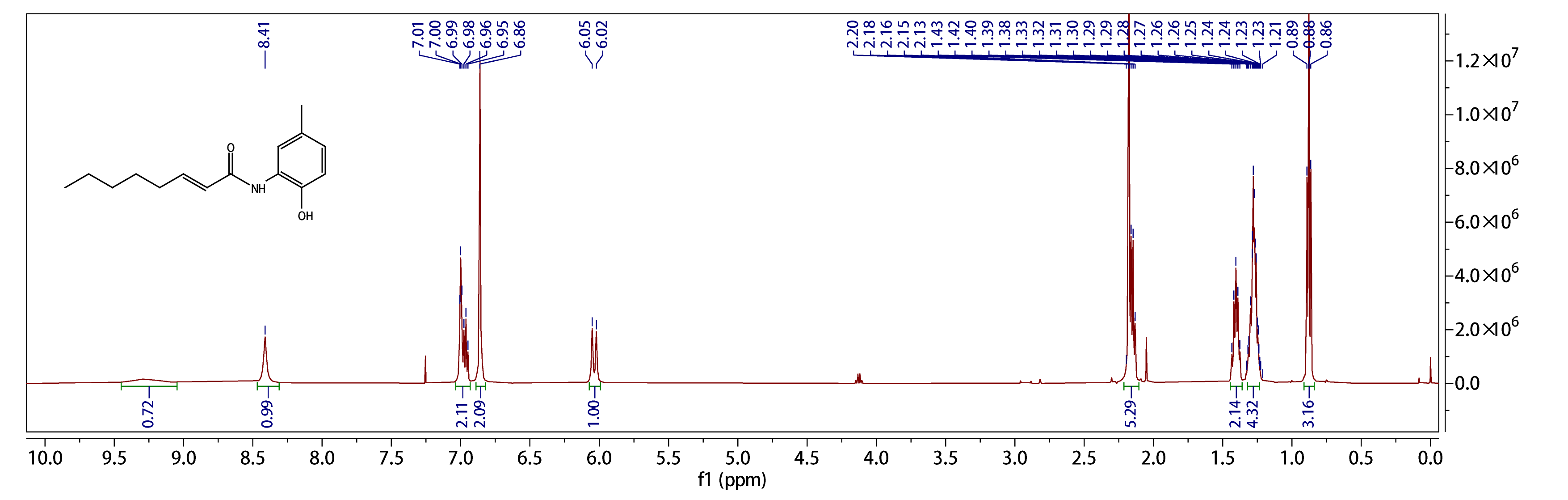

**Fig. 27 | ^13^C NMR (126 MHz) spectra of compound 29 in CDCl_3._**

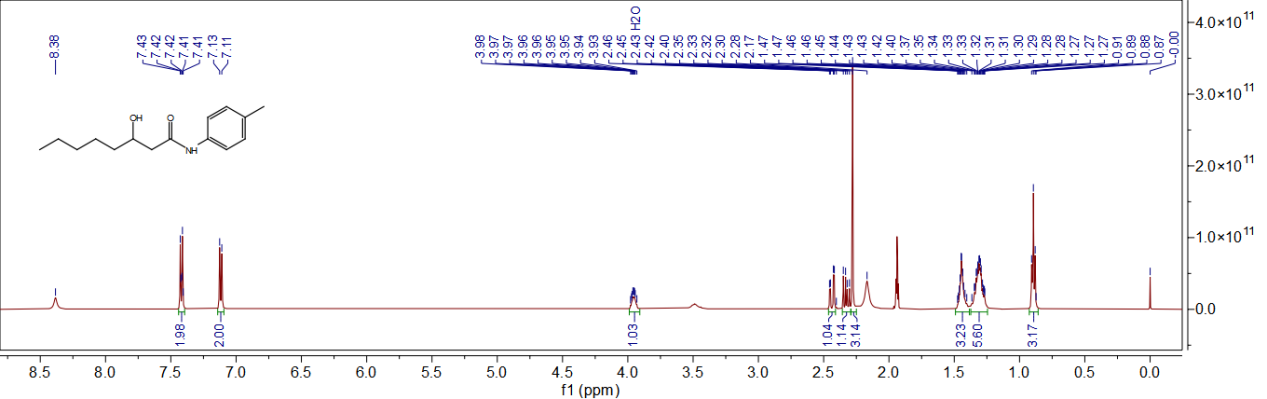

**Fig. 28 | ^1^H (500 MHz) NMR spectra of compound 40 in CD_3_CN.**

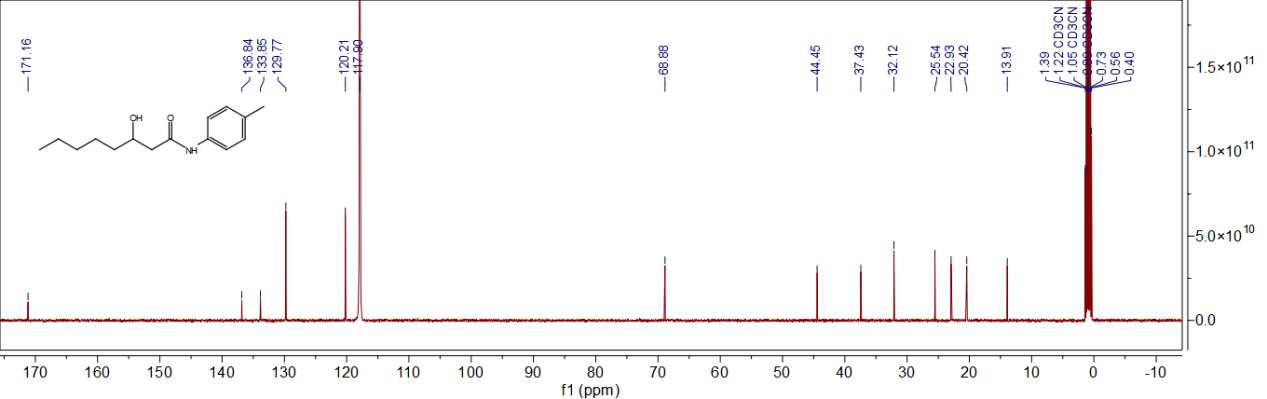

**Fig. 29 | ^13^C (126 MHz) NMR spectra of compound 40 in CD_3_CN.**

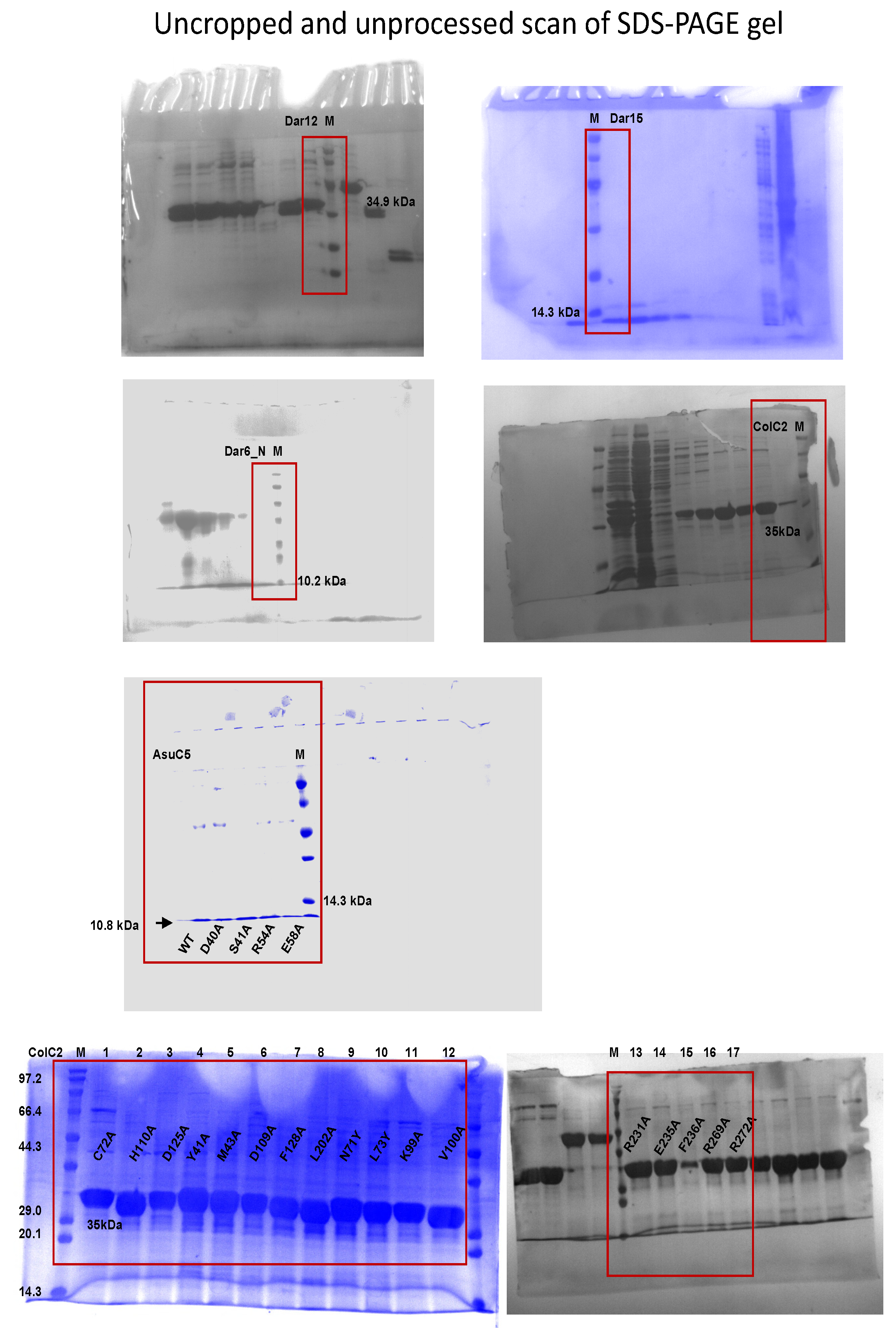
